# Integrated electrophysiological and genomic profiles of single cells reveal spiking tumor cells in human glioma

**DOI:** 10.1101/2024.03.02.583026

**Authors:** Rachel N. Curry, Qianqian Ma, Malcolm F. McDonald, Yeunjung Ko, Snigdha Srivastava, Pey-Shyuan Chin, Peihao He, Brittney Lozzi, Prazwal Athukuri, Junzhan Jing, Su Wang, Arif O. Harmanci, Benjamin Arenkiel, Xiaolong Jiang, Benjamin Deneen, Ganesh Rao, Akdes Serin Harmanci

**Author notes:** These authors contributed equally to this work (Rachel N. Curry, Qianqian Ma). These authors contributed equally to this work (Xiaolong Jiang, Benjamin Deneen, Ganesh Rao, Akdes Serin Harmanci).

## Abstract

Prior studies have described the complex interplay that exists between glioma cells and neurons, however, the electrophysiological properties endogenous to tumor cells remain obscure. To address this, we employed Patch-sequencing on human glioma specimens and found that one third of patched cells in *IDH* mutant (IDH^mut^) tumors demonstrate properties of both neurons and glia by firing single, short action potentials. To define these hybrid cells (HCs) and discern if they are tumor in origin, we developed a computational tool, Single Cell Rule Association Mining (SCRAM), to annotate each cell individually. SCRAM revealed that HCs represent tumor and non-tumor cells that feature GABAergic neuron and oligodendrocyte precursor cell signatures. These studies are the first to characterize the combined electrophysiological and molecular properties of human glioma cells and describe a new cell type in human glioma with unique electrophysiological and transcriptomic properties that are likely also present in the non-tumor mammalian brain.

## Introduction

Glioma is the most common central nervous system tumor with an estimated 20,000 cases diagnosed each year^1^. These diffuse glial tumors include isocitrate dehydrogenase (IDH) mutant (IDH^mut^) and *IDH* wildtype (IDH^WT^) subtypes, each of which presents with unique clinical and histopathological correlates. Prognostic outcomes for IDH^WT^ tumors are poor, conferring a median survival of less than 14 months^2^. In contrast, IDH^mut^ tumors confer significantly better prognoses, with a median survival of 31-65 months after diagnosis^3^. While IDH^WT^ tumors typically result from driver mutations in tumor suppressors or oncogenes^4^, IDH^mut^ tumors uniformly feature mutations in *IDH1* or *IDH2*. Robust molecular and genomic studies conducted over the last two decades have revealed that the disparity in survival outcomes between glioma subtypes is primarily attributed to differences in tumor cell proliferation and invasiveness^5^, which are mediated by an intricate compendium of tumor intrinsic and extrinsic factors. Among these, communication between tumor cells and their microenvironmental constituents has proven to be a critical mediator of glioma progression and is largely conducted by immunological and neural cellular components^6,7^. With regards to the latter, recent advances in the field of cancer neuroscience have revealed that glioma cells form functional synapses with peritumoral neurons and that excitatory neuronal activity promotes glioma progression via increased proliferation and infiltration^7–10^. Conversely, reports have suggested that inhibitory neuronal activity mediated by γ-aminobutyric acid (GABA) slows glioma growth^11^, however emerging studies have demonstrated pro-tumorigenic effects of GABAergic signaling as well^12^. While tumor-neuron interactions have garnered significant attention, the electrophysiological profiles of tumor cells as they exist *in situ* within the human brain remain poorly defined.

Over the past decade, technological advances in single cell genomics have generated an abundance of sequencing data, elucidating the robust transcriptional and genomic heterogeneity that exists in glioma^13–15^. Moreover, Patch-sequencing (Patch-seq), which integrates whole cell recordings, morphological analysis and single cell RNA-sequencing (scRNA-seq), permits characterization of both electrophysiological and transcriptomic features in individual cells^16^ and has been used to expound the extraordinary array of neuronal subtypes present in the mammalian brain^17^. While these technical advances have generated ample substrate in the form of sequencing data, the challenge of accurately identifying cell types from these data remains cumbersome. Specifically, a streamlined computational framework capable of annotating glioma cells has yet to be developed and cell annotation algorithms remain ill-equipped to assign integrated genomic and transcriptional profiles to single cells on a cell-by-cell basis. Because glioma cells frequently share molecular profiles with their non-tumor glial analogs, development of reliable methodologies for identifying tumor cells within the brain poses a difficult but necessary task.

To address the aforementioned computational limitations and improve our biological understanding of tumor cell electrophysiology, we performed *in situ* Patch-seq studies on surgically-resected human glioma samples and developed a new single cell computational tool, Single Cell Rule Association Mining (SCRAM), to characterize the genomic and transcriptomic features of recorded cells. Collectively, our studies demonstrate that a subset of human glioma cells fire single, short action potentials (APs) and are defined by an amalgamation of GABAergic neuron and oligodendrocyte precursor cell (OPC) transcriptomes, which we term GABA-OPCs.

## Results

### Hybrid cells fire single action potentials and are distinct from neurons and glia

To determine the electrophysiological properties of tumor cells in malignant glioma, we performed whole-cell patch clamp recordings followed by scRNA-seq (Patch-seq) on brain slices surgically-resected from nine patients, including six IDH^mut^ gliomas, two IDH^WT^ gliomas and one non-tumor sample (**Fig. 1a, Table S1-2**). We recorded from a total of 148 cells; 95 were used to recover high-quality RNA for scRNA-seq, and 44 were preserved with biocytin for morphological analysis. For the 95 cells used for sequencing, an average of 4491 genes were identified per cell (**Table S3**). Of the 148 cells, 105 showed electrophysiological and morphological profiles consistent with established neural cell types in the mammalian brain and could be broadly classified as pyramidal cells (PCs; n=54), inhibitory neurons (INs; n=24) or glia (GL; n=27) based on maximal AP firing rates, AP amplitudes, input resistances and morphology^18,19^ (**Figs. 1b-d; Fig. S1-3; Tables S2**). It should be noted that we are using the electrophysiological annotation of GL to generally describe electrically inert cells, which may also include immature neurons and neural precursor cells (NPCs). Intriguingly, 43 cells across four IDH^mut^ glioma and one non-tumor sample displayed select neuronal electrophysiological properties but were morphologically inconsistent with mature neuronal cell types, frequently resembling GL or NPCs (**Figs. 1e-g; Fig. S1-3; Tables S2**). These cells, which we assigned as hybrid cells (HCs), electrophysiologically represented 34% of IDH^mut^ and 30% of non-tumor patched cells (**Fig. 1h**), had higher input resistances than neurons or GL, and were uniformly capable of firing single, small APs (**Figs. 1i-k; Fig. S1-3, Fig. S4-7**). Here, we define AP by a minimum dv/dt of 20V/S, minimum peak height of 2mV, a minimum absolute peak level of −20mV, a maximum interval of 10mS, and thresh fraction of 0.05. In contrast, none of the 20 cells recorded from two IDH^WT^ tumors demonstrated HC profiles and no HCs or GL (**Fig. S1h**).

**Figure 1.**
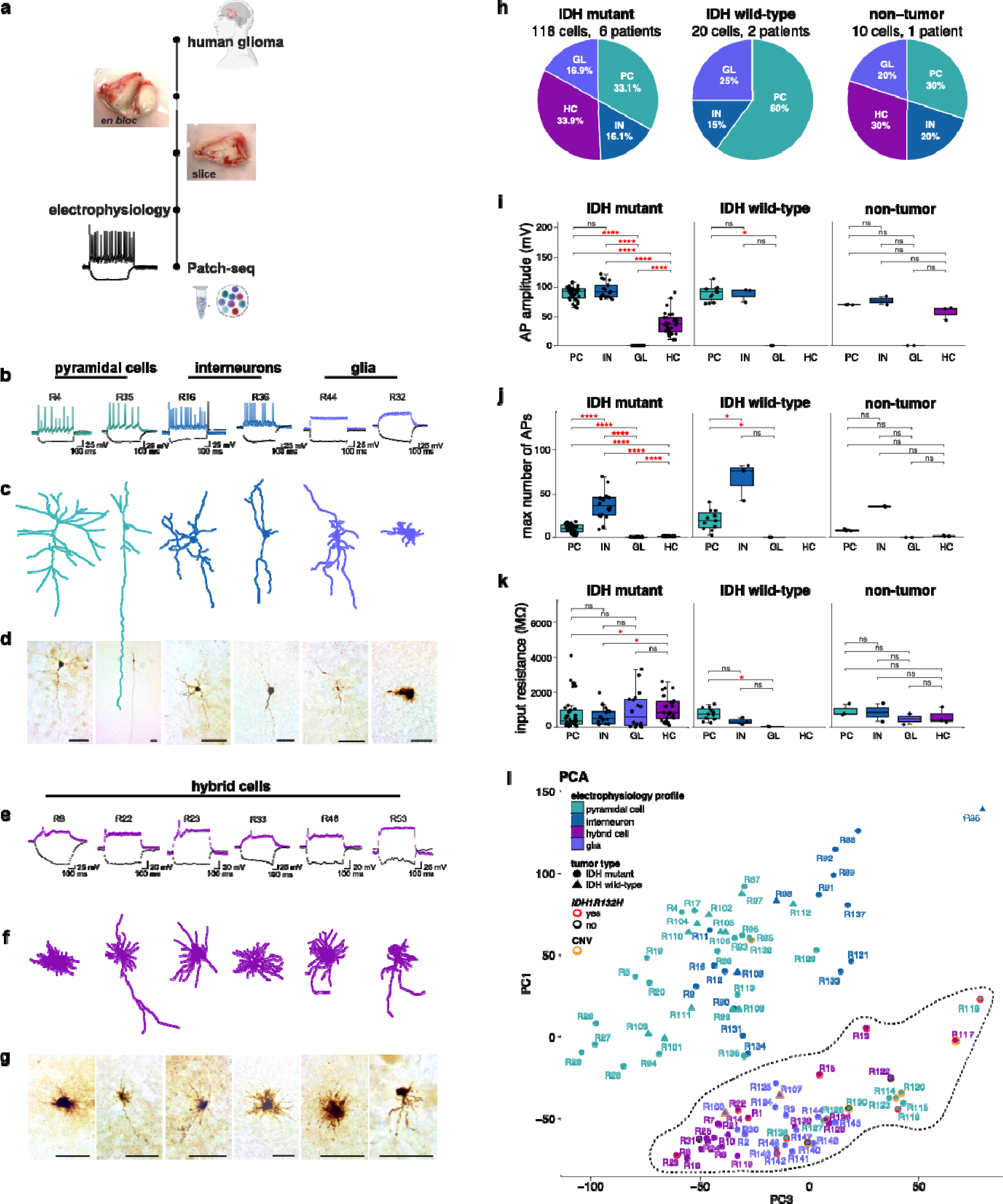
Patch-seq of human glioma samples reveals tumor cells fire action potentials. (a) Experimental workflow for whole-cell recordings and whole-cell patch clamp recordings, followed by singe-cell RNA-sequencing (Patch-seq) assays. (b) Exemplary membrane responses from patched PCs, INs and GL to a 600-ms hyperpolarizing current step (black) and suprathreshold depolarizing current step (colored). (c) Matched traced cell morphologies are shown for recorded neuronal and glial cells. (d) Matched images of biocytin-filled cell morphologies for patched neuronal and glial cells; scale bar: 50 µm. (e) Exemplary membrane responses from IDH^mut^ HCs to a 600-ms hyperpolarizing current step (black) and suprathreshold depolarizing current step (colored). (f) Matched traced cell morphologies are shown for recorded HCs. (g) Matched images of biocytin-filled cell morphologies for patched HCs; scale bar: 50 µm. (h) Pie chart showing percentage of PC, IN, GL and HC patched by experimental group. (i) Box plot showing HC cells have AP amplitudes compared to neurons. (j) Box plot showing HCs fire fewer spikes compared to neurons. (k) Box plot showing HCs have higher input resistance compared with non-tumor neurons. (l) Principal component analysis (PCA) plot of 95 Patch-seq cells shows clustering of cells based on electrophysiological properties. HCs are transcriptionally similar to each other, GL and select PCs. Cells are colored according to electrophysiological properties, *IDH1R132H* status and CNV status. Black dashed line denotes PCA cluster. For Patch-seq, two voltage traces are shown: the hyperpolarization trace obtained with injected currents (black) and the depolarization trace showing maximal AP firing rate; injected current: −100 pA. *p*-values for pairwise comparisons are noted in the figure. AP: action potential; GABA: γ-aminobutyric acid; GL: glia; GSC: glioma stem cell; IN: interneuron; OL: oligodendrocyte; OPC: oligodendrocyte precursor cell; PC: pyramidal cell.

To investigate the molecular profiles of Patch-seq cells, we first performed principal component analysis (PCA) and found that all HCs clustered together and with GL (**Fig. 1l**). Notably, a group of ten PCs (hereto referred to as ΔPCs) coming from one recurrent IDH^mut^ patient that were embedded within this PCA cluster (**Fig. 1l, black dashed line**) also featured abnormally high input resistances, like those of HCs, and had lower maximal AP firing rates than other recorded PCs (**Fig. S8**). We hypothesized that the HCs, GL and ΔPCs within this PCA cluster represented tumor cells, thus we sought to confirm this using RNA-inferred single nucleotide variant (SNV) and copy number variant (CNV) analysis. We looked for the canonical *IDHR132H* mutation, chromosome 1p and 19q deletions, and chromosome 7p amplifications, which are established genomic markers of glioma and found that in IDH^mut^ patients, seven HCs, six ΔPCs and two GL were tumor cells, and that two GL were tumor cells in IDH^WT^ samples (**Fig. 1m; Table S2**, **Fig. S9**). Two HCs from IDH^mut^ samples were euploid, had coverage of the *IDH1R132* locus and were not mutated, and three HCs were detected in the non-tumor sample, confirming that HC electrophysiology is not exclusive to tumor cells (**Fig. 1l, Fig. S2**). These initial studies are the first to show that *bona fide* glioma cells are neurophysiologically diverse and can present with inert, single-spiking or excitatory electrophysiology profiles.

### SCRAM is a reliable annotation tool for human scRNA-seq datasets

To better define the HCs identified in our Patch-seq studies, we sought to create a new computational platform that could annotate each cell from Patch-seq individually. Because of the low cell numbers obtained using Patch-seq and the rarity of human glioma samples for use in these experiments, our annotation tool needed to be capable of analyzing each cell without a dependency on clustering methodologies, which requires hundreds to thousands of cells for optimal analysis. Accordingly, we developed the Single Cell Rule Association Mining (SCRAM) tool that can annotate each cell on a cell-by-cell basis, independently of clusters. SCRAM uses a three-step orthogonal process to provide detailed transcriptional and RNA-inferred genomic profiles for each cell: (1) cell type transcriptional annotation using machine-learned neural network models (NNMs); (2) single nucleotide variant (SNV) profiling using the XCVATR^20^ tool; and (3) copy number variant (CNV) calling using the CaSpER^21^ and NUMBAT^22^ tools (**Fig. 2a-d**). SCRAM NNMs are trained on 11 previously published scRNA-seq (totaling ∼1M cells) human tumor and non-tumor datasets, including developing brain, immune, and glioma cell atlases^14,15,23–30^ **(Table S4-5)**. Each cell from Patch-seq is assigned a probability score for each cell type in the training datasets **(Fig. 2e)**. Using this methodology, individual cells may be annotated as more than one cell type, which permits for the characterization of hybrid cellular states like those of HCs from our Patch-seq experiments. SNVs and CNVs are also considered for each cell and added to the cell annotation. Cells are assigned as “tumor” if they have ≥2 tumor features (see *Methods*). We validated that SCRAM can reliably discriminate between tumor and non-tumor tissue, which it does with >99% sensitivity (true positive rate: TPR) in Allen Brain Atlas (ABA)^23^, Bhaduri et al.^25^, Aldinger et al. ^29^, CoDEx^26^ and >96% sensitivity in Human Protein Atlas (HPA)-Brain^30^ (**Table S6**). Additionally, we ran SCRAM on glioma scRNA-seq datasets^13,15,24^, in which tumor and non-tumor annotations are reported in cluster resolution, finding that SCRAM achieved 100% specificity (true negative rate: TNR) for tumor cells in IDH^mut^ and IDH^WT^ glioma datasets.

**Figure 2.**
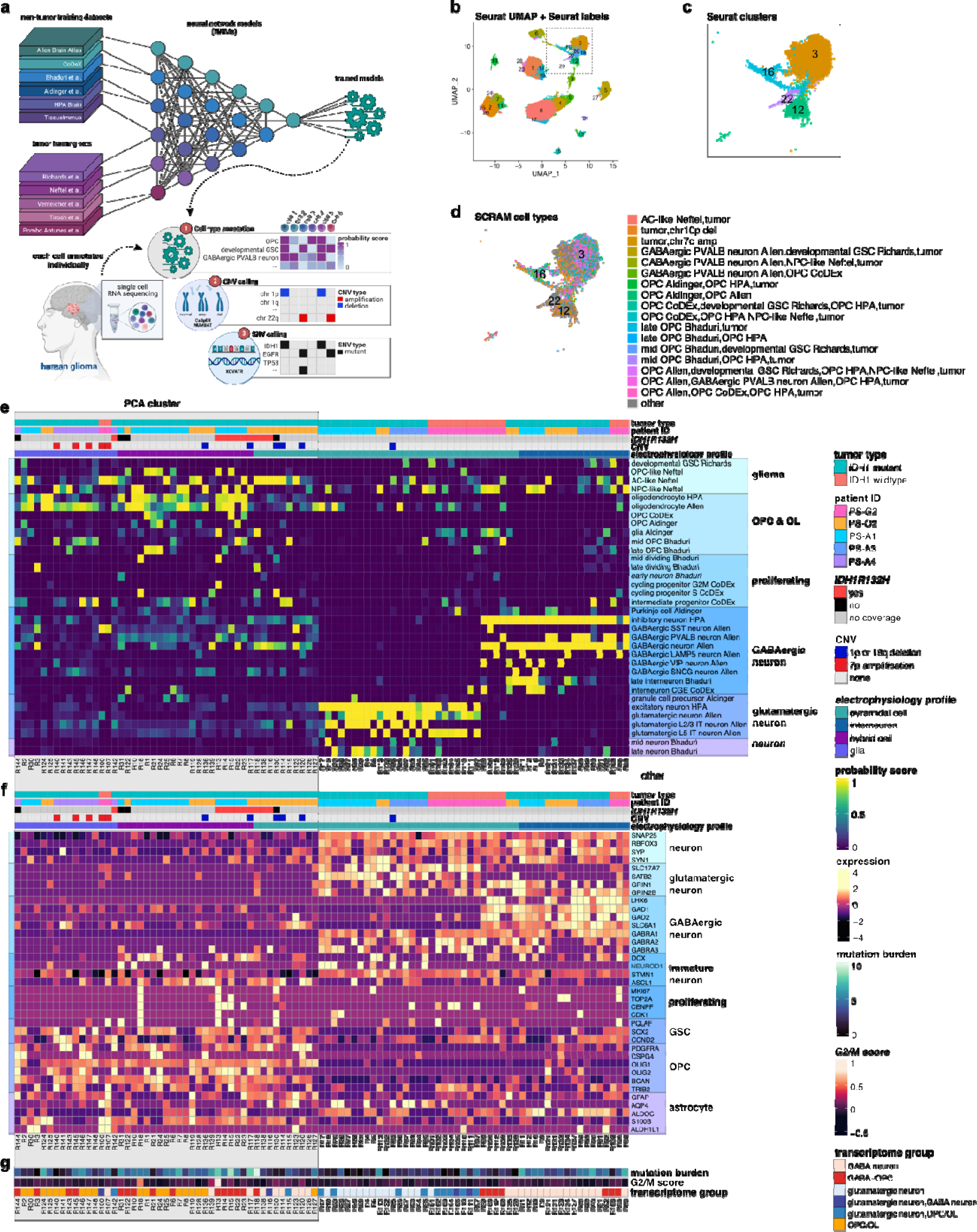
SCRAM reveals HC cells are GABA-OPC tumor cells. (a) Schematic of the SCRAM pipeline. Briefly, 11 scRNA-seq datasets were used to train cell type neural network models (NNMs). Each cell from scRNA-seq is then assigned cell type annotation independently of all other cells using NNM trained models. CNVs are added for each cell using CaSpER and NUMBAT. SNVs are added for each cell using XCAVTR. (b) Seurat clusters are shown for 234,880 cells from our in-house glioma scRNA-seq dataset. (c) Zoom-in of black dashed box from *(b)* Seurat clusters 3, 12, 16 and 22 colored by Seurat clusters. (d) Zoom-in of black dashed box from *(b)* Seurat clusters 3, 12, 16 and 22 colored by SCRAM cell-by-cell annotations. (e) Heatmap showing SCRAM cell type probability scores for 95 Patch-seq cells. (f) Heatmap showing cell-type markers for 95 Patch-seq cells. (g) Heatmap showing composite SNV scores (mutational burden) and cell cycle score (G2/M score) for 95 Patch-seq cells. Grey box denotes cells marked in PCA cluster from Figure 1l. CNV: copy number variant; SNV: single nucleotide variant.

### HCs express hybrid GABAergic neuron-OPC transcriptomes

We ran SCRAM on the 95 cells from Patch-seq experiments and found that all cells with IN electrophysiology (n=17) were correctly annotated as GABAergic neurons, and that 24 out of 41 cells with PC electrophysiology were appropriately annotated as glutamatergic neurons (**Fig. 2e**). All cells with GL electrophysiology (n=16) were annotated as glial subtypes and/or glioma cells. Thirteen PCs identified from one IDH^WT^ patient and one recurrent IDH^mut^ patient were annotated as GABAergic neurons. These cells featured maximal AP firing rates consistent with excitatory neurons despite robust GABAergic neuron transcriptomic profiles (**Figs. 2e-g**), a dichotomy reminiscent of the neurodevelopmental paradigm in which GABA neurotransmission confers excitatory signaling^31^. Using cell cycle scoring, we found that only three cells showed proliferative signatures and that two of these were *IDH1R132H* mutant (**Fig. 2g**); however, the majority of HCs showed low G2/M scores. Expectedly, mutation burden was highest in HCs and GL and was specifically enriched in HCs bearing the *IDH1R132H* mutation. Consistent with their electrophysiological profiles, all HCs (n=21) showed concurrent annotation as OPCs, oligodendrocytes (OLs) and GABAergic neurons (an amalgam hereto referred to as GABA-OPC) and tumor cells, signifying that HCs are endowed with a mixture of neuronal and oligodendroglial transcriptional features that may be responsible for the functional electrophysiological properties of these cells (**Figs. 2e-g; Fig. S10**).

To validate our Patch-seq findings in larger datasets, we used SCRAM to analyze our in-house scRNA-seq dataset consisting of 234,880 cells from 12 IDH^mut^ and IDH^WT^ glioma patients^32^ (**Fig. 2b-d, Fig. S11; Table S7**). SCRAM-assigned probability scores >0.9 were used for final cell annotations and to generate a SCRAM UMAP of our scRNA-seq glioma dataset, which clusters cells based on cell identity rather than similarity of transcriptional features (**Fig. 3a, Figs. S12-21**). Visualization of SNVs, CNVs and tumor expression markers specific to glioma subtypes revealed that SCRAM segregates the majority of IDH^mut^ from IDH^WT^ tumor cells (**Fig. 3b**). Given that the HCs we observed in our Patch-seq experiment were found in IDH^mut^ patients, we focused on SCRAM clusters encompassing the majority of IDH^mut^ tumor cells, which were predominantly distributed amongst three discrete SCRAM clusters (Clusters 13, 18, and 19) that collectively contained <5% of IDH^WT^ tumor cells (**Figs. 3c-e**). Importantly, all IDH^mut^ tumor patients had robust GABA-OPC expression profiles, whereas only one IDH^WT^ patient showed the same expression profile (**Figs. 3f-g, Figs. S22-24**). An analysis of tumor cells by patient revealed that on average 41.3% of IDH^mut^ tumor cells were GABA-OPCs (**Fig. 3h**), which was consistent with the presence of HC electrophysiology in 34% of recorded cells from these tumors. In contrast, only 2.2% of tumor cells received GABA-OPC annotations in IDH^WT^ tumors (**Fig. 3h**), which was consistent with the absence of HC electrophysiologies from Patch-seq experiments.

**Figure 3.**
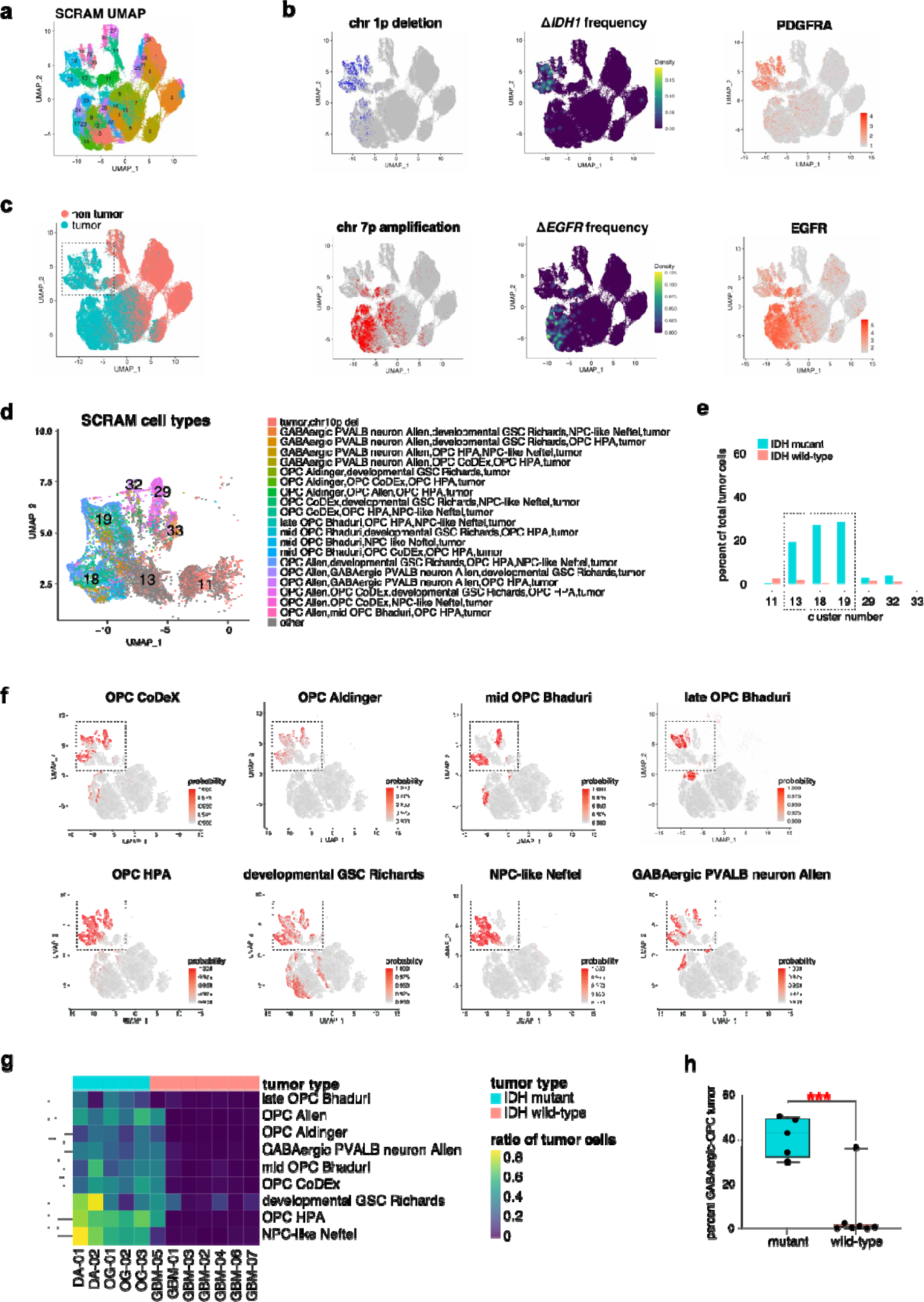
GABA-OPC tumor cells in human glioma. (a) SCRAM UMAP of 234,880 scRNA-seq cells. (b) Commonly used IDH^mut^ (top row) and IDH^WT^ (bottom row) tumor features are shown. Top (from left to right): chromosome 1p deletion feature plot, *IDH1* mutation density plot, *PDGFRA* expression feature plot. Bottom (from left to right): chromosome 7p amplification feature plot, *EGFR* mutation density plot, *EGFR* expression feature plot. (c) SCRAM tumor and non-tumor cell annotation. (d) Zoom-in of inset from *(c)* showing SCRAM cell type annotations for each cell. (e) Bar graph showing the majority of cells in *(d)* are from IDH^mut^ tumor patients. (f) SCRAM probability scores are shown for cell types of interest. (g) Heatmap of SCRAM cell type annotations for IDH^mut^ (n=5) and IDH^WT^ (n=7) glioma patients. (h) Bar graph showing the percentage of tumor cells with GABA-OPC annotations; *p*-value is noted in the figure.

### Feature extraction of GABA-OPCs reveals GABAergic neuron and OPC transcriptional dependencies

Having identified that HCs are defined by GABA-OPC transcriptomes and that these cells constitute a large proportion of tumor cells in IDH^mut^ patients, we sought to extract a GABA-OPC molecular signature by identifying transcriptional markers. To do this, we used SHapley Additive exPlanations (SHAP) analysis, which extracts essential features from machine-learning NNMs. We identified SHAP markers from each cell type in our training dataset (e.g. GABAergic PVALB neuron from ABA; **Figs. S25-30**) and compared these genes to the differentially expressed genes (DEGs) obtained for GABA-OPC tumor cells as compared to other tumor cells in our scRNA-seq dataset (**Fig. S31-32, Table S8)**. The intersection of SHAP genes and DEGs produced a list of 61 genes (log_2_FC > 1), which we present as the GABA-OPC gene set. The GABA-OPC gene set is comprised of SHAP genes that are critical transcriptional features of cell types from tumor and non-tumor training datasets (**Figs. 4a-b, Table S8**). The training cell types that were most highly represented by GABA-OPC SHAP features were GABAergic neuron and OPC, demonstrating that the majority of GABA-OPC transcriptional characteristics derive from GABAergic neurons and OPCs. Given that OPCs and OLs represent a complex and transcriptionally heterogenous spectrum of cells, we studied OPC and OL cell state signatures from three different studies to better understand which specific OPC and OL annotations were enriched in GABA-OPCs^33,34^. We found that GABA-OPC cells are transcriptionally most like late-stage OPCs and early differentiated OLs, which implies that GABA-OPCs exist in a transitional state between precursor OPC and fully differentiated OL lineages (**Fig. S33-34**). Further bioinformatics analyses determined that on average 48% percent of OPCs from non-tumor and developmental brain atlases possess GABA-OPC molecular profiles and that these cell profiles are more frequent in the adult brain than they are in neurodevelopmental contexts (**Fig. S35)**. These results suggest that GABA-OPC tumor cells are malignant manifestations of a GABA-enriched OPC subclass that is normally found in non-tumor human brain.

**Figure 4.**
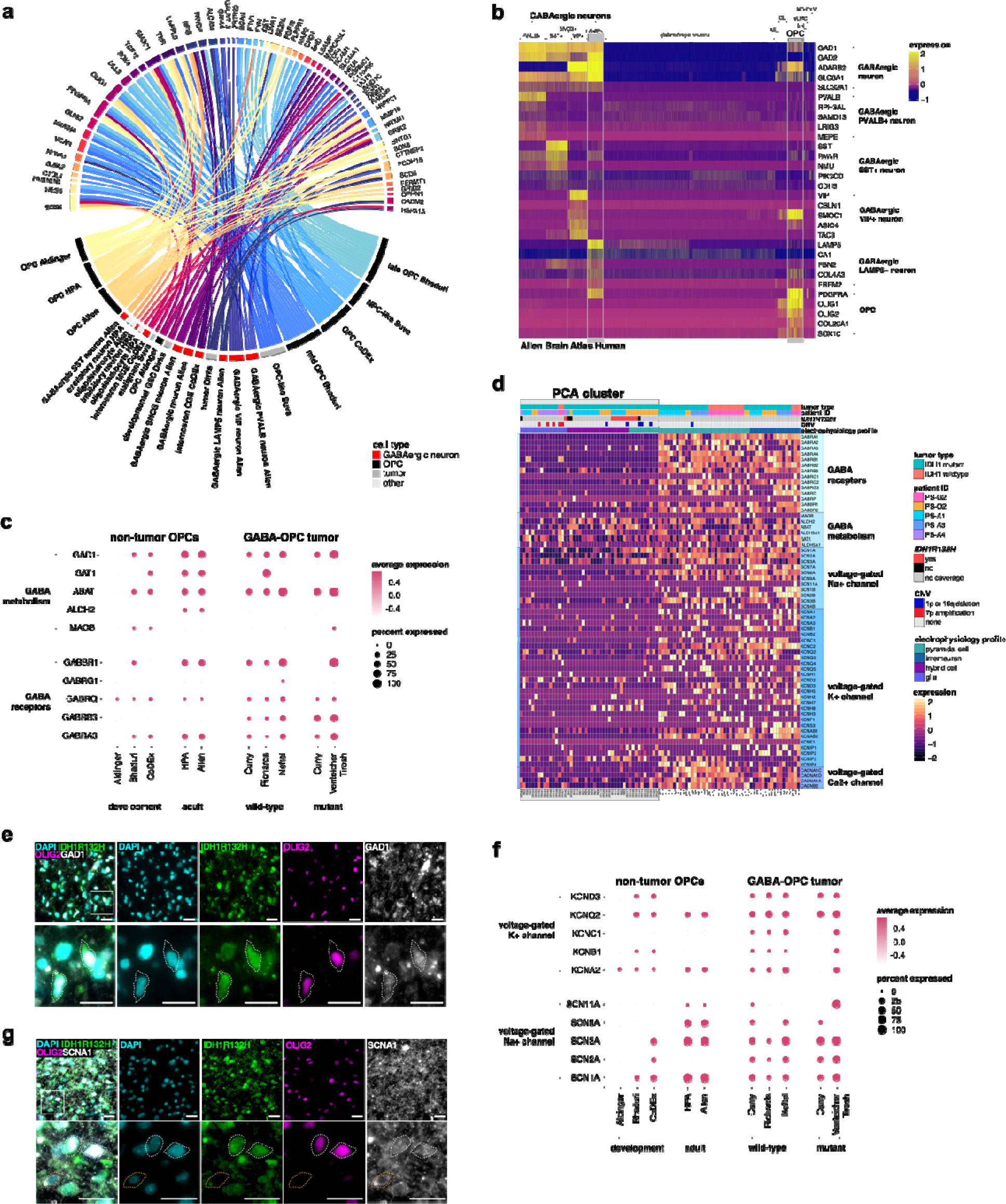
GABAergic, neuronal and OPC features are hallmarks of GABA-OPCs. (a) Circos plot showing the 61 genes (log_2_FC > 1) comprising the GABA-OPC tumor cell signature. DEGs from GABA-OPC tumor cells versus all other tumor cells were extracted from our scRNA-seq dataset and crossmatched with SHAP genes from our trained NNMs. (b) Heatmap of Allen Brain Atlas non-tumor human scRNA-seq data showing GABAergic neurons and OPCs share subtype-specific markers. Grey boxes outline LAMP5+ GABAergic neurons and OPCs. (c) DotPlot showing the average expression of GABARs and GABA metabolism genes in human non-tumor OPCs and in GABA-OPC tumor cells. (d) Heatmap showing the expression of GABARs, GABA metabolism genes, voltage-gated sodium channels (Na_v_s), voltage-gated potassium channels (K_v_s) and voltage-gated calcium channels (Ca^2+^) in Patch-seq data. (e) Immunostaining for GAD1 (white), IDHR132H (green) and OLIG2 (pink) in a human IDH^mut^ tumor sample; scale bar = 20 μm. White box denotes inset. White dashed lines denote IDH1R132H+OLIG2+GAD1+ tumor cells. (f) DotPlot showing the average expression of voltage-gated sodium channels (Na_v_s) and voltage-gated potassium channels (K_v_s) genes in human non-tumor OPCs and also in GABA-OPC tumor cells. (g) Immunostaining for Na_V_1.1 (SCN1A; white), IDHR132H (green) and OLIG2 (pink) in a human IDH^mut^ tumor sample; scale bar = 20 μm. White box denotes inset. White dashed lines denote IDH1R132H+OLIG2+SCN1A+ tumor cells. NNM: neural network module; SHAP: SHapley Additive exPlanations.

Having extracted a GABA-OPC transcriptional signature, we next sought to elucidate the molecular constituents that transcriptionally confer GABAergic neuronal properties in HCs. Preceding reports have documented the existence of GABARs and GABA synthesis genes in OPCs^35–37^, which prompted us to investigate the expression of these genes in GABA-OPC tumor cells. We found high expression of *glutamate decarboxylase 1 (GAD1)* and *GABA transaminase (ABAT)*, which are crucial for GABA synthesis and metabolism (**Fig. 4c**). Analysis of these genes in our Patch-seq cohort confirmed that *ABAT* and *GAD1* are highly expressed by HCs (**Fig. 4d**), the latter of which we confirmed through immunostaining in human IDH^mut^ glioma (**Fig. 4e**). Select GABARs were also expressed in GABA-OPC tumor cells as compared to other tumor cells, suggesting that GABA-OPC tumor cells are transcriptionally equipped to receive GABA-mediated inputs from neurons (**Figs. 4c-d**). Given this precedent, our analyses suggest that the electrophysiological activity of GABA-OPC tumor cells may be in part conferred through the expression of GABAergic neuronal gene sets.

We next sought to understand how GABA-OPCs are mechanistically producing APs. Importantly, prior investigations have demonstrated that a subgroup of white matter OPCs in the rat brain fire single^38^, short APs that are similar to those observed in HCs from our Patch-seq studies. The AP capacity of these spiking OPCs is dependent on voltage-gated ion channels, particularly voltage-gated sodium channels (Na_v_s) and voltage-gated potassium channels (K_v_s), which we found were selectively expressed by non-tumor OPCs (**Fig. 4f**). We found that most GABA-OPC tumor cells have high expression of Na_v_s but that expression of K_v_s is restricted to a smaller population of cells and is not uniformly present across HC transriptomes (**Fig. 4c**). Immunostaining for the voltage-gated sodium channel Na_v_1.1, encoded by the *SCN1A* gene, in a human IDH^mut^ tumor confirmed GABA-OPC tumor cells express Na_v_s, which are essential for the rising phase of APs^38,39^ (**Fig. 4g**). These data suggest that GABA-OPC tumor cells are endowed with requisite machinery that can mechanistically produce the APs observed in these cells.

### Strong GABA-OPC signatures confer increased survival in IDH^mut^ glioma

To validate our observation that GABA-OPC signatures were more prominent in our IDH^mut^ cohort as compared to IDH^WT^, we used the GABA-OPC-like glioma gene set to score 216 IDH^WT^ and 366 IDH^mut^ bulk RNA-seq samples from TCGA (**Fig. 5a**). Consistent with our internal scRNA-seq dataset, we found that IDH^mut^ samples had higher GABA-OPC scores than IDH^WT^ samples. An analysis of scores by IDH^WT^ molecular subtypes revealed the highest GABA-OPC scores belonged to OPC-like and NPC-like molecular subtypes (**Fig. 5b**). Consistent with these observations, OPC-like samples had significantly higher GABA-OPC tumor percentages than the other molecular subtypes^14^ (**Fig. 5c**). Reexamination of our internal scRNA-seq dataset revealed that the only IDH^WT^ patient in whom a strong GABA-OPC signature was detected was an OPC-like IDH^WT^ subtype, which explains why GABA-OPC tumor cells were enriched in this sample (see **Fig. 3g**). An analysis of the TCGA IDH^mut^ samples by histopathological subtype showed that GABA-OPC scores were higher in low-grade glioma (LGG) than high-grade glioma (HGG), suggesting that stronger GABA-OPC phenotypes are associated with lower grade tumors (**Fig. 5d**). RNAvelocity pseudotime analyses of three IDH^mut^ and one OPC-like IDH^WT^ samples revealed that GABA-OPC tumor cells emerge from more primitive tumor cell types to become the largest population of glioma cells in tumors that bear them, demonstrating that some tumor cells from the divergent genetic backgrounds of IDH^mut^ and IDH^WT^ converge at a shared transcriptional phenotype (**Figs. 5e-f**).

**Figure 5.**
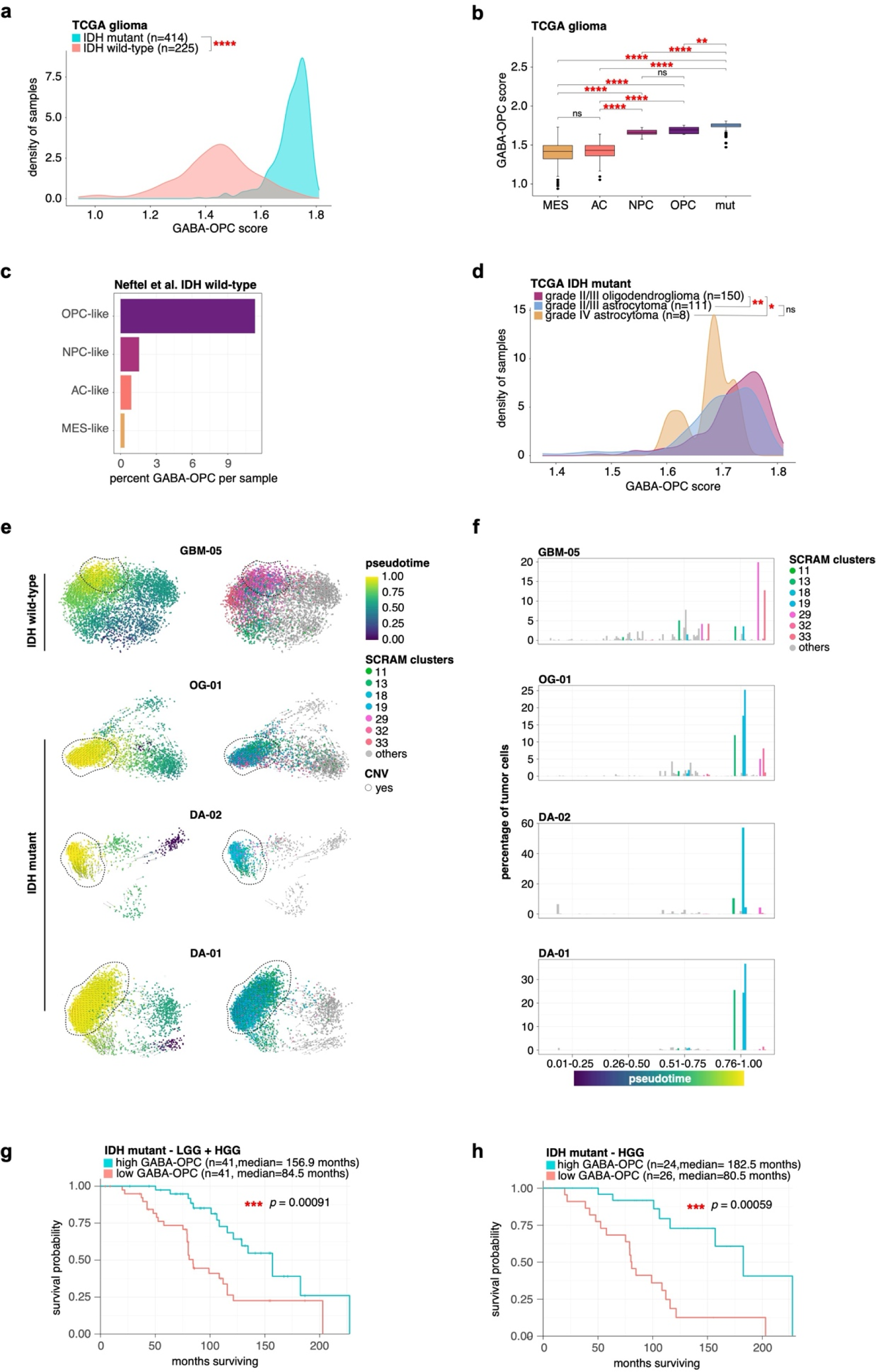
GABA-OPC tumor cells are protective in IDH^mut^ tumors. (a) Density plot of GABA-OPC tumor scores is shown for IDH^mut^ (n=366) and IDH^WT^ (n=216) TCGA bulk RNA-seq glioma samples; *p*-value is noted in the figure. (b) Box and whiskers plot showing GABA-OPC scores for TCGA samples by IDH^WT^ tumor subtype: MES-like (n=149); AC-like (n=157); NPC-like (n=12); OPC-like (n=9); *p*-values for pairwise comparisons are noted in the figure. (c) Bar graph showing the percentage of tumor cells that are GABA-OPC tumor from Neftel et al.’s IDH^WT^ SMART-seq dataset. (d) Density plot of GABA-OPC tumor scores is shown for IDH^mut^ TCGA bulk RNA-seq glioma samples by histology subtype: oligodendroglioma (n=150); astrocytoma (n=111); diffuse astrocytoma (n=8); *p*-values for pairwise comparisons are noted in the figure (e) RNAvelocity pseudotime analyses are shown for one IDH^WT^ and three IDH^mut^ samples. Black dashed lines denote cells of interest. (f) Bar graphs showing percentage of tumor cells by cluster over pseudotime. (g) Kaplan-Meier survival analysis in IDH^mut^ patient cohort. (h) Kaplan-Meier survival analysis in HGG IDH^mut^ patient cohort.

To determine the effect of GABA-OPC tumor cells on glioma progression, we assigned GABA-OPC scores to bulk RNA-seq IDH^mut^ glioma samples for which long-term survival follow up was collected^40^. Samples were split into low (n=41) and high (n=41) GABA-OPC groups based on median expression. Kaplan-Meier survival analysis showed that low GABA-OPC tumor patients had a median survival of 84.5 months whereas high GABA-OPC tumor patients showed a median survival of 156.9 months (**Fig. 5g**). Even amongst the high grade IDH^mut^ patients, low GABA-OPC scores conferred worse survival outcomes, with a median survival of 80.5 and 182.5 for low and high GABA-OPC groups, respectively (**Fig. 5h**). Collectively, these analyses confirm that GABA-OPC tumor cells are a defining feature of IDH^mut^ glioma and select subtypes of IDH^WT^ glioma and demonstrate that reduced GABA-OPC signatures confer significantly worse survival outcomes in IDH^mut^ glioma patients **(Figs. S36-37).** To our knowledge, long-term survival data for IDH^WT^ glioma with matched expression data using Neftel et al.’s classification system is not available. Accordingly, future studies should investigate the correlation of GABA-OPC signatures with prognostic outcomes in these patients.

### Tumor intrinsic depolarizations differentially alter proliferation in an IDH subtype-dependent manner

Given that GABA-OPC signatures correlate with improved survival outcomes in IDH^mut^ glioma patients, we sought to understand the effects of GABA-OPC cells on tumor cell proliferation. To do this in IDH^mut^ tumors, we utilized immunostaining with an IDH1R132H-specific antibody and OLIG2 to estimate the percentage of GABA-OPC tumor cells in four IDH^mut^ patient samples including grade II oligodendroglioma, grade II astrocytoma, grade III astrocytoma and grade IV astrocytoma. Our bioinformatics analyses revealed that approximately 85% of GABA-OPC tumor cells are OLIG2+ and that 58% of tumor cells that are not GABA-OPCs also express OLIG2 (**Fig S38**). Immunostaining analyses revealed that 38% of cells are IDH1R132H+OLIG2+; factoring in the percentages of GABA-OPCs and non-GABA-OPC tumor cells that similarly express OLIG2, we estimate roughly half of these IDH132H+OLIG2+ cells to be true GABA-OPCs, which is approximately 18-20% (**Fig. 6a**). Adding KI67 immunostaining to this analysis showed that IDH1R132H+OLIG2+ cells are largely non-proliferative, with only 3.1% double positive cells also showing KI67 positivity (**Figs. 6b-c**). Intriguingly, we noted that KI67+ cells were frequently negative for OLIG2 but retained IDH1R132H positivity, suggesting that actively proliferating cells in IDH^mut^ tumors lose *OLIG2* expression as compared to cells not undergoing G2/M transitions (**Fig. 6c-d**). Indeed, our Patch-seq analyses confirmed that the three GABA-OPCs with highest G2/M cell cycle scores show reduced *OLIG2* expression when compared to GABA-OPCs with low G2/M scores. These results are consistent with our data showing high GABA-OPC scores confer better survival outcomes in IDH^mut^ glioma and support the notion that HCs with short, tumor intrinsic APs are not largely proliferative.

**Figure 6.**
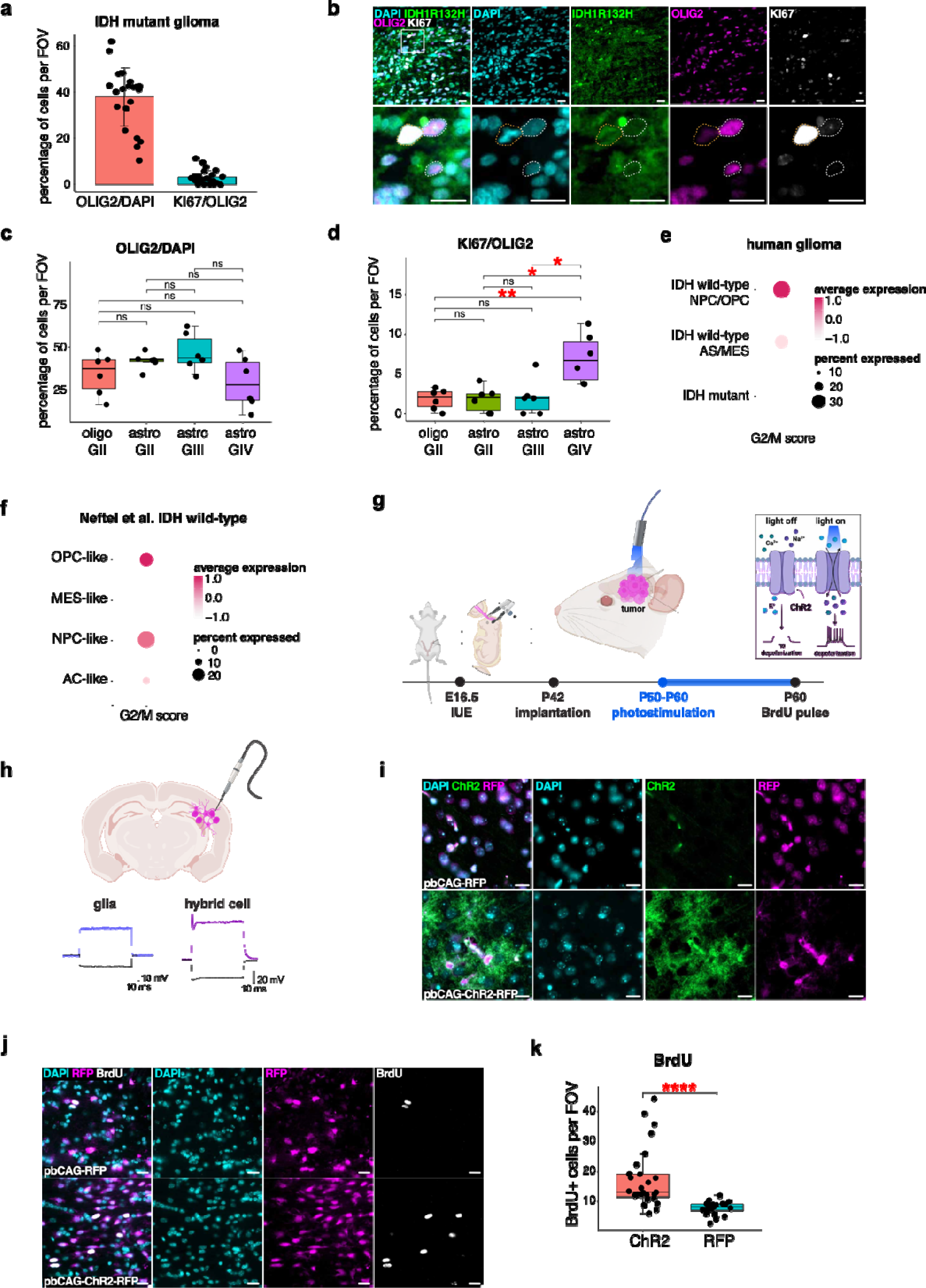
Tumor cell depolarization differentially alter glioma cell proliferation in an IDH-dependent manner. (a) Bar plot showing the percentage of IDH1R132H+OLIG2+ cells and IDH1R132H+OLIG2+KI67+ cells detected in IDH^mut^ tumor samples (n=4) using immunostaining. (b) Representative images of IDH1R132H (green), OLIG2 (pink) and KI67 (white) immunostaining shows IDH1R132H+OLIG2+ cells are largely negative for KI67. White box denotes inset; scale bar = 20 μm. (c) Box and whiskers plot showing the percentage of IDH1R132H+OLIG2+ cells detected by immunostaining for each IDH^mut^ tumor sample. (d) Box and whiskers plot showing the percentage of IDH1R132H+OLIG2+KI67+ cells detected by immunostaining for each IDH^mut^ tumor sample. (e) DotPlot of cell cycle scoring showing the percentage and average expression of GABA-OPC tumor cells undergoing G2/M in our in-house scRNA-seq dataset of human glioma. (f) DotPlot of cell cycle scoring showing the percentage and average expression of GABA-OPC tumor cells undergoing G2/M in IDH^WT^ glioma from Neftel et al.’s scRNA-seq dataset. (g) Schematic showing the experimental design used in our optogenetics experiment. (h) Schematic illustrating whole cell recording experiments in fluorescent-labeled IUE tumor mice. Representative traces of cells with glial and HC electrophysiologies are shown. (i) Representative images of optogenetic IUE tumor mice showing positive immunostaining for ChR2 (green) is detected in pbCAG-ChR2-RFP mice and is not detected in pbCAG-RFP mice; scale bar = 20 μm. (j) Representative images of optogenetic IUE tumor mice showing immunostaining for Olig2 (pink) BrdU (white); scale bar = 20 μm. (k) Box and whiskers plot showing quantification of BrdU+ cells per field of view (FOV).

While performing our bioinformatics analyses, we observed that GABA-OPCs in NPC- and OPC-like IDH^WT^ tumors showed high G2/M scores, which suggests that GABA-OPCs may have opposing effects on tumor cell proliferation that are dependent on IDH-subtype (**Figs. 6e-f; Fig Fig S39**). In contrast to the *IDH1R132H* mutation that occurs in more than 90% of IDH^mut^ gliomas, mutations occurring in IDH^WT^ tumors are heterogenous and thus antibodies specific for IDH^WT^ tumor cells are lacking. To overcome this limitation and examine the effects of tumor intrinsic depolarizations on proliferation in IDH^WT^ tumors, we employed optogenetics to induce tumor cell depolarizations using an RFP-labeled *in utero* electroporation (IUE) mouse model of *de novo* IDH^WT^ glioma^41^ (**Fig. 6g**). Critically, an analogous population of GABA-OPC tumor cells and corresponding HC electrophysiology have been identified in our IUE tumor mice, making it an appropriate model in which these experiments can be performed (**Fig. 6h; Figs. S40-41**). Overexpression of RFP with or without channelrhodopsin 2 (ChR2) was driven by piggyBac transposase in Glast-expressing progenitor cells alongside CRISPR/Cas9 guides targeting three of the most frequently mutated tumor suppressors in IDH^WT^ tumors: tumor protein 53 (TP53), phosphatase and tensin homolog (PTEN) and neurofibromin 1 (NF1). Briefly, IUEs were performed at E16.5 and fiber optic implants were placed ipsilaterally to the tumor at P40. After one week of recovery, mice received 10 consecutive days of photostimulation to induce repeated depolarizations over 10 minutes and were then pulsed with BrdU before brains were harvested for processing. Immunostaining for ChR2 and BrdU revealed that tumors expressing ChR2 (pbCAG-ChR2-RFP) were more proliferative than tumors without ChR2 (pbCAG-RFP) (**Figs. 6i-k**). These data suggest that repeated tumor cell depolarizations promote glioma cell proliferation in an IDH^WT^ context. Despite these differences in proliferation, overall survival outcomes for IDH^WT^ glioma patients based on high and low GABA-OPC scores were not significantly different, which likely reflects the smaller percentages of GABA-OPC glioma cells in these tumors as compared to IDH^mut^ tumors (**Fig. S42**). Taken together with the results of our IDH^mut^ immunostaining, these collective experiments implicate tumor cell depolarization as a differential regulator of glioma proliferation that is dependent upon the molecular and genetic context in which they occur.

## Discussion

In the 1990s, whole cell patch clamp experiments reported that cells firing single, short APs were the majority of cells found in human glioma slices. These early electrophysiology studies described spiking cells that were dependent on voltage-gated sodium currents, however, definitively showing these cells were tumor in origin necessitated the advent of single cell transcriptomics^42–44^. Separately, scientists identifying a class of spiking OPCs in the healthy rat brain posited that neurons are not the only cells capable of firing APs and suggested that an analogous population of spiking OPCs exists in human^38^. Given these precedents, we believe that these previously described cell types are electrophysiologically equivalent to our HCs, which are transcriptionally defined by GABA-OPC signatures and represent a heterogenous group malignant and non-malignant cells. There is mounting evidence to support the OPC as a cell of origin in glioma^45–47^, which represents the largest proliferative neural cell population in the adult brain and are frequently mutated in non-tumor brain^48,49^. Similarly large percentages of GABA-OPCs detected in all IDH^mut^ glioma samples used for Patch-seq, whole cell recordings and scRNA-seq in this study support the theory of OPC as cell of origin and implicate the malignant transformation of GABA-OPCs as an initiating event in IDH^mut^ and NPC- and OPC-like IDH^WT^ tumors.

Indeed, GABAergic neurons and OPCs share common neurodevelopmental origins in which most cells from each lineage emerge from Nkx2.1-expressing precursors in the medial, lateral and caudal ganglionic eminences^50,51^. In addition to emanating from the same embryonic loci, GABAergic neurons and OPCs sit at a transcriptional intersection that is uniquely shared by these two cell types, which includes the expression of OLIG2^52^ and GABARs^53^ and PDGFRA (**Fig. 4b**). These features, which are also hallmarks of GABA-OPCs, render glioma cells well-equipped to participate in the complex relay of tumor and neuronal communication that manifests as cancer neuroscience. In recent years, studies in this field have elucidated how glioma cells interact with surrounding neural networks to direct disease progression^41,54^. These reports demonstrate that human glioma cells receive synaptic inputs from the surrounding neuronal circuitry, which can be sufficient to evoke tumor cell excitatory postsynaptic currents (EPSCs)^8,9,12^. Moreover, tumor cells form intricately connected networks mediated by calcium signaling, the ablation of which limits tumor cell proliferation and progression^10^. Our studies build upon these earlier findings to demonstrate that glioma cells are capable of AP firing, raising the question of whether this tumor intrinsic activity contributes to the aberrant neurophysiology and frequent seizure incidence encountered in glioma patients. Up to 75% of IDH^mut^ glioma patients suffer from glioma-related epilepsy (GRE), which is more than double the seizure incidence in IDH^WT^ glioma patients^55,56^. Given the high percentage of GABA-OPC tumor cells in IDH^mut^ glioma, future endeavors should aim to discern whether epileptic peritumoral neuronal networks are also driven in part by tumor cell AP firing.

Perhaps one of the most intriguing and unexpected findings of our study is the discovery of two HCs from two IDH^mut^ glioma samples that retained *IDH1R132* wild-type homozygosity and euploidy despite being found within the surgically-defined core of *IDH1R132H*-mutant tumors. While these cells are few in number, an average of 5 million reads per Patch-seq cell leaves us confident that the absence of the *IDH1R132H* mutation is an accurate representation of their genomic status and clearly demonstrates that HC electrophysiology is not exclusive to a tumor state. Concordantly, the presence of three HCs detected in a histopathologically-diagnosed non-tumor sample, strongly support the conclusion that HCs are present in the non-tumor human brain. The implication of AP-firing non-neuronal cells stands as a biological iconoclast, insofar as the prevailing tenets of neuroscience hold that neurons are the only cells capable of firing APs^57^. Whether GABA-OPCs with HC electrophysiology are endemic to the healthy human brain remains to be determined, however given that OPCs are estimated to represent 3-4% of all grey matter cells and 8-9% of white matter cells in the mammalian brain^58–60^, the cumulative neurophysiological contributions of these cells are poised to be significant and should not be ignored in either tumor or non-tumor contexts.

## Supporting information

Supplemental Meterial

## Acknowledgements

This work was supported by grants from the NIH (R35-NS132230 to BD, R01NS124093 to BD and GR, R01CA223388 to BD, U01CA281902 to BD, R01NS094615 to GR). RNC is supported by grants from the NIH (5T32HL92332-15, F31CA265156, and F99CA274700). The Baylor College of Medicine Single Cell Genomics Core is supported by NIH Shared Instrument Grants (S10OD023469, S10OD025240), P30EY002520, and CPRIT grant RP200504. Schematics were created using Biorender.com.

## Data availability

In our previous study, we published the scRNA-seq datasets of 12 samples under the accession number GSE221534^32^. The Patch-seq and scRNA-seq datasets generated during this study will be made available through the NCBI Gene Expression Omnibus (GEO) website. All other study data are included in the article and/or supporting information.

## Author contributions

RNC and ASH are responsible for conception of this project, the study and pipeline design, and interpretation of the results. RNC devised all experimental set ups, performed IUEs, immunostaining and statistical analyses, and prepared all scRNA-seq samples with assistance from MFM. ASH wrote the code for the SCRAM pipeline. QM and JJ performed all human electrophysiology and Patch-seq experiments. SS and PSC performed all optogenetics experiments. RNC, AOH, SW and ASH performed bioinformatics analyses on human scRNA-seq and bulk RNA-seq datasets. BL, YJK, PH and PA provided experimental support. GR identified and obtained consent from patients for the study. RNC and ASH prepared the manuscript. BA, XJ, BD, and GR contributed to the manuscript with feedback from all authors.

## Contact for reagent and resource sharing

Dr. Akdes Serin Harmanci is the lead contact for reagent and resource sharing. All published reagents will be shared on an unrestricted basis; reagent requests should be directed to the corresponding author.

## Experimental Methods

### Human data

Adult patients at St. Luke’s Medical Center and Ben Taub General Hospital provided preoperative informed consent to participate in the study and gave consent under Institutional Review Board Protocol H35355. Patients included males and females. Clinical characteristics were maintained in a deidentified patient database and are summarized in **Table S1 and S7.**

Tumor samples were collected during surgery and immediately placed on ice. Tissue was divided for use in subsequent transcriptomic, histopathological, proteomic, or biochemical studies. Patient samples were collected separately for pathology and molecular subtyping. Histopathology and molecular subtyping of *IDH* and 1p19q deletion status were confirmed by board-certified pathologists. Samples for scRNA-seq and immunoprecipitation assays were fixed in LN_2_ and kept at −80°C.

### Single cell RNA-sequencing

Human tumors were prepared as single-cell suspensions. Briefly, samples were coarsely chopped with surgical scissors and enzymatically digested with Papain supplemented with DNase I (Worthington Biochemical Corporation, LK003150). Samples were incubated for 15 minutes at 37°C on a thermocycler kept at 1400×*g* and briefly pipetted every 5 minutes. Cells were pelleted at 13,000×*g* for 10 seconds and resuspended in phosphate-buffered saline (PBS) before processing for debris and dead cell removal. Cell suspensions were processed using the MACS Debris Removal Kit (Miltenyl, 130-109-398) and MACS Dead Cell Removal Kit (Miltenyl, 130-090-101), according to the manufacturer’s instructions. Live cells were collected through negative selection using an MS Column in the magnetic field of a MiniMACS Separator (Miltenyl, 130-042-102). Eluted cells were spun at 300×*g* for 5 minutes and resuspended in Gibco Dulbecco’s Modified Eagle Medium with GlutaMAX (DMEM; ThermoFisher, 10566016) supplemented with 10% foetal bovine serum (FBS; ThermoFisher, 16000044). Single cells were processed with the 10X Chromium 3′ Single-Cell Platform using the Chromium Single-Cell 3′ Library, Gel Bead, and Chip Kits (10X Genomics) following the manufacturer’s protocol. Briefly, approximately 5,000–15,000 cells were added to each channel of a chip to be partitioned into Gel Beads in Emulsion (GEMs) in the Chromium instrument, followed by cell lysis and barcoded reverse transcription of RNA in droplets. GEMs were broken, and cDNA from each single cell was pooled. Clean-up was performed using Dynabeads MyOne Silane Beads (ThermoFisher, 37002D). Subsequently, the cDNA was amplified and fragmented to optimal size before end repair, A-tailing, and adaptor ligation. Libraries were run individually using a NextSeq 500/550 High Output Kit v2.5 (75 Cycles) (Illumina, 20024907) and sequenced on an Illumina NextSeq550 instrument.

### Human tumor slice preparation

Fresh tumor samples were immediately placed into a cold (0−4°C) oxygenated N-methyl-d-glucamine (NMDG) solution (93 mM NMDG, 93 mM HCl, 2.5 mM KCl, 1.2 mM NaH_2_PO_4_, 30 mM NaHCO_3_, 20 mM HEPES, 25 mM glucose, 5 mM sodium ascorbate, 2 mM thiourea, 3 mM sodium pyruvate, 10 mM MgSO_4_, and 0.5 mM CaCl_2_, pH 7.35). Slices were cut at 300-μm thickness with a microslicer (Leica VT 1200) and kept at 37.0±0.5°C in oxygenated NMDG solution for 10–15 minutes before being transferred to artificial cerebrospinal fluid (ACSF, 125 mM NaCl, 2.5 mM KCl, 1.25 nM NaH_2_PO_4_, 25 mM NaHCO_3_, 1 mM MgCl_2_, 25 mM glucose, and 2 mM CaCl_2_, pH 7.4) for 1 hour before recording.

### Single cell processing

We ran samples on the 10X Chromium platform to produce next-generation sequencing libraries. We performed standard procedures for filtering, mitochondrial gene removal, and variable gene selection using the Seurat pipeline. The criteria for cell/gene inclusion were as follows: genes present in more than three cells were included, cells that expressed >300 genes were included, the number of genes detected in each cell was >200 and <5000, and the mitochondria ratio was 10. We integrated cells from different patients using the Harmony algorithm^61^. Next, we visualized clusters using a uniform manifold approximation and projection constructed from the Harmony-corrected PCA. We also performed lineage tracing, trajectory analysis, and RNA velocity assessments to create developmental hierarchies and lineage histories of glioma cells using the scvelo R package^62^ and IntrExtract^63^.

### SCRAM pipeline and methodology

SCRAM input consisted of aligned scRNA-seq reads and our neural network model trained on 11 diverse single-cell RNA-Seq datasets encompassing 1 million cells of publicly available data from healthy adult and developing brain samples, as well as brain tumor samples. Tumor and normal cells were annotated independently for two reasons. (1) Significant overlap exists between tumor and non-tumor expression markers. For example, *EGFR* and *PDGFRA* are often used to denote tumor cells^14^ but are also cell type markers for OPCs and ependymal host cells, respectively (**Supplemental Figure S43**) (2) We hypothesize that by separating tumor-specific and normal-specific features, we can achieve more robust identification of hybrid cells (HCs). This hypothesis is supported by our observation that existing cell type assignment methods, which typically classify both tumor and normal cells together, fail to accurately characterize HCs. These tumor/normal features were systematically employed in our pipeline as follows:

#### Step 1. Annotation of non-tumor cells

##### Training Neural Network Models (NNMs)

We trained our neural network model (NNMs) on 11 diverse single-cell RNA-Seq datasets, which collectively contain 1 million cells. These datasets comprise publicly available data sourced from various datasets, including healthy adult and developing brain samples, as well as brain tumor samples^13–15,23–30^. We trained our model using a deep neural network (DNN), with an input layer of around 20K genes, three intermediate layers (256, 64, 32), and an output layer of size 16 or 21, depending on the number of referenced cell types. Following the dense connection within each hidden layer, there are batch normalization, activation, and dropout functions. We use the popular Rectified Linear Unit (ReLU) for hidden layer activation and set dropout rate to be 0.1. The output layer uses Softmax activation function so that each node outputs a non-negative value smaller than 1 and all the values sum up to 1. Therefore, each output corresponds to the probability of one cell type. We compile the model using categorical crossentropy as loss function, Adam as optimizer, and accuracy as metrics. In order to achieve a more balanced class distribution, we opted to subsample cell types within our training model. We train one neural network-based classifier on each developmental-like, normal, tumor cells datasets and save the model in repository. We predicted the developmental-like, normal and tumor cells, in our glioma scRNA-Seq data using our trained NNMs. Model probability scores >0.9 were used for final cell annotations. In building our NNMs, we utilized the Python packages TensorFlow and Keras. Additionally, we used the Python Scipy package for processing the scRNA-Seq data. Prerequisite packages for data preprocessing and model training include Numpy 1.19.5, Pandas 1.1.5, Scanpy 1.7.2, Anndata 0.7.8, Scipy 1.5.4, and Scikit-learn 0.24.2.

For a single cell to be classified as “immune” within our framework, it must be annotated as immune with a probability score above 0.9 in three or more trained datasets.

##### SHapley Additive exPlanations (SHAP) analysis

SHAP analysis was employed to gain insights into the main features that affect the output of the NNMs, we used SHAP method, to explain how each feature affects the NNMs in inhouse glioma single-cell dataset. To perform the SHAP analysis, the model predictions were decomposed into contributions from individual features, allowing us to assess their impact on the final outcome using the *shap* python package.

#### Step 2. Annotation of tumor cells

The SCRAM pipeline integrates multiple orthogonal tumor features to identify tumor cells at a single-cell resolution. These features include:

##### Module 1. Neural Network Model-Based Tumor Cell Prediction

Above explained NNMs is used to predict tumor cells based on the probability score. Cells with a probability score above 0.9 are classified as tumor cells.

##### Module 2. Marker based Expression Modeling

SCRAM employs finite Gaussian mixture modeling to model marker expression of three tumor marker genes: *SOX2, EGFR PDGFRA*. This approach helps to distinguish tumor cells based on their specific marker gene expression profiles (details explained below section “*Marker Expression Modeling for tumor annotation”*).

##### Module 3. RNA-Inferred Genotyping of Chromosome Alterations

A modified version of our CNV-calling algorithm, CaSpER^21^, another state of art CNV calling method numbat^22^ is used in SCRAM to perform RNA-inferred genotyping of large-scale chromosome alterations.

##### Module 4. RNA-Inferred Mutational Profiling

SCRAM utilizes our XCVATR^20^ tool, a recently developed tool, to deduce rare deleterious single-nucleotide variants (SNVs) present in the tumor cells. This analysis involves considering SNVs that are reported in the COSMIC^64^-database and have a frequency of less than 0.1% in the dbSNP^65^database.

These orthogonal tumor features are called separately in the SCRAM pipeline. By combining these different approaches, SCRAM aims to accurately identify tumor cells at a single-cell resolution.

For a single cell to be classified as “tumor” within our framework, it needs to meet two or more of the following criteria:

1. Neural Network Model-Based Tumor Cell Prediction (Module 1): The cell is annotated as “tumor” when it receives a probability score greater than 0.9 from the trained Neural Network Models (NNMs) in Richards et al.^15^, or Venteicher et al.^24^ or Neftel et al.^14^ or Tirosh et al.^13^ datasets.
2. Marker Expression Modeling (Module 2): The expression levels of at least two tumor cell markers, (*SOX2*, *EGFR*, or *PDGFRA*) should surpass a predetermined threshold. This threshold is established using finite Gaussian mixture modeling (details explained below section “*Marker Expression Modeling for tumor annotation”*), as depicted in **Supplemental Figure S43**.
3. RNA-Inferred Genotyping of Chromosome Alterations (Module 3): The presence of large-scale copy number variations (CNVs) is considered a tumor cell.
4. RNA-Inferred Mutational Profiling (Module 4): Tumor cells that have SNVs in genes *IDH* (R132H/R132C) or EGFR.

###### Module 2. Marker Expression Modeling for tumor annotation

Given the expressional heterogeneity of tumor markers in normal cells, we used previously published tumor and non-tumor cell datasets to establish a marker expression–based tumor classification model (i.e., thresholding requirements for “high expression” annotation) for the tumor markers *PDGFRA, EGFR,* and *SOX2*. For each tumor marker, an independent classifier model was built using (1) Allen Brain Atlas human scRNA-seq data, which represent the largest compendium of healthy brain data as a training set for normal cells; and (2) a compendium of publicly available brain tumor scRNA-seq datasets as a training set for tumor cells^14^. The following statistical models were used to infer the class (normal vs tumor) of our in-house tumor scRNA-seq data.

We modelled expression as a mixture of Gaussian distributions to identify and classify normal and tumor cells:

Let X_j_ = { x_1_,x_2_,…,x_i_,…,x_n_} be the training expression vector of normal and tumor cells for gene j, where x_i_ is the expression value of cell i. The distribution of every expression value is specified by a probability density function through a finite mixture model of *G*=2 classes (normal vs tumor):

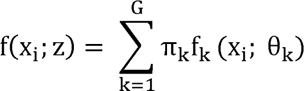

where z= {π_1,….,_π_G_, θ_1_,…θ_G_} represents the parameters of the mixture model and *f_k_* (*x_i_*; *θ_k_*) is the kth component density, which is assumed to follow a Gaussian distribution *f_k_*(*x_t_*; *θ_k_*) ∼ *N*(*µ_k_*,*σ_k_*). {*π*_1,….,_*π_G_*} is the vector of probabilities, non-negative values that sum to 1, known as the mixing proportion. The mixing proportion, π, follows a multinomial distribution.

We used the above model to predict normal vs tumor class in our in-house glioma cells. For each gene, j, z parameters were estimated by maximizing the log-likelihood function via the expectation-maximization algorithm. The log-likelihood function is formulated as:

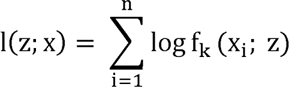

For each tumor marker, we generated a matrix, with genes indicated by rows and cells indicated by columns, and the cell value index was 1 if the cell had a high “tumor class” probability for the corresponding gene. A cell was classified as “tumor” if at least two markers had high “tumor class” probabilities. We used the mclust R package for Gaussian mixture model implementation^66^.

###### Module 3. RNA-Inferred Genotyping of Chromosome Alterations

CNVs are hallmark features of tumor cells that can be used to classify tumor vs non-tumor cells with or without expression markers. However, CNV detection from scRNA-seq data is inherently noisy due to dropouts and unmatched control sets, among other factors, requiring a set of known tumor cells. To estimate a “clean” set of CNV calls to provide reliable CNV-based tumor scores, we used a *pure tumor pseudobulk* sample.

##### Estimation of CNV profiles using patient-specific pure tumor pseudobulk samples

We first used our marker expression–based and NNMs models from Module 1 and Module 2 to identify tumor cells. Cells assigned as “tumor” cells using Module 1 and 2 were treated as a pure tumor cell cohort. Cells assigned as “immune” cells using our NNMs are considered control cells.

##### CNV calling of patient-specific pure pseudobulk samples

We hypothesized that the pseudobulk sample contained representative sets of CNVs with high probability and, therefore, should be useful to identify a clean CNV call set. CNV calling of the pseudobulk samples from each patient was performed using our CNV-calling algorithm, CaSpER. CaSpER CNV calls were used as the ground truth, large-scale CNV calls for each patient.

##### Genotyping of CNVs of all cells

After CNVs were identified from the pseudobulk sample, we genotyped the CNVs in all cells and generated a binary matrix that represents the existence of CNVs in cells, i.e., *CNV_i,j_*.

###### Module 4. RNA-Inferred Mutational Profiling

We performed RNA-inferred rare deleterious (COSMIC^64^-reported and dbSNP^65^, <0.1% frequency) mutational profiling via our recently developed XCVATR^20^ tool. We detected mutations in *IDH (R132H/R132C)*, *EGFR* and annotated cells harboring these mutations as tumor cells.

### Visualization and clustering of single cells

We used the probability score output from NNMs instead of relying solely on the most variable genes for clustering and visualization of our in-house single-cell data. This, named as SCRAM UMAP, involved applying UMAP and clustering techniques to the model probability scores using the Seurat package’s runUMAP, FindNeighbors, and FindClusters functions. Additionally, we employed the most variable genes for cell data clustering and visualization, referring to this UMAP representation as the original/Seurat UMAP.

### Summarizing co-occurring cell types using maximally frequent gene set identification

We summarized co-occurring cell types using a frequent itemset rule mining approach. CNV and SNV calls were added to provide an integrated transcriptomic and genomic summary for each cell. An example SCRAM output for a single cell is given as “glioma stem cell, oligodendrocyte precursor cell, chr1p_deletion, chr19q_deletion *+ IDH:2:208248389* mutation”. We used the tumor and normal cell assignments of Step 1 and Step 2 to integrate co-occurring tumor and normal cell features.

The simplest method for detecting maximally frequent tumor and host feature sets is a brute force approach in which each possible subset of features is a candidate frequent set. The *a priori* algorithm is an efficient implementation for finding maximally frequent sets with support above a given threshold. In the *a priori* algorithm, the minimum *support* threshold is set to min (50, number_cells_in cluster*0.1), and the maximum number of genes in a gene set is set to 50. Using the *a priori* algorithm, we identified co-occurring gene sets expressed concurrently within each cell and provided annotation of high-resolution cellular identities using a three-step co-occurrence analysis. We performed our co-occurrence analysis at multiple levels: 1) cell type level (example output of this step: *tumor* AND *radial glia* AND *astrocyte*); 2) cell class level (example output of this step: *tumor* AND *neural cells* are commonly upregulated).

#### Maximally frequent cell type (or cell class) co-occurrence analysis

Within each cluster, *m,* we calculated the maximally frequent cell types (or cell lineage or cell class) using the *a priori* algorithm. The input was the binarized matrix *E^m^*, where the cell types (or cell lineage or cell class) were the rows, and the cells in cluster *m* were the columns.

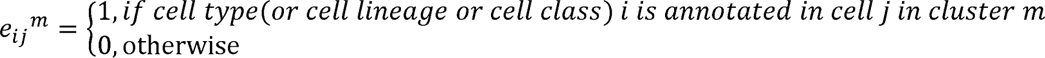

### Analyzing bulk expression data and survival analysis

TCGA-GBM (high grade glioma), TCGA-LGG (low grade glioma) raw read counts and accompanying clinical data are downloaded using TCGAbiolinks R package^67^. TCGA-GBM, TCGA-LGG and our bulk RNA-Seq data of the IDH Mutant cohort were both normalized and applied variance stabilizing transformation using the DESeq2 package^68^. Single sample gene set enrichment analysis (ssGSEA) was performed using GSVA R package. We used our GABA-OPC tumor gene sets and also the MES-like, AC-like, NPC-like and OPC-like gene sets reported in a previous study^14^. SsGSEA GABA-OPC scores were split by median to assign high-OPC-GABA and low-OPC-GABA scored samples. Those groups are compared against “overall survival” in a Cox Proportional Hazards (Cox) survival model. used in survival analysis and compared using a Log-rank test P-value. We used survminer and survival R package for the survival analysis.

### Optogenetics

This experimental setup forces expression of ChR in tumor cells, which will be activated by light using fiber optic implants, resulting in the depolarization of tumor cells. ChR causes depolarization of cells by allowing sodium to flow into the cell when in the presence of light. Mice were implanted with fiber optic cables at P42 and began light stimulation sessions at P50. Our protocol was modified from Venkatesh et al.^7^ for use in our IUE system. Briefly, light pulses at 20 Hz, 473 nm, and 5mWatt for 30 seconds were administered followed by 90 seconds of recovery, over 10 minutes for 10 consecutive days. Following final stimulation, mice were injected with a single 200mg/kg BrdU pulse and harvested 1 hour later.

### Patch-seq recording procedures

Electrophysiological, morphological, and transcriptomic data from the same cell were obtained simultaneously using the Patch-seq protocol described previously^16,69^. Briefly, patch pipettes (5−7 MΩ) were filled with RNase-free intracellular solution (111 mM potassium gluconate, 4 mM KCl, 10 mM HEPES, 0.2 mM EGTA, 4 mM MgATP, 0.3 mM Na_3_GTP, 5 mM sodium phosphocreatine, and 13.4 mM biocytin). Whole-cell recordings were performed using a Quadro EPC 10 amplifier (HEKA Electronic). After 5–10 minutes of whole-cell recording of firing patterns, the nucleus was extracted using gentle and continuous negative pressure. The contents in the pipette were ejected into a 0.2-mL PCR tube containing 4 mL lysis buffer^69^. RNA in the lysis buffer was denatured, reverse transcribed, amplified, and purified following the Smart-seq2-based protocol^70^. Only high-quality cDNA samples (yield ≥2 ng, average length ≥1500 bp) were sequenced.

Sequencing libraries were constructed from the cDNA using the Illumina Nextera XT DNA Library Preparation Kit (Illumina, FC-131-1096). The cDNA library was sequenced on a NovaSeq 6000 instrument using 150-bp paired-end reads.

### Biocytin staining and morphological reconstruction

Following slice recording, slices were fixed by immersion in the fixation solution at 4°C for at least 48 hours and processed with an avidin-biotin-peroxidase method to reveal the cell morphology. The morphology of the cells was reconstructed and analysed using a 100× oil-immersion objective lens and camera lucida system (Neurolucida, MicroBrightField).

### Histology

Human samples were retrieved from the operating room on ice and then fixed in 4% paraformaldehyde in PBS for 12 hours at 4°C before being transferred to 70% EtOH. Paraffin embedding was performed by the Breast Cancer Pathology Core at Baylor College of Medicine. All human specimens were evaluated by a board-certified neuropathologist according to current guidelines and standard practices.

### Immunostaining

For immunostaining, 10 μm paraffin-embedded human glioma sections were cut, deparaffinized and subject to heat-induced epitope retrieval (HIER) using antigen retrieval buffer (10 mM sodium citrate, 0.05% Tween 20, pH 6.0) when needed. Sections were blocked for 1 hour at room temperature and kept in primary antibody incubation overnight at 4°C. The following primary antibodies were used: rat anti-BrdU (1:200; Abcam, ab6326), mouse anti-ChR2 (1:100, Progen, 651180), mouse anti-IDHR132H (1:50; Dianova, DIA-H09), rabbit anti-GAD1 (1:200; Synaptic Systems, 198013), goat anti-OLIG2 (1:100; R&D, AF2418), rabbit anti-RFP (1:1000; Rockland, 600-401), rabbit anti-SCN1A (Na_V_1.1) (1:200; Alomone Labs, ASC-001). Species-specific secondary antibodies tagged with Alexa Fluor corresponding to emission spectra 488 nm, 568 nm, or 647 nm (1:1,000, ThermoFisher) were used for immunofluorescence and Hoechst nuclear counter staining (1:50,000; ThermoFisher, H3570) was performed before coverslipping with Vectashield antifade mounting medium (Vector Laboratories, H-1000). For quantification, n≥3 biological samples were used. For imaging, n≥3 images were taken per tissue section × n≥3 sections × n≥3 biological samples.

### Patch-seq data processing

The Patch-seq reads were mapped using STAR^71^ to hg38 assemblies for humans. Read count matrices were generated using FeatureCounts^72^ with the latest gene annotations from GENCODE^73^ consortia. DEGs and transcripts were identified using DESeq2^68^ and limma^74^. Cells were clustered and visualized using PCA methods. Cell type enrichment analyses were performed with enrichR^75^ using the *PanglaoDB_Augmented_2021* and *CellMarker_Augmented_2021* cell type marker sets. *IDH* mutations were identified using our variant detection tool, XCVATR^20^, and visually confirmed using the Integrative Genomics Viewer^76^.

### Statistical analysis

For electrophysiology analyses, a Kruskal–Wallis test or two-way ANOVA was used, followed by unpaired t-tests with a two-stage step-up (Benjamini, Krieger, and Yekutieli). For RT-qPCR, a two-tailed Student’s t-test was used. Significant differences are denoted by asterisks in associated graphs. Data are presented as the mean±standard error of the mean. Levels of statistical significance are indicated as follows: ns: not significant, \**p*<0.05, \*\**p*<0.01, \*\*\**p*<0.001, and \*\*\*\**p*<0.0001.

## References

1. Mesfin, F. B. & Al-Dhahir, M. A. Gliomas. (StatPearls Publishing, 2023).

2. Stupp, R. et al. Radiotherapy plus concomitant and adjuvant temozolomide for glioblastoma. N. Engl. J. Med. 352, 987–996 (2005).

3. Yan, H., et al. *IDH1*and*IDH2*mutations in gliomas. N. Engl. J. Med. 360, 765–773 (2009).

4. Cancer Genome Atlas Research Network. Comprehensive genomic characterization defines human glioblastoma genes and core pathways. Nature 455, 1061–1068 (2008).

5. Verhaak, R. G. W. et al. Integrated genomic analysis identifies clinically relevant subtypes of glioblastoma characterized by abnormalities in PDGFRA, IDH1, EGFR, and NF1. Cancer Cell 17, 98–110 (2010).

6. Charles, N. A., Holland, E. C., Gilbertson, R., Glass, R. & Kettenmann, H. The brain tumor microenvironment. Glia 59, 1169–1180 (2011).

7. Venkatesh, H. S. et al. Neuronal activity promotes glioma growth through neuroligin-3 secretion. Cell 161, 803–816 (2015).

8. Venkatesh, H. S. et al. Electrical and synaptic integration of glioma into neural circuits. Nature 573, 539–545 (2019).

9. Venkataramani, V. et al. Glutamatergic synaptic input to glioma cells drives brain tumour progression. Nature 573, 532–538 (2019).

10. Venkataramani, V. et al. Glioblastoma hijacks neuronal mechanisms for brain invasion. Cell 185, 2899–2917.e31 (2022).

11. Tantillo, E. et al. Differential roles of pyramidal and fast-spiking, GABAergic neurons in the control of glioma cell proliferation. Neurobiol. Dis. 141, 104942 (2020).

12. Barron, T. et al. GABAergic neuron-to-glioma synapses in diffuse midline gliomas. (2022) doi:10.1101/2022.11.08.515720.

13. Tirosh, I. et al. Single-cell RNA-seq supports a developmental hierarchy in human oligodendroglioma. Nature 539, 309–313 (2016).

14. Neftel, C. et al. An Integrative Model of Cellular States, Plasticity, and Genetics for Glioblastoma. Cell 178, 835–849.e21 (2019).

15. Richards, L. M. et al. Gradient of Developmental and Injury Response transcriptional states defines functional vulnerabilities underpinning glioblastoma heterogeneity. Nat Cancer 2, 157–173 (2021).

16. Cadwell, C. R. et al. Electrophysiological, transcriptomic and morphologic profiling of single neurons using Patch-seq. Nat. Biotechnol. 34, 199–203 (2016).

17. Lipovsek, M. et al. Patch-seq: Past, present, and future. J. Neurosci. 41, 937–946 (2021).

18. Jiang, X. et al. Principles of connectivity among morphologically defined cell types in adult neocortex. Science 350, aac9462 (2015).

19. Scala, F. et al. Phenotypic variation of transcriptomic cell types in mouse motor cortex. Nature 598, 144–150 (2021).

20. Harmanci, A. O., Harmanci, A. S., Klisch, T. & Patel, A. J. XCVATR: Characterization of Variant Impact on the Embeddings of Single -Cell and Bulk RNA-Sequencing Samples. Preprint at 10.1101/2021.06.01.446668.

21. Harmanci, A. S., Harmanci, A. O. & Zhou, X. CaSpER identifies and visualizes CNV events by integrative analysis of single-cell or bulk RNA-sequencing data. Nature Communications vol. 11 Preprint at 10.1038/s41467-019-13779-x (2020).

22. Gao, T. et al. Haplotype-aware analysis of somatic copy number variations from single-cell transcriptomes. Nat. Biotechnol. 41, 417–426 (2023).

23. Bakken, T. E. et al. Comparative cellular analysis of motor cortex in human, marmoset and mouse. Nature 598, 111–119 (2021).

24. Venteicher, A. S. et al. Decoupling genetics, lineages, and microenvironment in IDH-mutant gliomas by single-cell RNA-seq. Science 355, (2017).

25. Bhaduri, A. et al. An atlas of cortical arealization identifies dynamic molecular signatures. Nature 598, 200–204 (2021).

26. Polioudakis, D. et al. A single-cell transcriptomic atlas of human neocortical development during mid-gestation. Neuron 103, 785–801.e8 (2019).

27. Pombo Antunes, A. R., et al. Single-cell profiling of myeloid cells in glioblastoma across species and disease stage reveals macrophage competition and specialization. Nat. Neurosci. 24, 595–610 (2021).

28. Domínguez Conde, C., et al. Cross-tissue immune cell analysis reveals tissue-specific features in humans. Science 376, eabl5197 (2022).

29. Aldinger, K. A. et al. Spatial and cell type transcriptional landscape of human cerebellar development. Nat. Neurosci. 24, 1163–1175 (2021).

30. Sjöstedt, E. et al. An atlas of the protein-coding genes in the human, pig, and mouse brain. Science 367, (2020).

31. Ganguly, K., Schinder, A. F., Wong, S. T. & Poo, M. GABA itself promotes the developmental switch of neuronal GABAergic responses from excitation to inhibition. Cell 105, 521–532 (2001).

32. Curry, R. N. et al. Glioma epileptiform activity and progression are driven by IGSF3-mediated potassium dysregulation. Neuron 111, 682–695.e9 (2023).

33. Marques, S. et al. Oligodendrocyte heterogeneity in the mouse juvenile and adult central nervous system. Science 352, 1326–1329 (2016).

34. Marques, S. et al. Transcriptional convergence of oligodendrocyte lineage progenitors during development. Dev. Cell 46, 504–517.e7 (2018).

35. Bai, X., Kirchhoff, F. & Scheller, A. Oligodendroglial GABAergic signaling: More than inhibition! *Neurosci*. Bull. 37, 1039–1050 (2021).

36. Luyt, K. et al. Developing oligodendrocytes express functional GABA(B) receptors that stimulate cell proliferation and migration. J. Neurochem. 100, 822–840 (2007).

37. Zhang, X. et al. NG2 glia-derived GABA release tunes inhibitory synapses and contributes to stress-induced anxiety. Nat. Commun. 12, 5740 (2021).

38. Káradóttir, R., Hamilton, N. B., Bakiri, Y. & Attwell, D. Spiking and nonspiking classes of oligodendrocyte precursor glia in CNS white matter. Nat. Neurosci. 11, 450–456 (2008).

39. Ge, W.-P. et al. Long-term potentiation of neuron-glia synapses mediated by Ca2+-permeable AMPA receptors. Science 312, 1533–1537 (2006).

40. Lee, S. et al. Role of CX3CR1 signaling in malignant transformation of gliomas. Neuro. Oncol. 22, 1463–1473 (2020).

41. Yu, K. et al. PIK3CA variants selectively initiate brain hyperactivity during gliomagenesis. Nature 578, 166–171 (2020).

42. Labrakakis, C. et al. Action potential-generating cells in human glioblastomas. J. Neuropathol. Exp. Neurol. 56, 243–254 (1997).

43. Patt, S. et al. Neuron-like physiological properties of cells from human oligodendroglial tumors. Neuroscience 71, 601–611 (1996).

44. Labrakakis, C., Patt, S., Hartmann, J. & Kettenmann, H. Glutamate receptor activation can trigger electrical activity in human glioma cells. Eur. J. Neurosci. 10, 2153–2162 (1998).

45. Galvao, R. P. et al. Transformation of quiescent adult oligodendrocyte precursor cells into malignant glioma through a multistep reactivation process. Proc. Natl. Acad. Sci. U. S. A. 111, E4214–23 (2014).

46. Liu, C. et al. Mosaic analysis with double markers reveals tumor cell of origin in glioma. Cell 146, 209–221 (2011).

47. Monje, M. et al. Hedgehog-responsive candidate cell of origin for diffuse intrinsic pontine glioma. Proc. Natl. Acad. Sci. U. S. A. 108, 4453–4458 (2011).

48. Ganz, J. et al. Rates and patterns of clonal oncogenic mutations in the normal human brain. Cancer Discov. 12, 172–185 (2022).

49. Geha, S. et al. NG2+/Olig2+ cells are the major cycle-related cell population of the adult human normal brain. Brain Pathol. 20, 399–411 (2010).

50. Kessaris, N. et al. Competing waves of oligodendrocytes in the forebrain and postnatal elimination of an embryonic lineage. Nat. Neurosci. 9, 173–179 (2006).

51. Wamsley, B. & Fishell, G. Genetic and activity-dependent mechanisms underlying interneuron diversity. Nat. Rev. Neurosci. 18, 299–309 (2017).

52. Miyoshi, G., Butt, S. J. B., Takebayashi, H. & Fishell, G. Physiologically distinct temporal cohorts of cortical interneurons arise from telencephalic Olig2-expressing precursors. J. Neurosci. 27, 7786–7798 (2007).

53. Benamer, N., Vidal, M. & Angulo, M. C. The cerebral cortex is a substrate of multiple interactions between GABAergic interneurons and oligodendrocyte lineage cells. Neurosci. Lett. 715, 134615 (2020).

54. John Lin, C.-C., et al. Identification of diverse astrocyte populations and their malignant analogs. Nat. Neurosci. 20, 396–405 (2017).

55. Kerkhof, M., Benit, C., Duran-Pena, A. & Vecht, C. J. Seizures in oligodendroglial tumors. CNS Oncol. 4, 347–356 (2015).

56. Correia, C. E., Umemura, Y., Flynn, J. R., Reiner, A. S. & Avila, E. K. Pharmacoresistant seizures and IDH mutation in low-grade gliomas. Neurooncol Adv 3, vdab146 (2021).

57. Fields, R. D. Oligodendrocytes changing the rules: Action potentials in Glia and oligodendrocytes controlling action potentials. Neuroscientist 14, 540–543 (2008).

58. Dawson, M. R. L., Polito, A., Levine, J. M. & Reynolds, R. NG2-expressing glial progenitor cells: an abundant and widespread population of cycling cells in the adult rat CNS. Mol. Cell. Neurosci. 24, 476–488 (2003).

59. Levine, J. M., Reynolds, R. & Fawcett, J. W. The oligodendrocyte precursor cell in health and disease. Trends Neurosci. 24, 39–47 (2001).

60. Beiter, R. M. et al. Evidence for oligodendrocyte progenitor cell heterogeneity in the adult mouse brain. Sci. Rep. 12, 12921 (2022).

61. Korsunsky, I. et al. Fast, sensitive and accurate integration of single-cell data with Harmony. Nat. Methods 16, 1289–1296 (2019).

62. Bergen, V., Lange, M., Peidli, S., Wolf, F. A. & Theis, F. J. Generalizing RNA velocity to transient cell states through dynamical modeling. Nat. Biotechnol. 38, 1408–1414 (2020).

63. Harmanci, A. S., et al. scRegulocity: Detection of local RNA velocity patterns in embeddings of single cell RNA-Seq data. bioRxiv (2021) doi:10.1101/2021.06.01.446674.

64. Tate, J. G. et al. COSMIC: The Catalogue Of Somatic Mutations In Cancer. Nucleic Acids Res. 47, D941–D947 (2019).

65. Sherry, S. T. et al. dbSNP: the NCBI database of genetic variation. Nucleic Acids Res. 29, 308–311 (2001).

66. Scrucca, L., Fop, M., Murphy, T. B. & Raftery, A. E. Mclust 5: Clustering, classification and density estimation using Gaussian finite mixture models. R J. 8, 289–317 (2016).

67. Mounir, M., et al. Analyses of cancer data in the Genomic Data Commons Data Portal with new functionalities in the TCGAbiolinks R/Bioconductor package. bioRxiv (2018) doi:10.1101/350439.

68. Love, M. I., Huber, W. & Anders, S. Moderated estimation of fold change and dispersion for RNA-seq data with DESeq2. Genome Biol. 15, 550 (2014).

69. Cadwell, C. R. et al. Multimodal profiling of single-cell morphology, electrophysiology, and gene expression using Patch-seq. Nat. Protoc. 12, 2531–2553 (2017).

70. Picelli, S. et al. Full-length RNA-seq from single cells using Smart-seq2. Nat. Protoc. 9, 171–181 (2014).

71. Dobin, A. et al. STAR: ultrafast universal RNA-seq aligner. Bioinformatics 29, 15–21 (2013).

72. Liao, Y., Smyth, G. K. & Shi, W. featureCounts: an efficient general purpose program for assigning sequence reads to genomic features. Bioinformatics 30, 923–930 (2014).

73. Harrow, J. et al. GENCODE: The reference human genome annotation for The ENCODE Project. Genome Res. 22, 1760–1774 (2012).

74. Ritchie, M. E. et al. limma powers differential expression analyses for RNA-sequencing and microarray studies. Nucleic Acids Res. 43, e47 (2015).

75. Kuleshov, M. V. et al. Enrichr: a comprehensive gene set enrichment analysis web server 2016 update. Nucleic Acids Res. 44, W90–7 (2016).

76. Robinson, J. T. et al. Integrative genomics viewer. Nat. Biotechnol. 29, 24–26 (2011).

