## Supplemental Meterial for "Integrated electrophysiological and genomic profiles of single cells reveal spiking tumor cells in human glioma"

#### Supplemental Figures & Figure Legends

#### Extended Data Figure S1

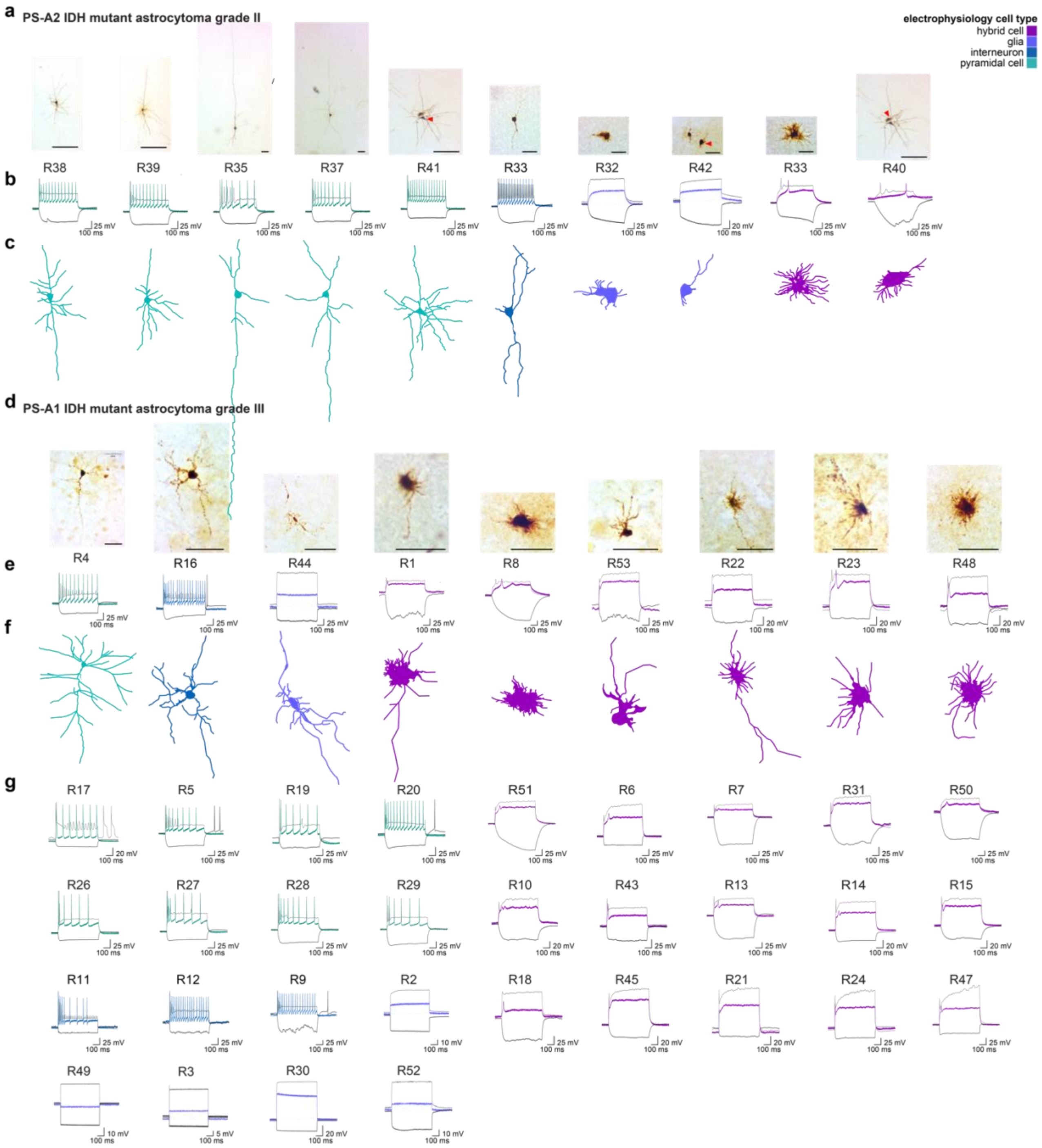

**Extended Data Figure S1. Cell Morphologies and Membrane Responses in Patched Cells from Patients PS-A1 and PS-A2** (a) Images of biocytin-reconstructed cell morphologies for patched cells from an IDH mutant (IDH$^{mut}$) diffuse astrocytoma (grade II); scale bar: 50 μm. (b) Membrane responses to a 600-ms hyperpolarizing current step (black) and a suprathreshold depolarizing current step (colored). (c) Traced cell morphologies are shown for recorded cells. (d) Images of biocytin-reconstructed cell morphologies for patched cells from an IDH$^{mut}$ diffuse astrocytoma (grade III); scale bar: 50 μm. (e) Membrane responses to a 600-ms hyperpolarizing current step (black) and a suprathreshold depolarizing current step (colored). (f) Traced cell morphologies are shown for recorded cells. (g) Continuation of *d*: Membrane responses to a 600-ms hyperpolarizing current step (black) and a suprathreshold depolarizing current step (colored). Three voltage traces are shown: the hyperpolarization trace obtained with injected currents (black), the first depolarization trace (grey), and the depolarization trace showing maximal AP firing rate; injected current: −100 pA. Traces are listed below corresponding images unless otherwise noted.

18 **Extended Data Figure S2**

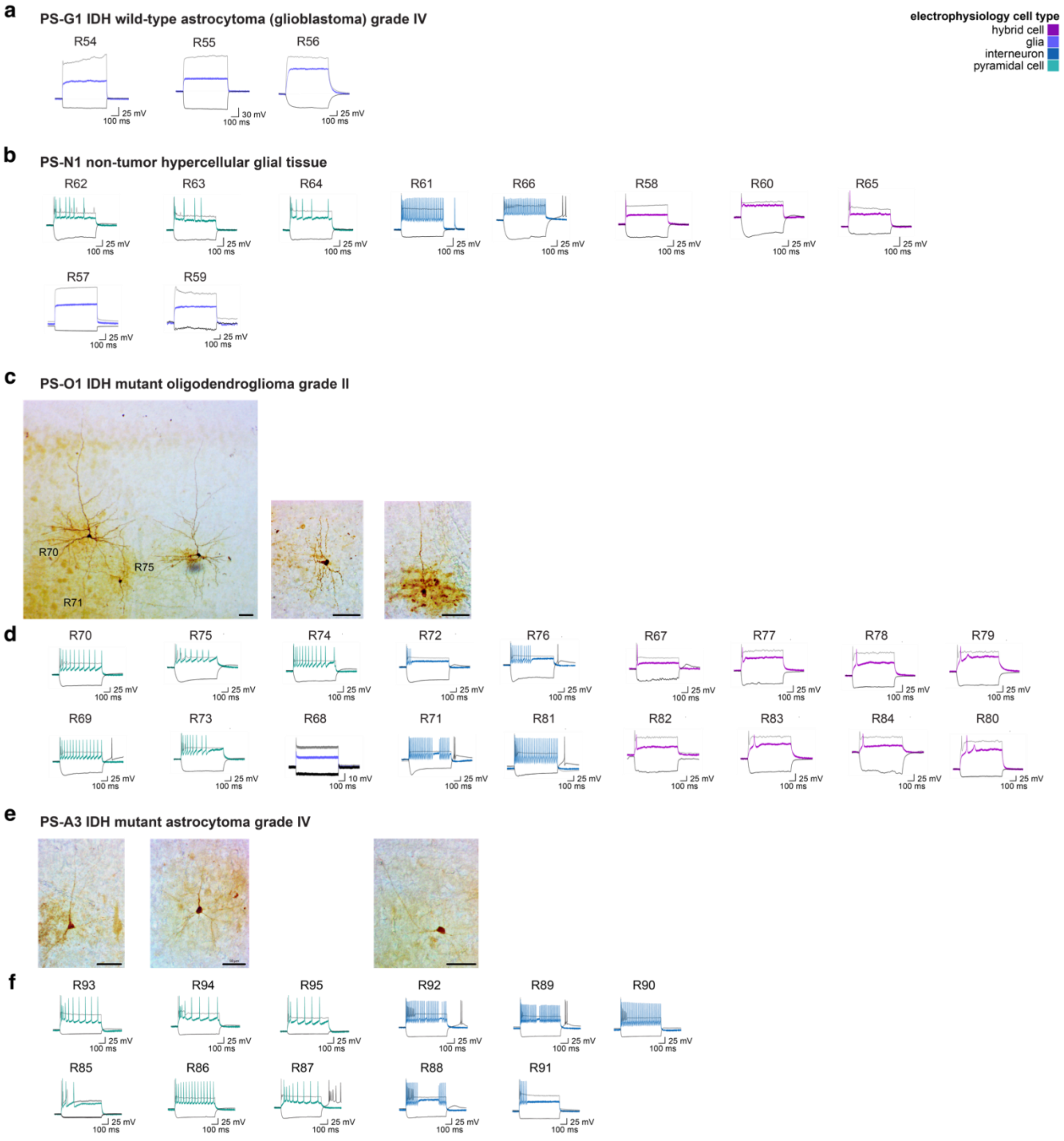

**Extended Data Figure S2. Cell Morphologies and Membrane Responses in Patched Cells from Patients PS-G1, PS-N1, PS-O1 and PS-A3.** (a) Membrane responses to a 600-ms hyperpolarizing current step (black) and a suprathreshold depolarizing current step (colored) for a IDH wild-type ($IDH^{WT}$) astrocytoma (glioblastoma; grade IV). (b) Membrane responses to a 600-ms hyperpolarizing current step (black) and a suprathreshold depolarizing current step (colored) for a non-tumor sample. (c) Images of biocytin-reconstructed cell morphologies for patched cells from an IDH mutant ($IDH^{mut}$) oligodendroglioma (grade II); scale bar: 50 μm. (d) Membrane responses to a 600-ms hyperpolarizing current step (black) and a suprathreshold depolarizing current step (colored). (e) Images of biocytin-reconstructed cell morphologies for patched cells from an $IDH^{mut}$ astrocytoma (grade IV); scale bar: 50 μm. (f) Membrane responses to a 600-ms hyperpolarizing current step (black) and a suprathreshold depolarizing current step (colored). Three voltage traces are shown: the hyperpolarization trace obtained with injected currents (black), the first depolarization trace (grey), and the depolarization trace showing maximal AP firing rate; injected current: −100 pA. Traces are listed below corresponding images unless otherwise noted.

35 **Extended Data Figure S3**

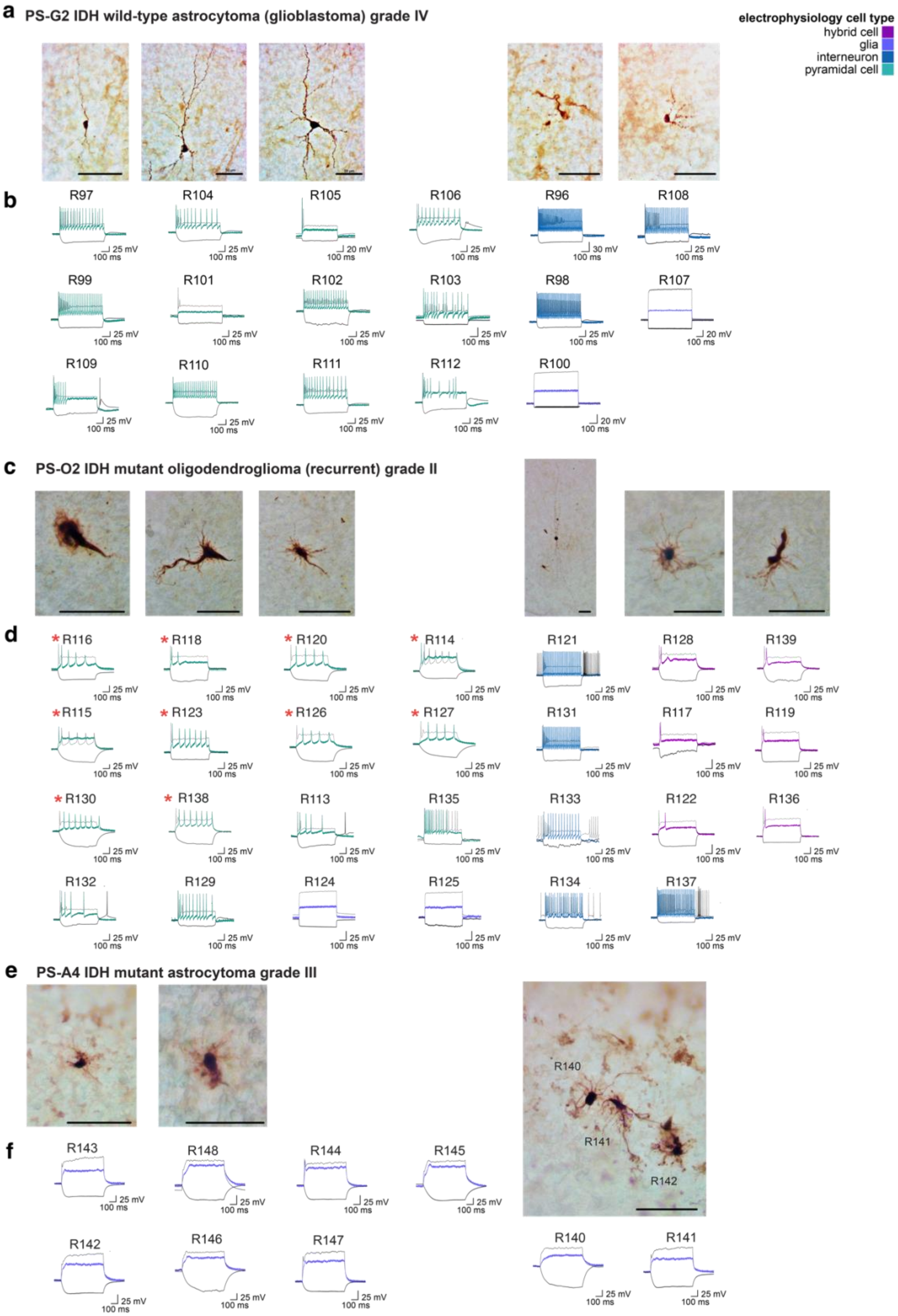

**Extended Data Figure S3. Cell Morphologies and Membrane Responses in Patched Cells from Patients PS-G2, PS-O2 and PS-A4.** (a) Images of biocytin-reconstructed cell morphologies for patched cells from an IDH wild-type ($IDH^{WT}$) astrocytoma (glioblastoma; grade IV); scale bar: 50 μm. (b) Membrane responses to a 600-ms hyperpolarizing current step (black) and a suprathreshold depolarizing current step (colored). (c) Images of biocytin-reconstructed cell morphologies for patched cells from a recurrent IDH mutant ($IDH^{mut}$) oligodendroglioma (grade II); scale bar: 50 μm. (d) Membrane responses to a 600-ms hyperpolarizing current step (black) and a suprathreshold depolarizing current step (colored); ∆PCs are denoted by red asterisks. (e) Images of biocytin-reconstructed cell morphologies for patched cells from an $IDH^{mut}$ astrocytoma (grade III); scale bar: 50 μm. (f) Membrane responses to a 600-ms hyperpolarizing current step (black) and a suprathreshold depolarizing current step (colored). Three voltage traces are shown: the hyperpolarization trace obtained with injected currents (black), the first depolarization trace (grey), and the depolarization trace showing maximal AP firing rate; injected current: −100 pA. Traces are listed below corresponding images unless otherwise noted.

#### Extended Data Figure S4

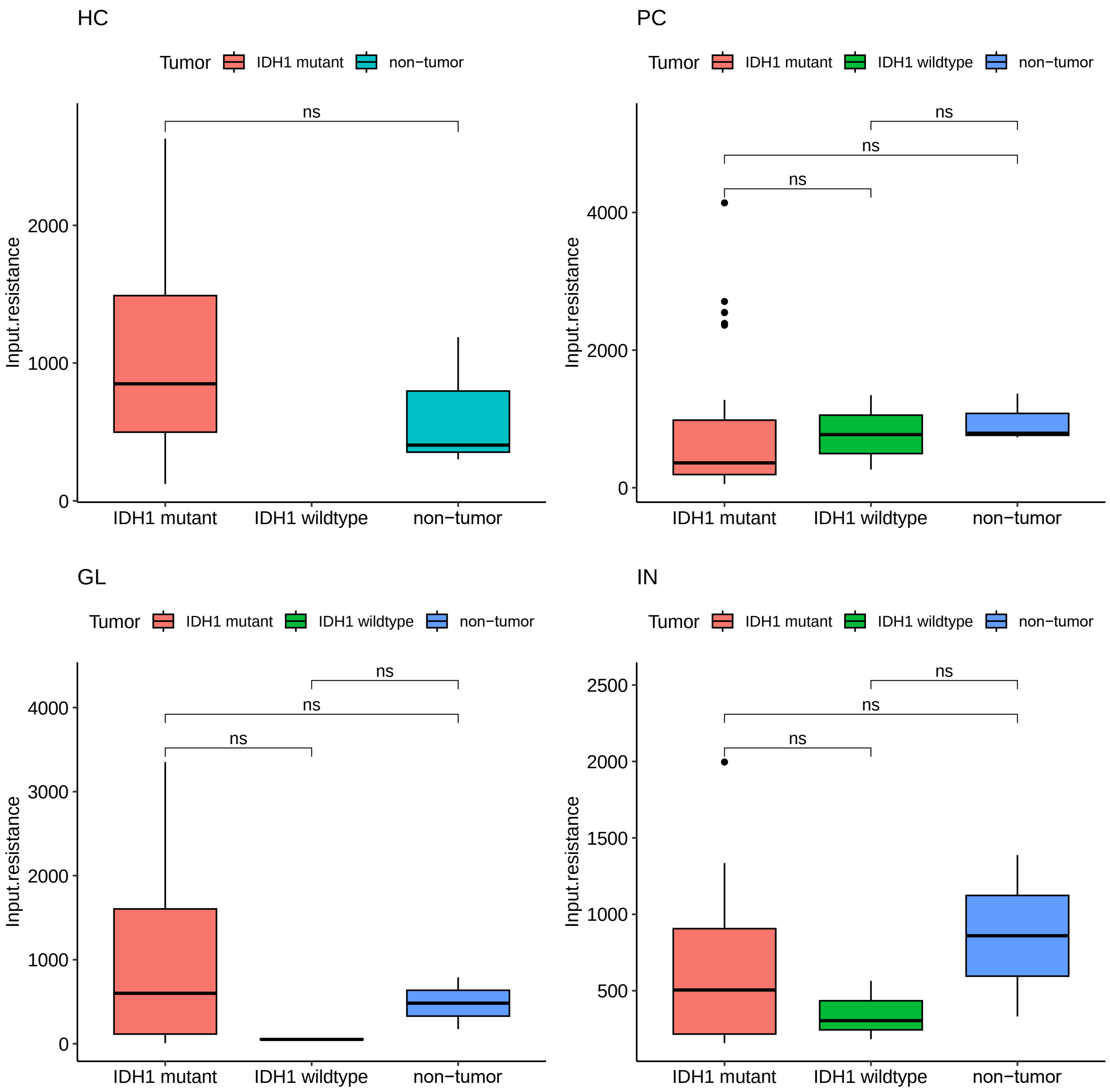

**Extended Data Figure S4. Input Resistance Differences Among Tumor Subtypes within Each Cell Type.** Box plot showing input resistance differences among each tumor subtype within HC, PC, GL and IN cells.

#### Extended Data Figure S5

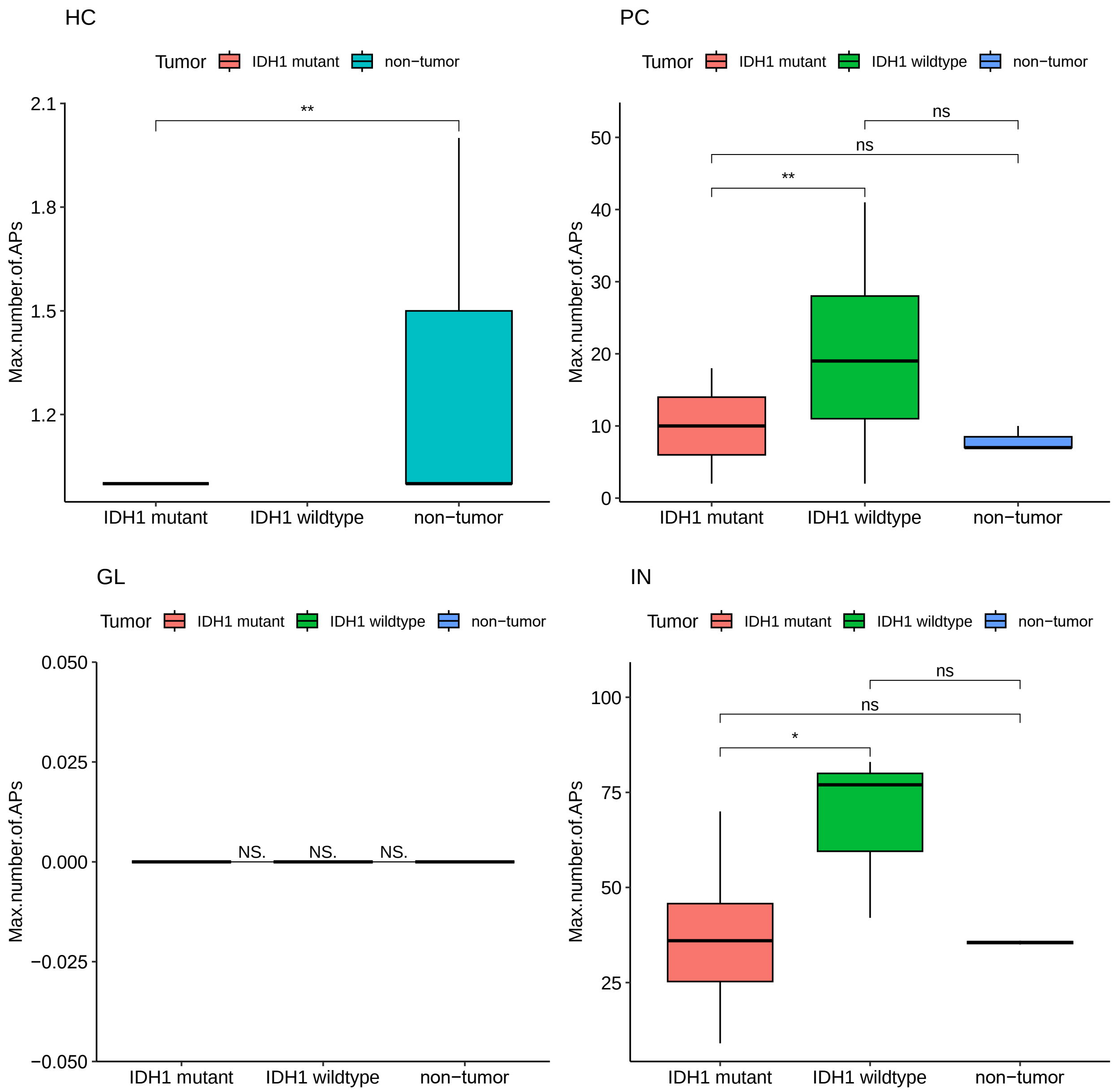

**Extended Data Figure S5. Max Number of Action Potentials (APs) Differences Among Tumor Subtypes within Each Cell Type.** Box plot showing max number of APs differences among each experimental group within HC, PC, GL and IN cells.

#### Extended Data Figure S6

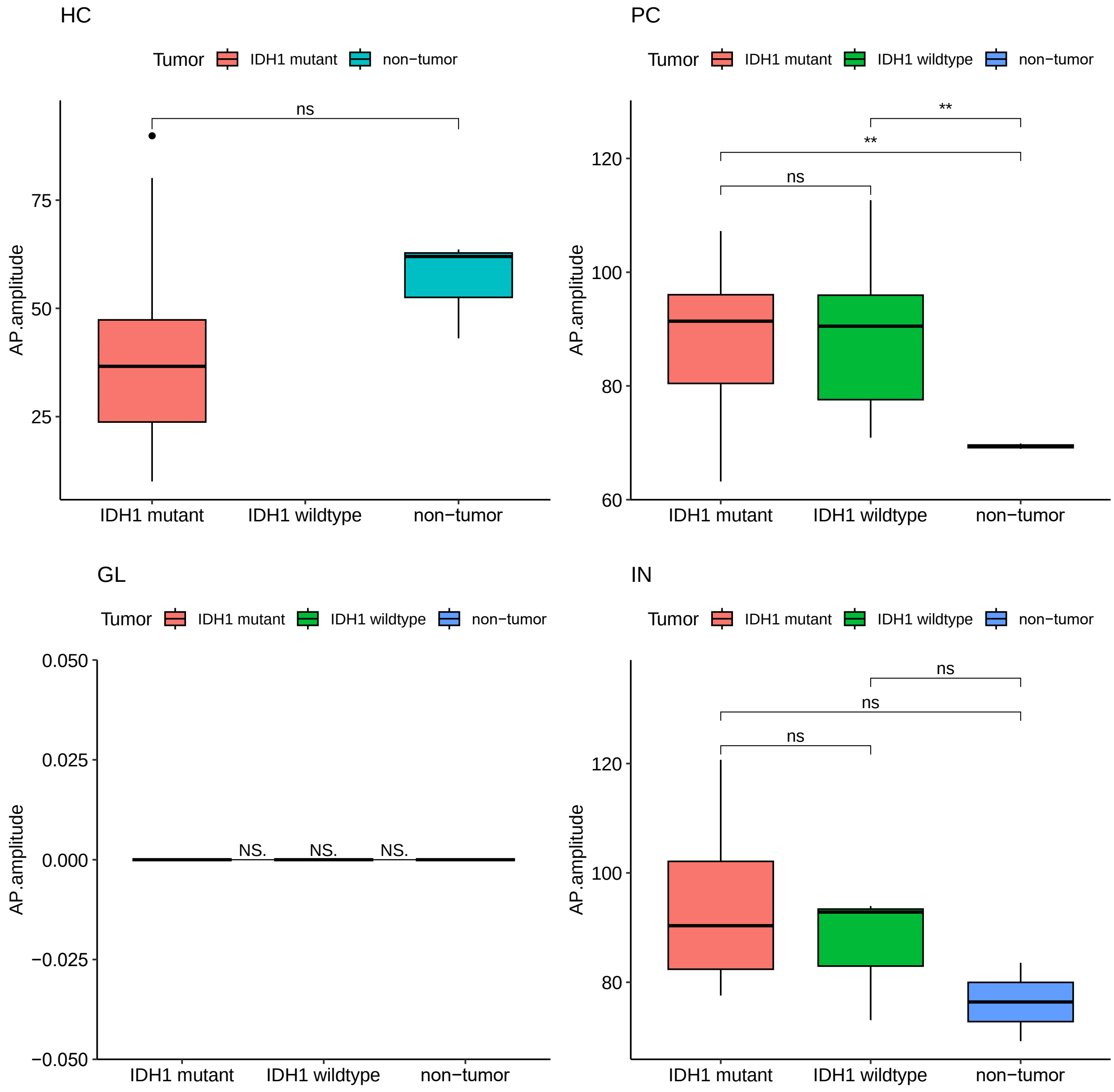

**Extended Data Figure S6. AP Amplitude Difference Among Tumor Subtypes within Each Cell Type.** Box plot showing AP amplitude differences among each experimental group within HC, PC, GL and IN cells.

**Extended Data Figure S7**

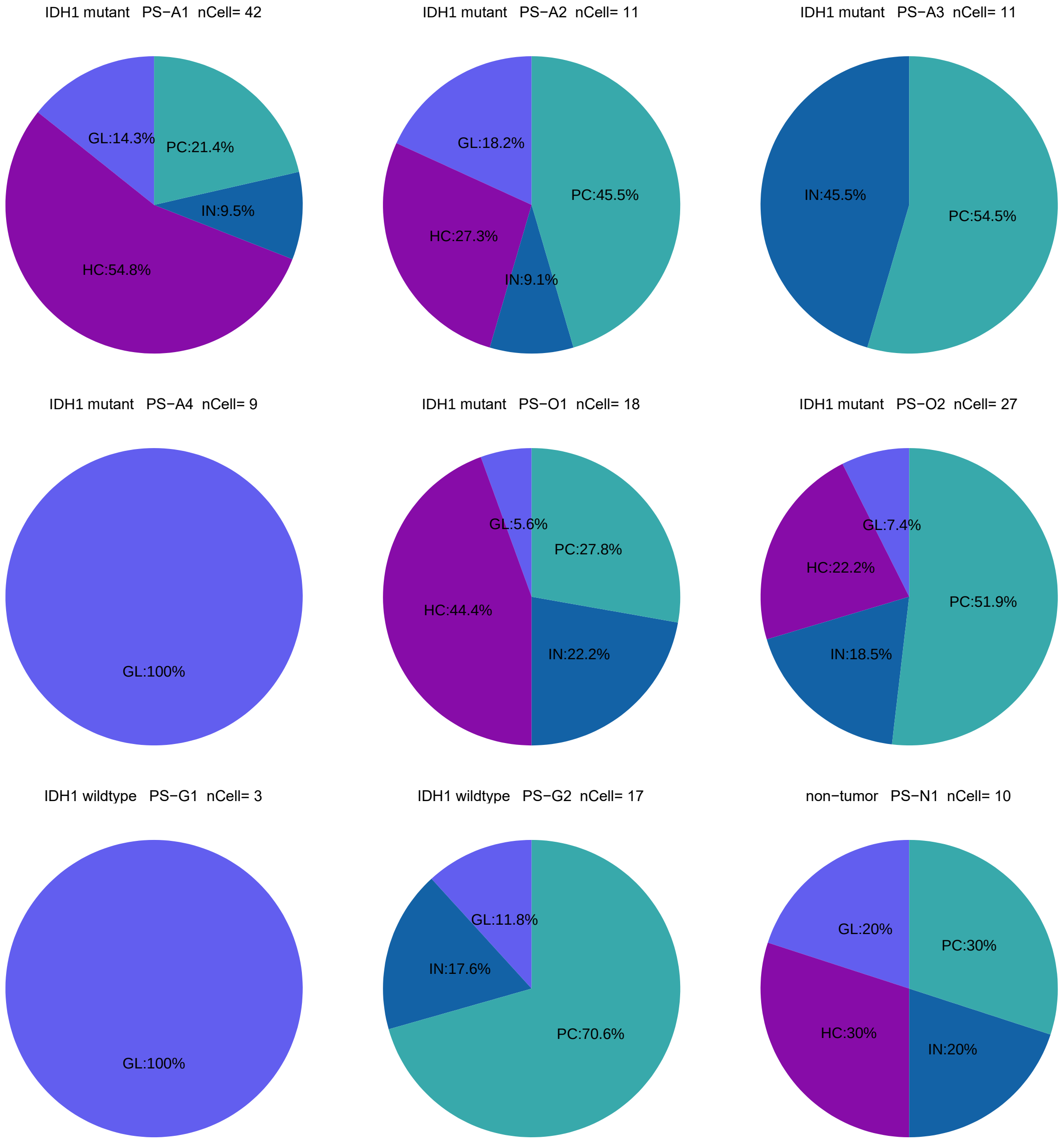

**Extended Data Figure S7. Summary of Cell Types for Each Patient.** Pie chart showing percentage of PC, IN, GL and HC patched by each patient sample.

### Extended Data Figure S8

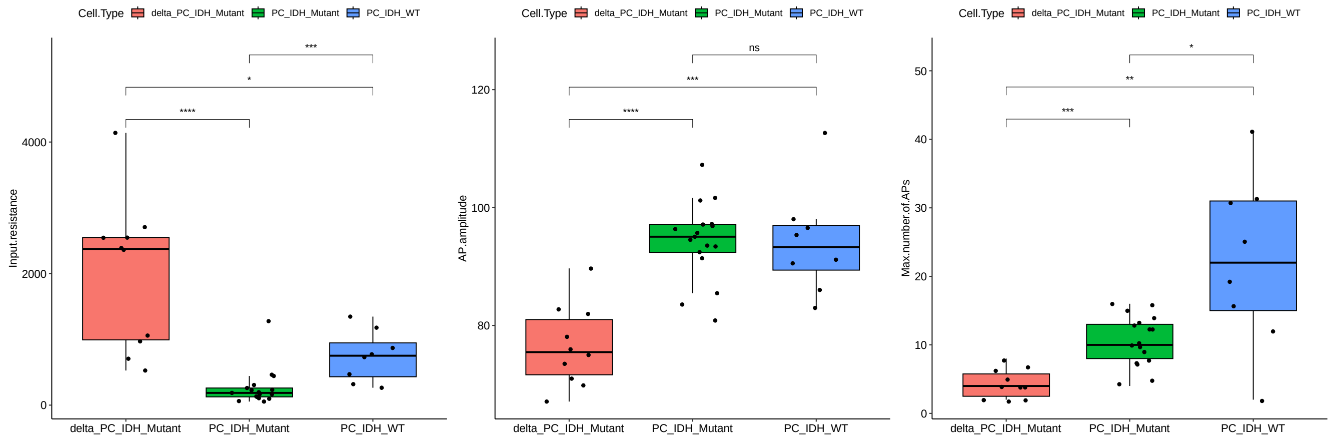

**Extended Data Figure S8. Differences in Input Resistance, AP Amplitude, and Maximum Number of APs Between Delta PCs and Other PCs.** Box plot showing the differences in input resistance, action potential (AP) amplitude, and maximum number of APs between delta PCs and other PCs.

#### Extended Data Figure S9

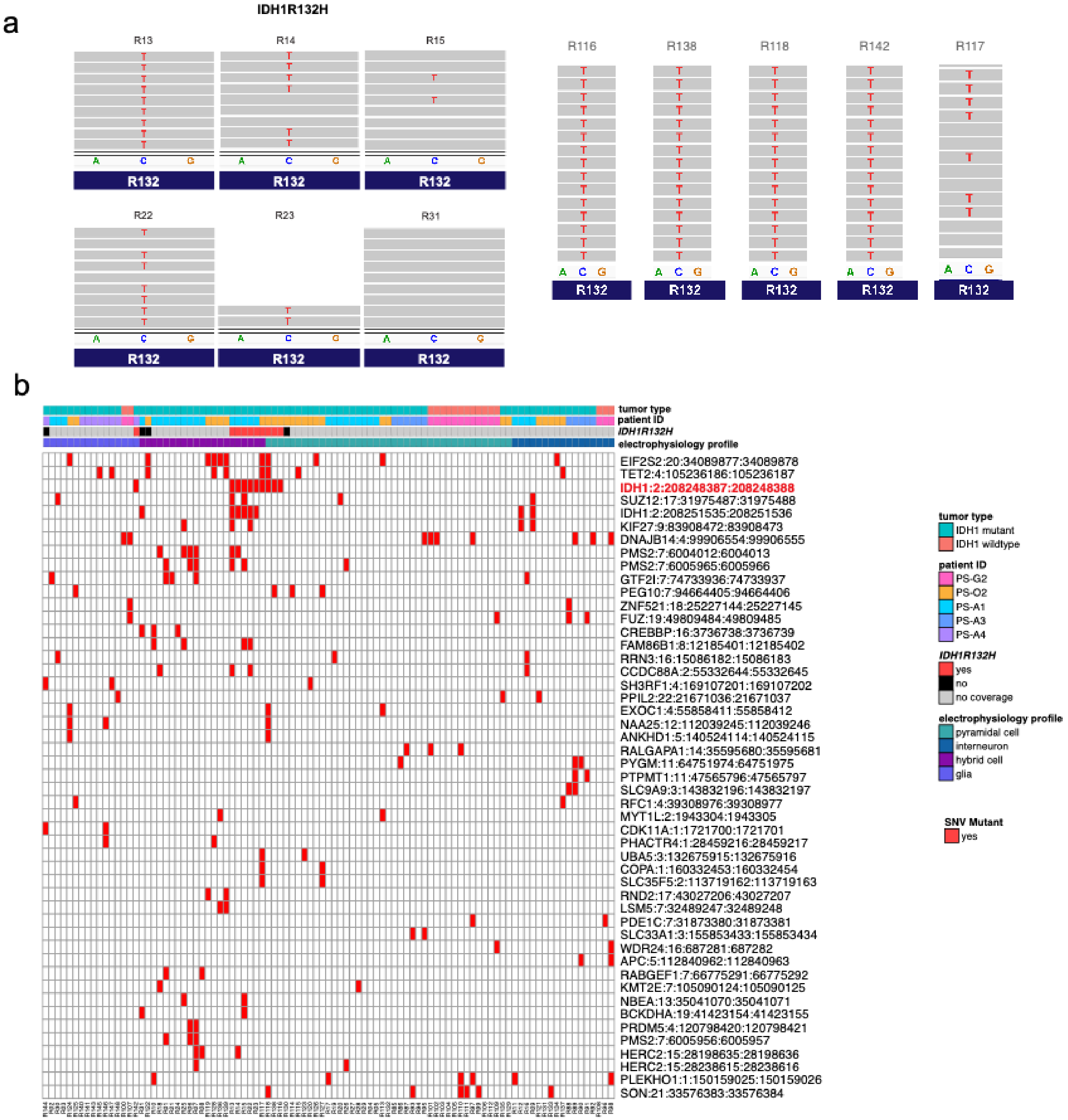

**Extended Data Figure S9. Single-Nucleotide Variant (SNV) and IDH1R132H Mutation Calls in Patch-seq** (a) Examples of whole-cell patch clamp recordings, followed by singe-cell RNA-sequencing (Patch-seq) sequencing results showing the *IDH1R132H* mutation is detected in ten of HC cells with coverage of the *IDH1* locus. (b) Whole-cell patch clamp recordings, followed by singe-cell RNA-sequencing (Patch-seq) single-nucleotide variant (SNV) analysis is shown for 95 Patch-seq cells.

#### Extended Data Figure S10

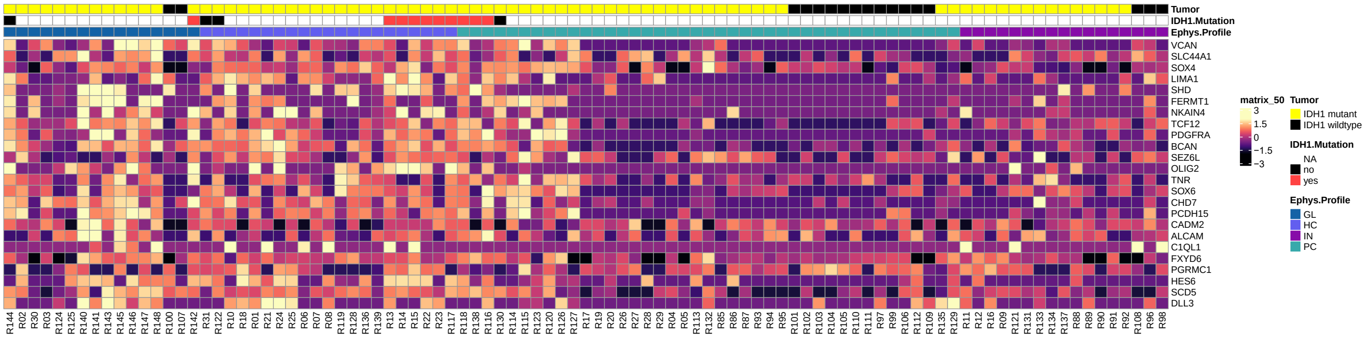

**Extended Data Figure S10. OPC genes are highly expressed in GABA-OPC tumor cells.** Heatmap of Patch-seq cells showing enrichment of OPC genes in hybrid cells (GABA-OPCs).

**Extended Data Figure S11**

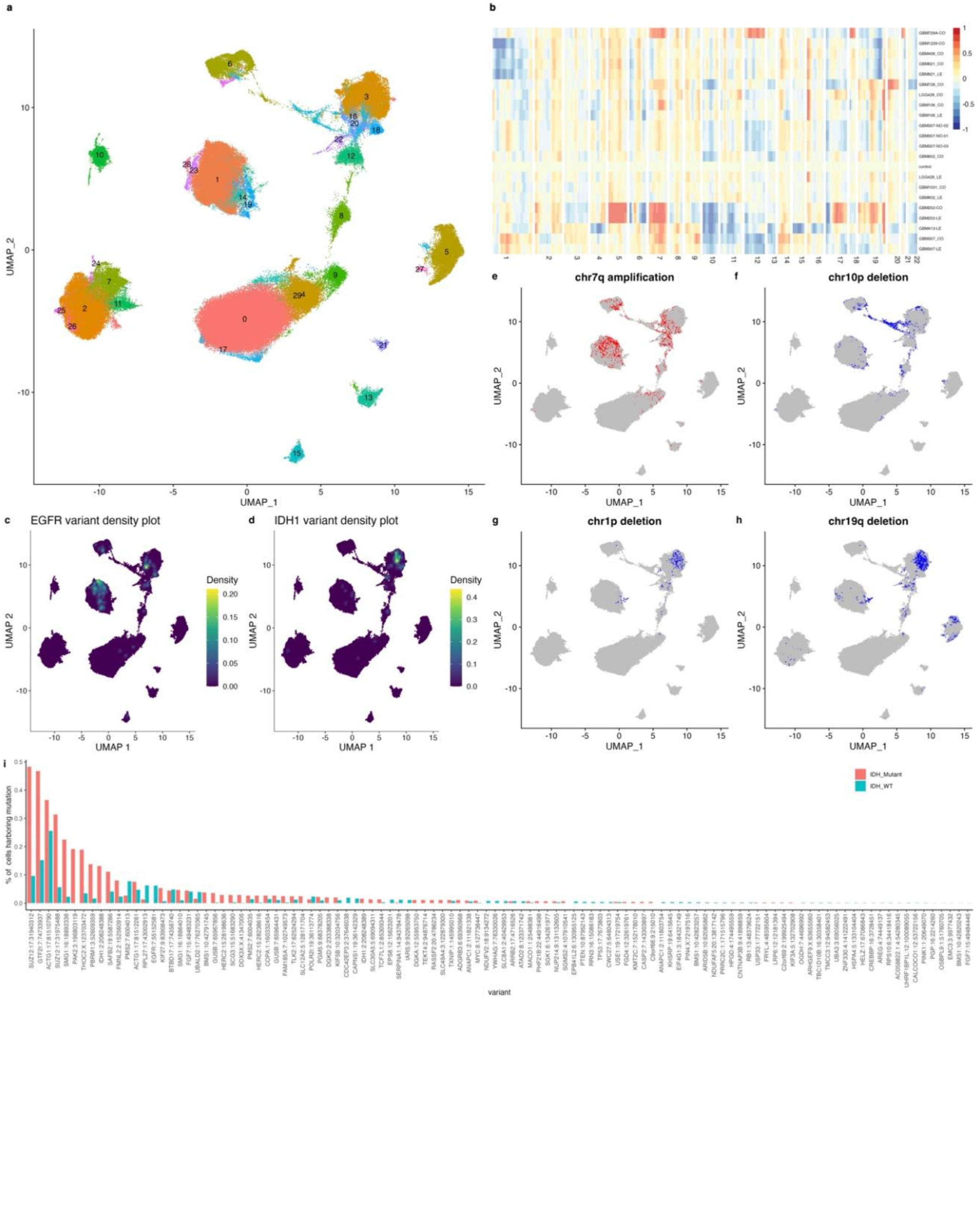

**Extended Data Figure S11. Single-Nucleotide Variant (SNV) and Copy-Number Variant (CNV) Analysis in IDH1$^{WT}$ and IDH1$^{mut}$ Glioma Subtypes Using Single-Cell RNA-Sequencing Data** (a) Seurat clusters are shown for 234,880 cells comprising our integrated *IDH1*–wild-type (IDH1$^{WT}$) and *IDH1*-mutant (IDH1$^{mut}$) single-cell RNA-sequencing glioma dataset (n=12 patients). (b) Smoothed expression signal of pseudobulk samples used for CNV calling. Rows represent genes ordered by chromosomal location. (c) Density plot showing *EGFR* mutations are detected in IDH1$^{WT}$ samples. (d) Density plots showing *IDH1* mutations are detected in IDH1$^{mut}$ samples. (e-f) Feature plots show SCRAM detects large-scale chromosome 7p amplifications and chromosome 10p deletions in IDH1$^{WT}$ glioma using CaSpER. (g-h) Feature plots show SCRAM detects large-scale chromosome 1p19q co-deletions in IDH1$^{mut}$ oligodendroglioma using CaSpER. (i) Bar plots showing the percentage of cells harboring rare and COSMIC SNVs in IDH1$^{WT}$ and IDH1$^{mut}$ glioma.

#### Extended Data Figure S12

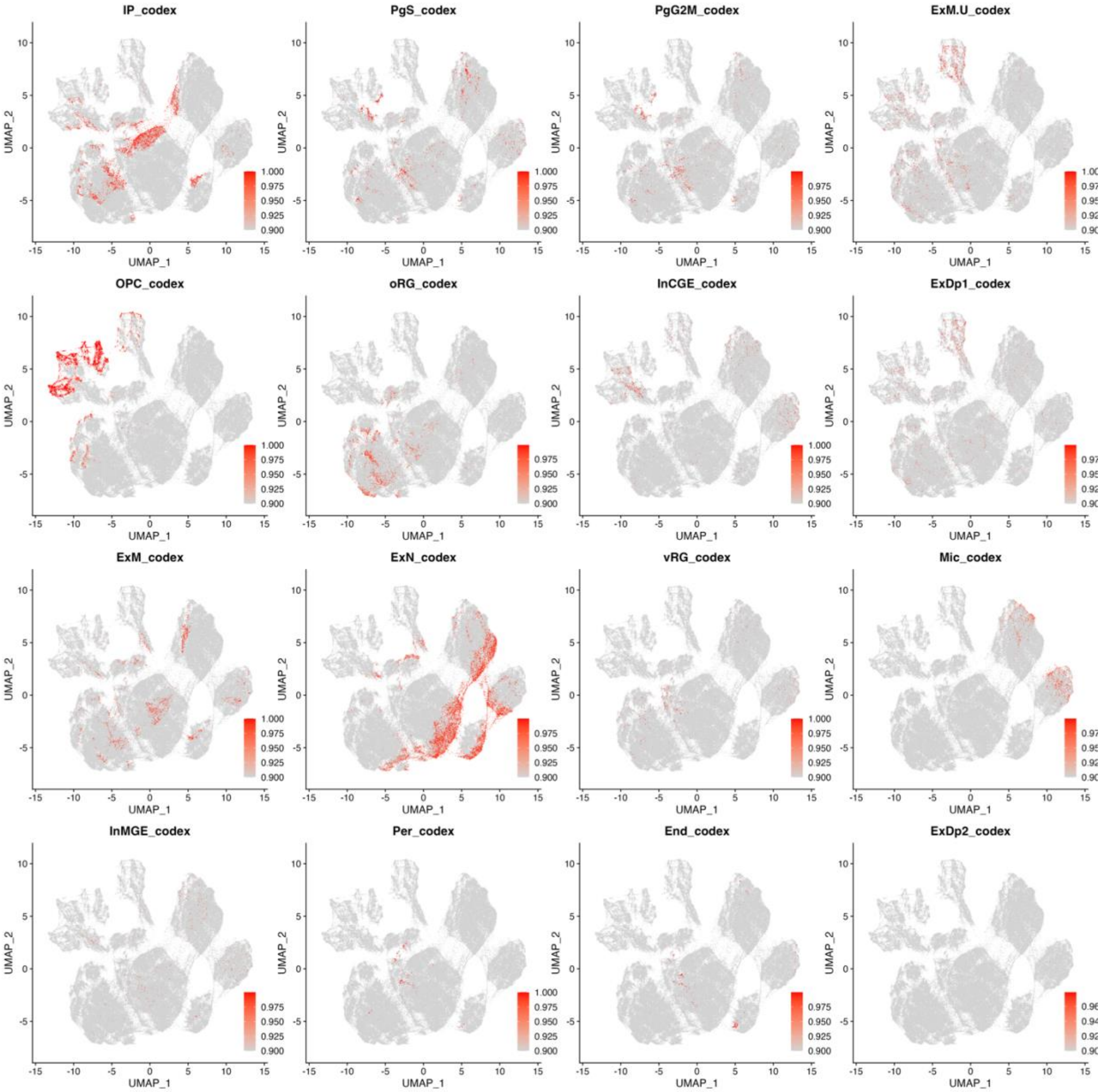

**Extended Data Figure S12. Neural Network Model Score Plot for CoDEx Cell Types.**

Neural Network model score plot for CoDEx[1] cell types on glioma SCRAM UMAP.

#### Extended Data Figure S13

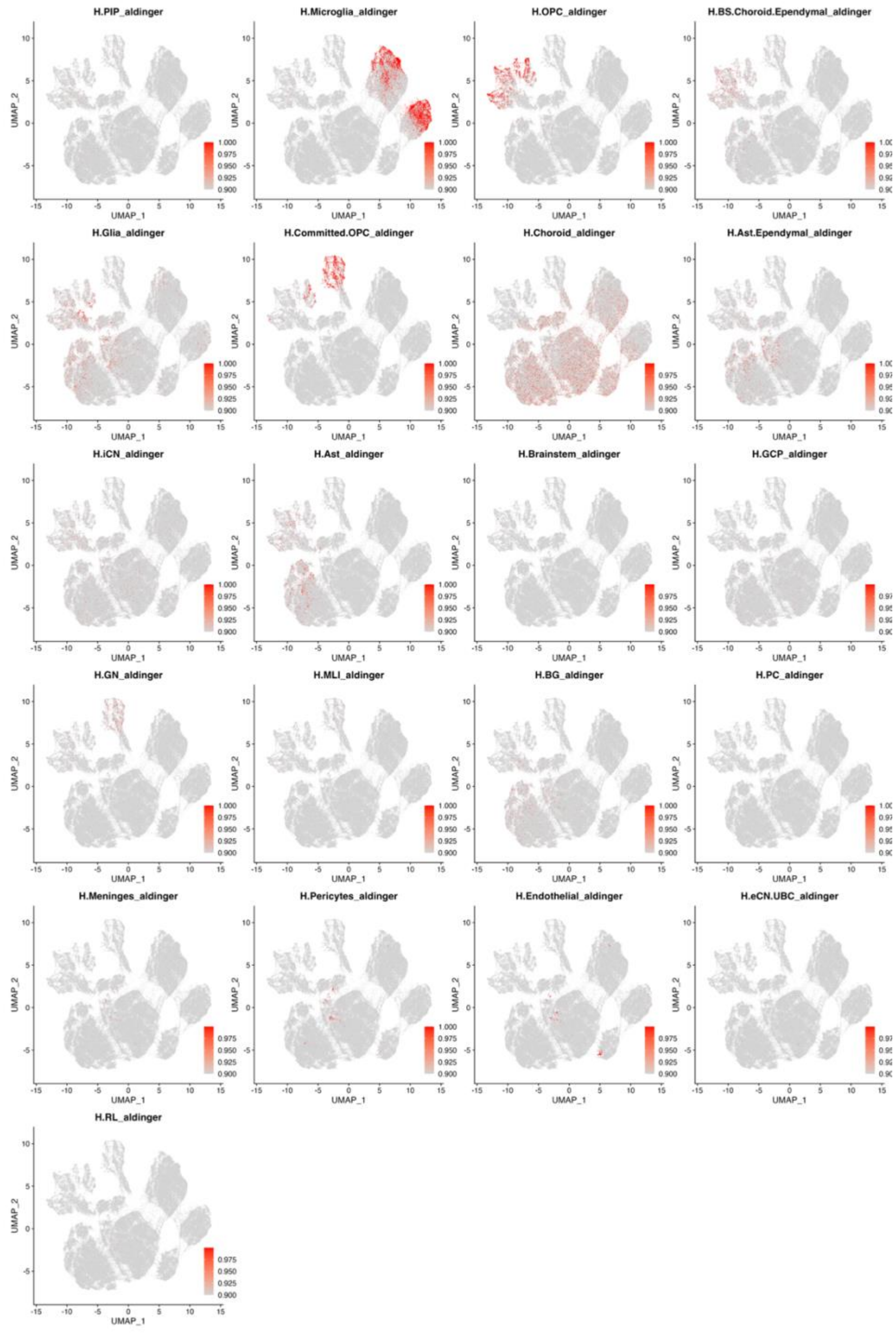

**Extended Data Figure S13. Neural Network Model Score Plot for Aldinger et al.**[2] **Cell Types.** Neural Network model score plot for Aldinger et al.[2] cell types on glioma SCRAM UMAP.

**Extended Data Figure S14**

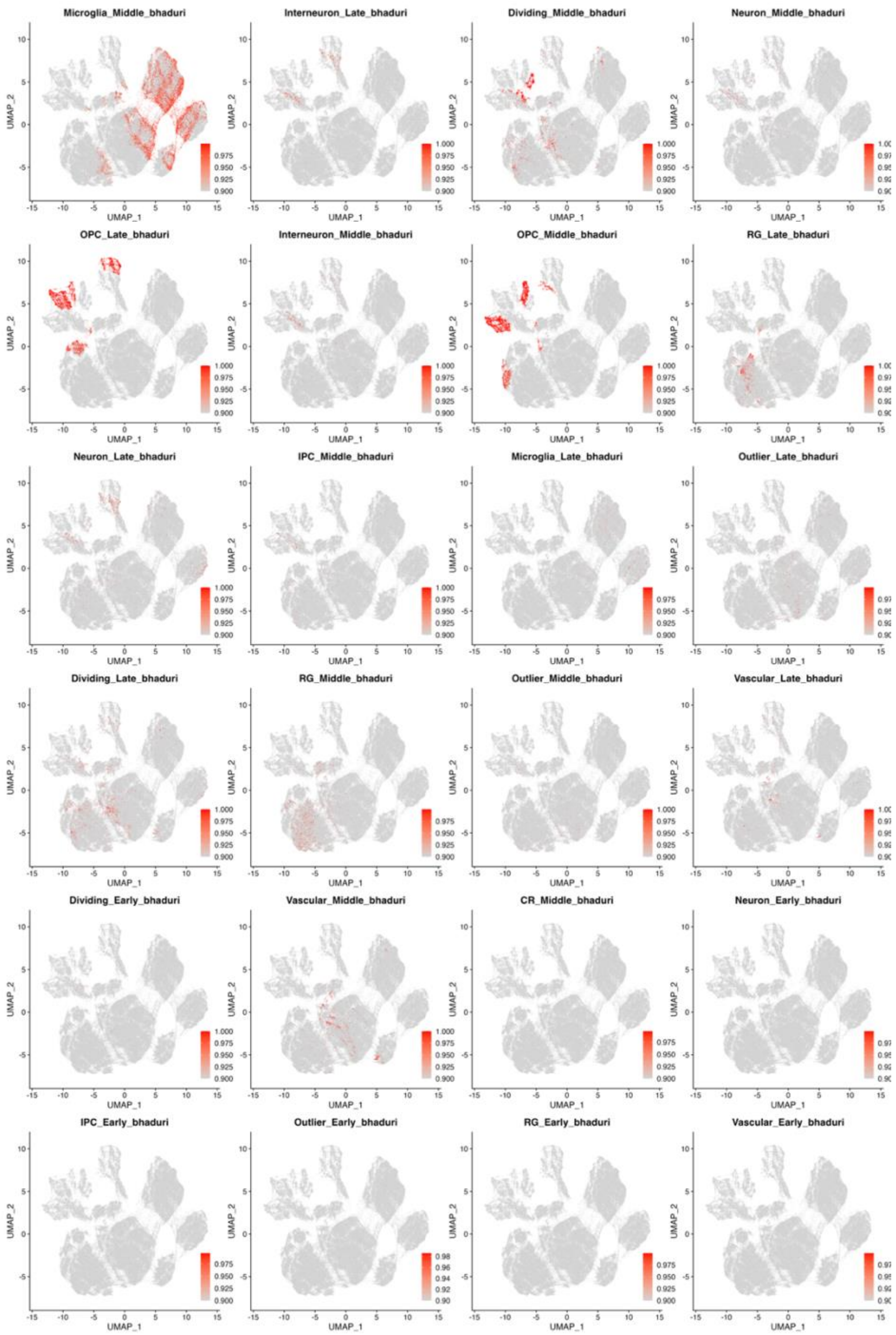

**Extended Data Figure S14. Neural Network Model Score Plot for Bhaduri et al.[3] Cell Types.** Neural Network Model score plot for Bhaduri et al.[3] cell types on glioma SCRAM UMAP.

#### Extended Data Figure S15

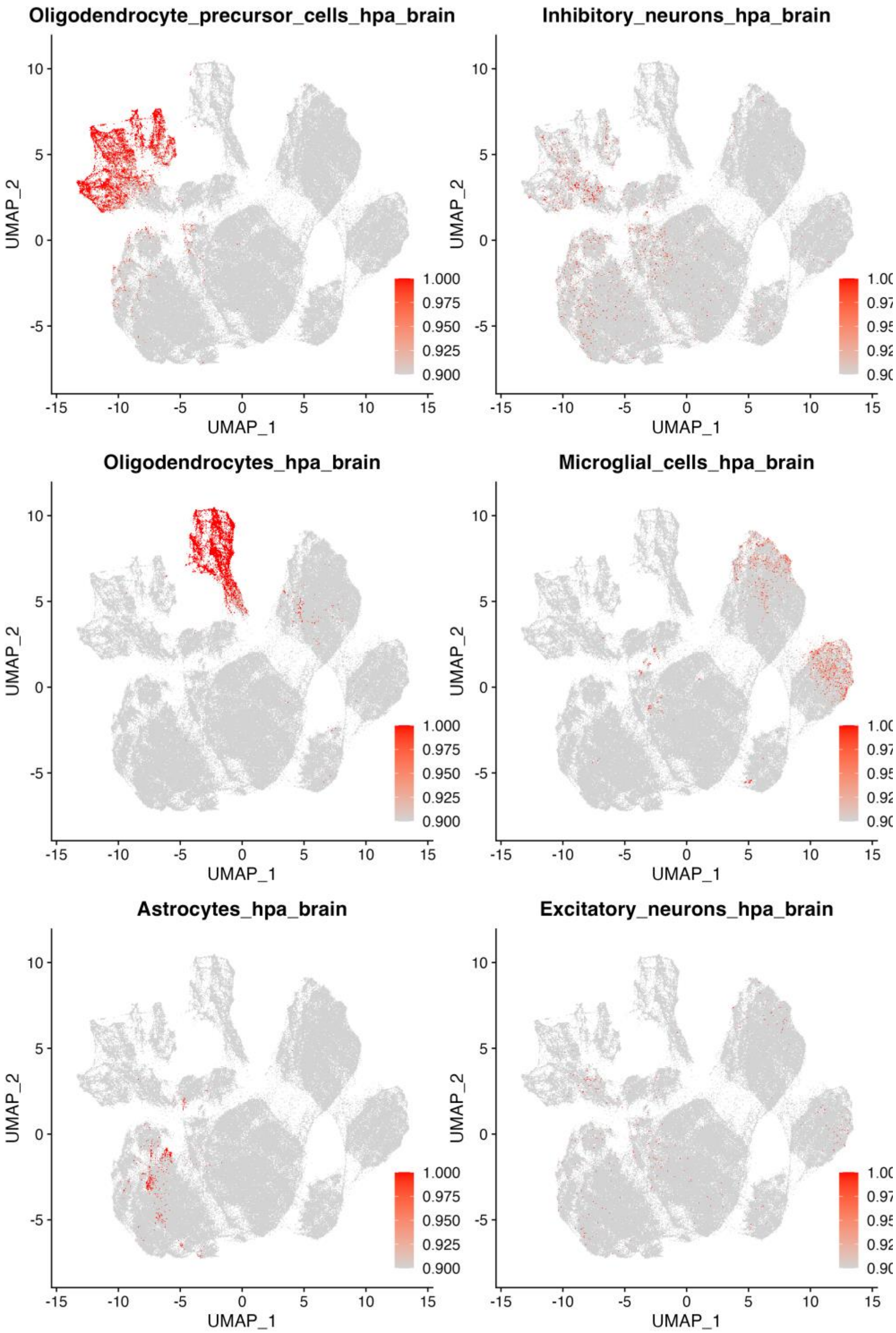

**Extended Data Figure S15. Neural Network Model Score Plot for (HPA)-Brain[4] Cell Types.** Model score plot for Human Protein Atlas (HPA)-Brain[4] cell types on glioma SCRAM UMAP.

**Extended Data Figure S16**

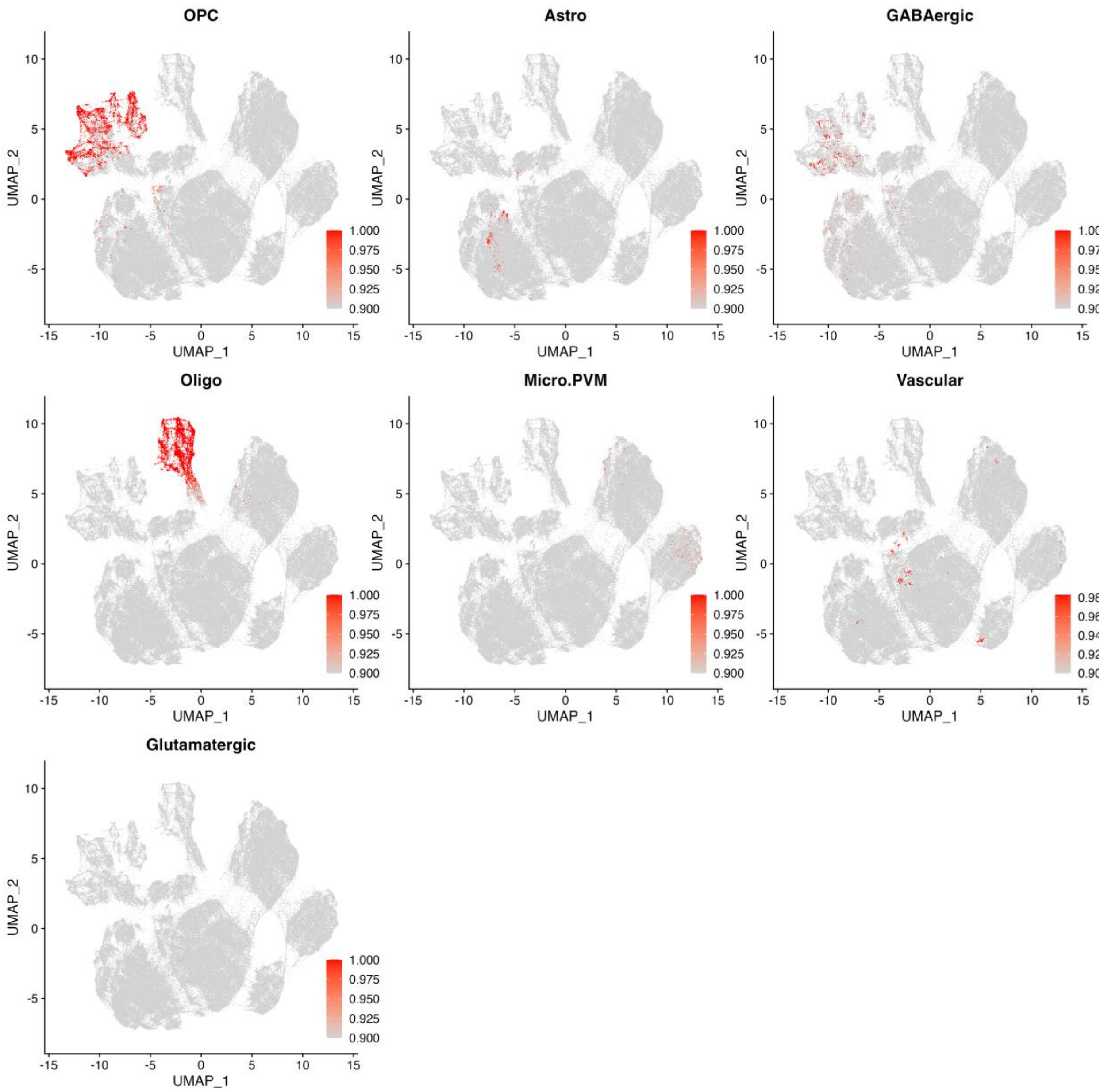

**Extended Data Figure S16. Neural Network Model Score Plot for Allen Brain Atlas[5] Cell Types.** Neural Network model score plot for Allen Brain Atlas[5] cell types on glioma SCRAM UMAP.

**Extended Data Figure S17**

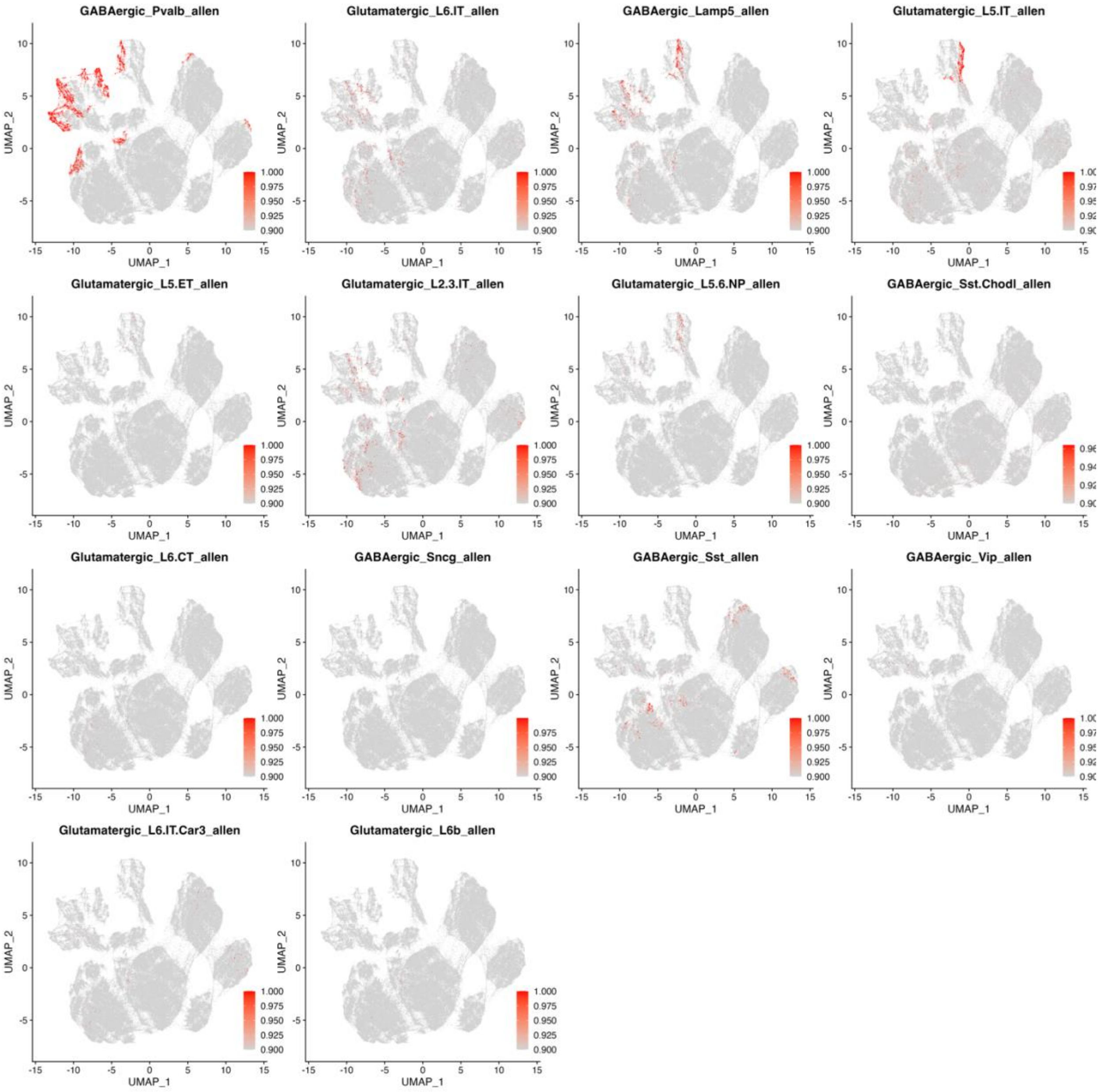

**Extended Data Figure S17. Neural Network Model Score Plot for Allen Brain Atlas[5] Neuron Cell Types.** Neural Network model score plot for Allen Brain Atlas[5] neuron cell types on glioma SCRAM UMAP.

#### Extended Data Figure S18

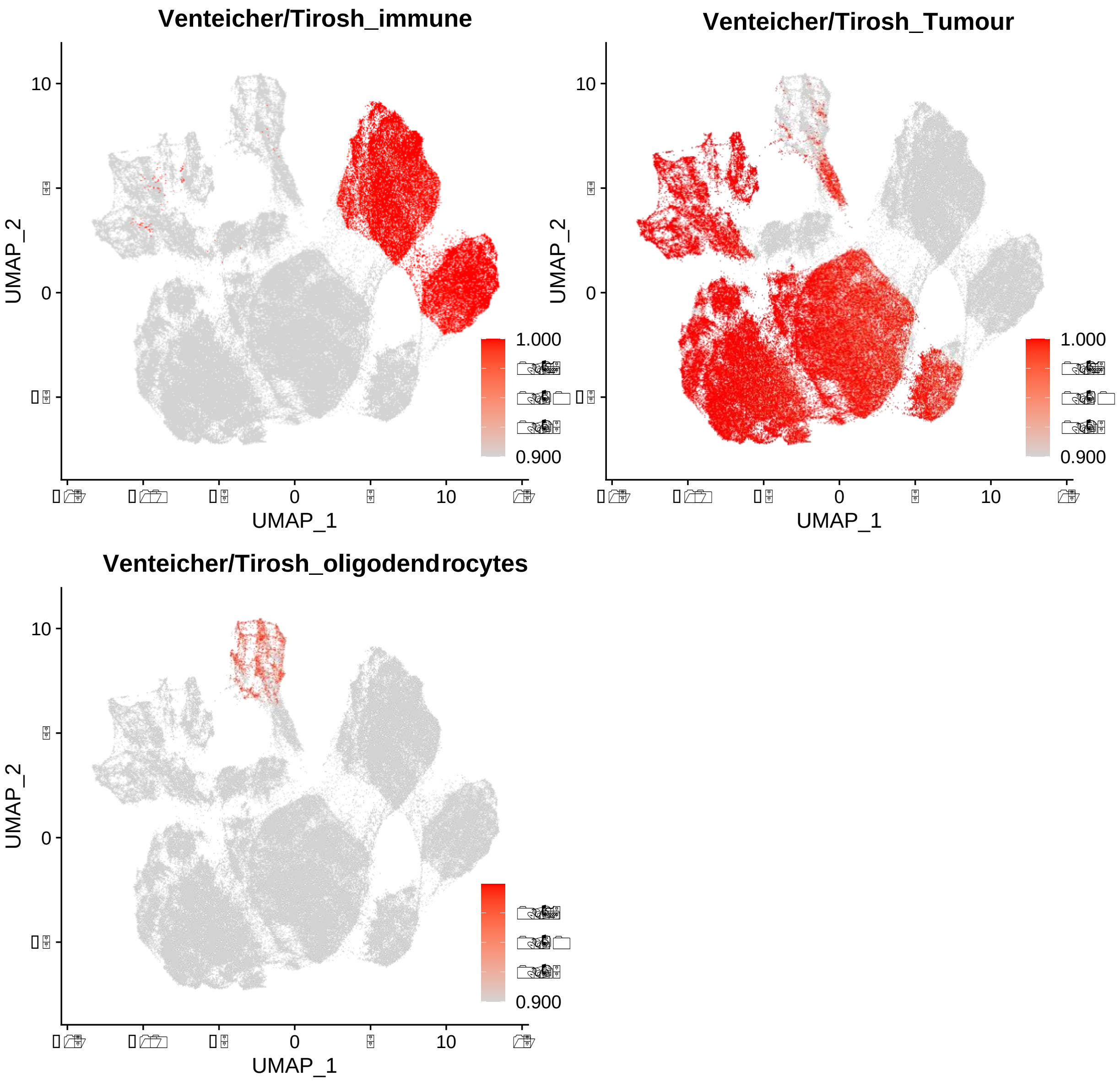

**Extended Data Figure S18. Neural Network Model Score Plot for Venteicher et al. and Tirosh et al Cell Types.** Neural Network model score plot for Venteicher et al. and Tirosh et al. cell types on glioma SCRAM UMAP [6,7].

#### Extended Data Figure S19

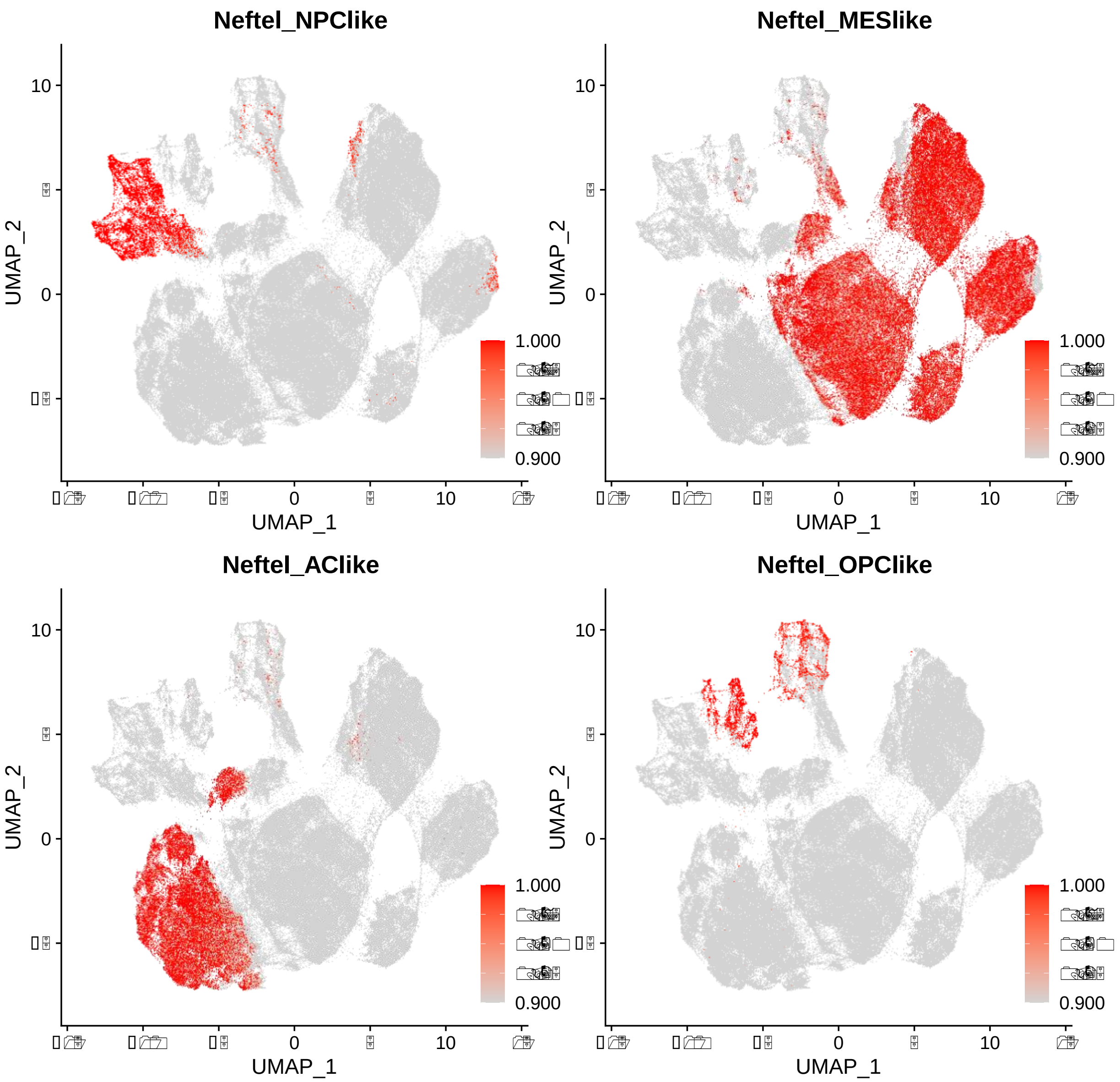

**Extended Data Figure S19. Neural Network Model Score Plot for Neftel et al. Cell Types.** Neural Network model score plot for Neftel et al.[8] cell types on our glioma SCRAM UMAP.

### Extended Data Figure S20.

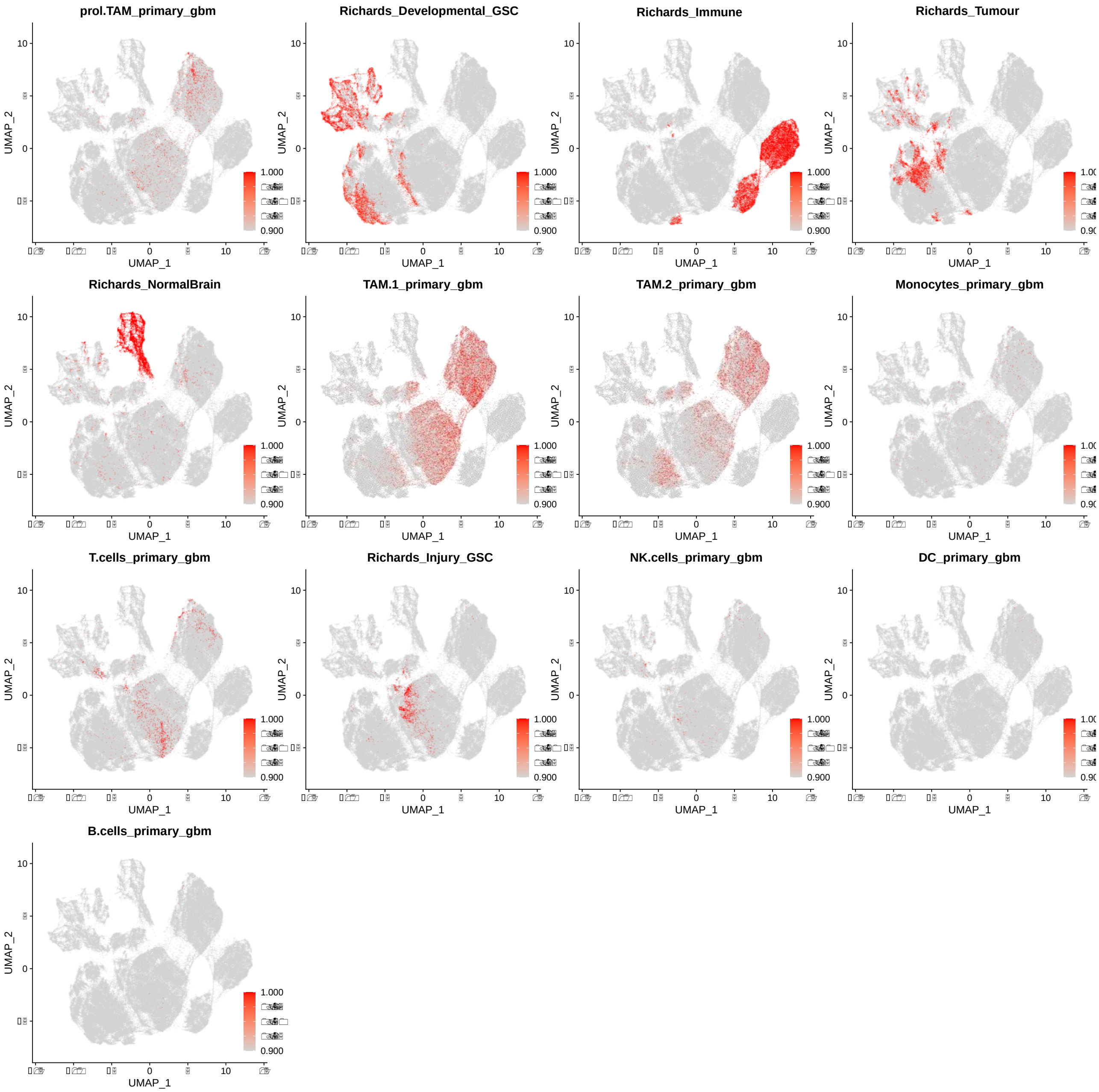

**Extended Data Figure S20. Neural Network Model Score Plot for Richard et al.[9] and GBM Immune Atlas[10] Cell Types.** Neural Network model for Richard et al.[9] and GBM Immune Atlas[10] cell types on glioma SCRAM UMAP.

#### Extended Data Figure S21

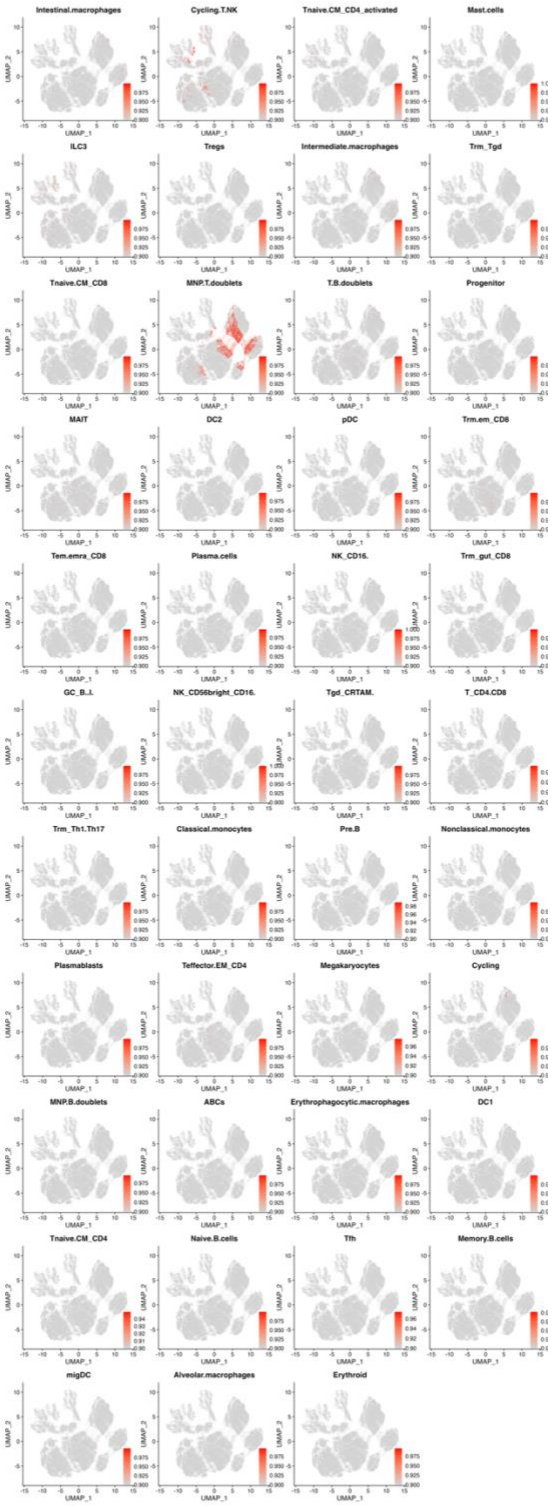

**Extended Data Figure S21. Neural Network Model Score Plot for Tissue Immune Atlas[11] Cell Types.** Neural Network model score plot for Tissue Immune Atlas[11] cell types on glioma SCRAM UMAP.

**Extended Data Figure S22**

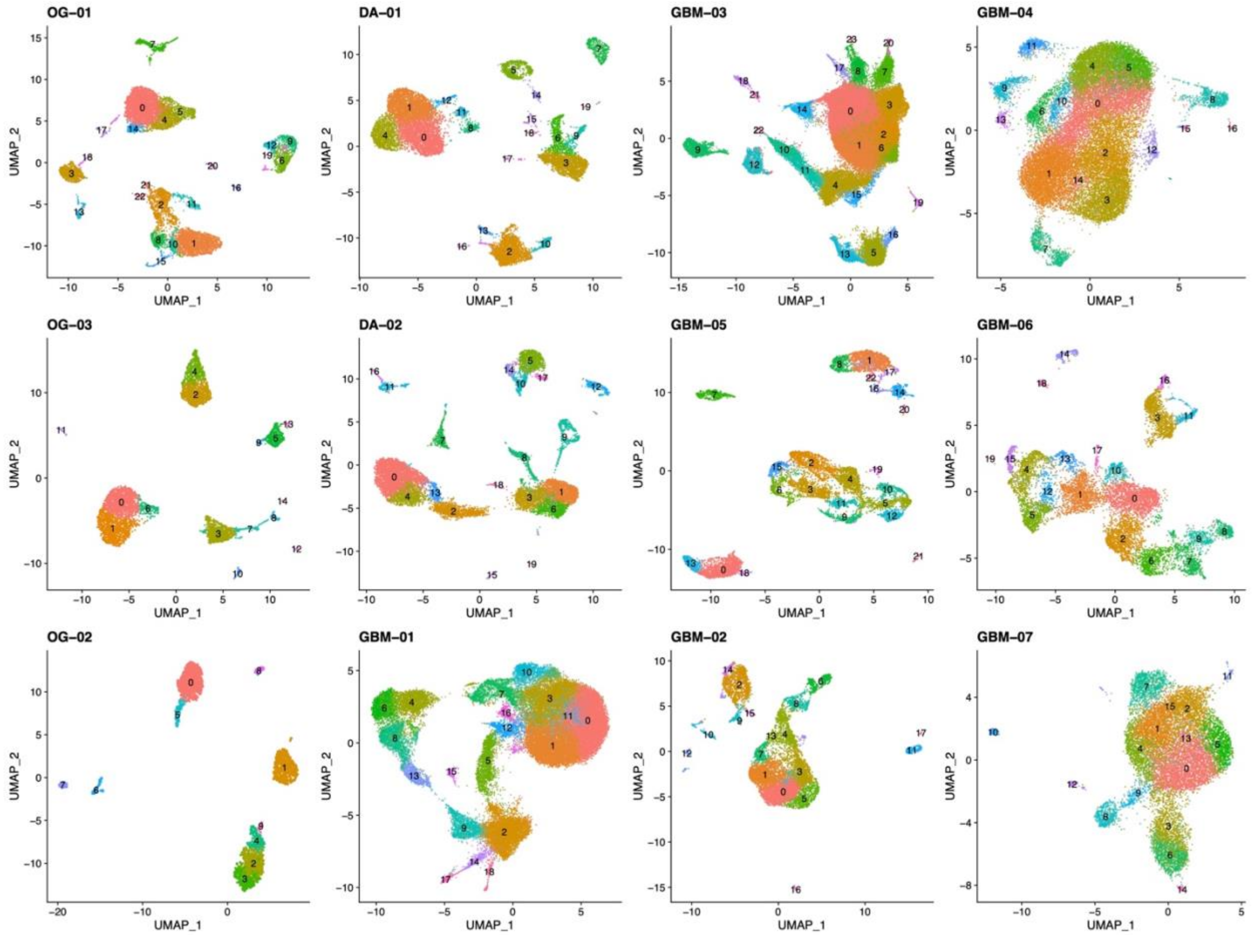

**Extended Data Figure S22. Per-patient Seurat UMAP and Clusters.**

**Extended Data Figure S23**

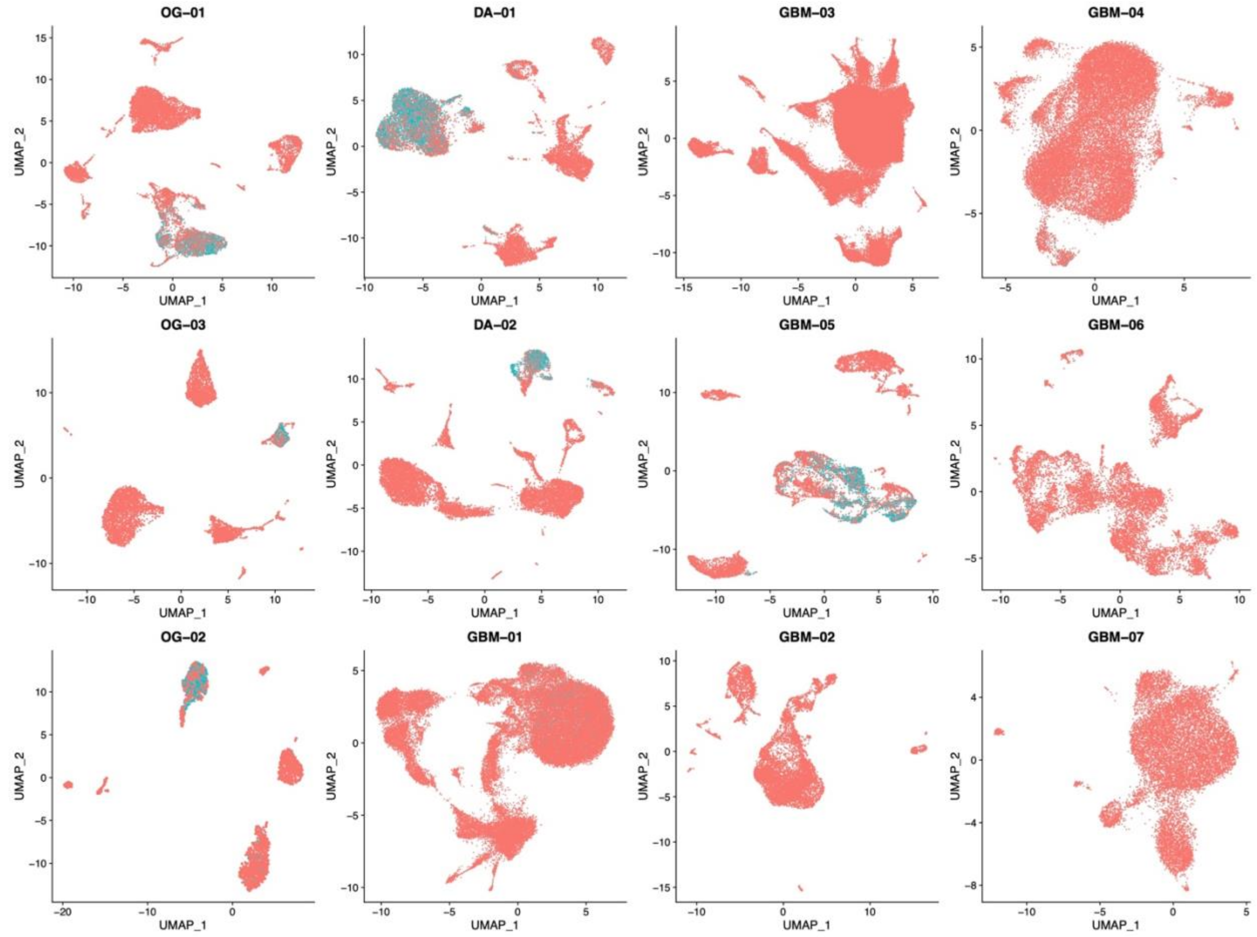

**Extended Data Figure S23. GABA-OPC Tumor Cell Annotations.** GABA-OPC tumor cells are shown as blue dots on Seurat based UMAP.

**Extended Data Figure S24**

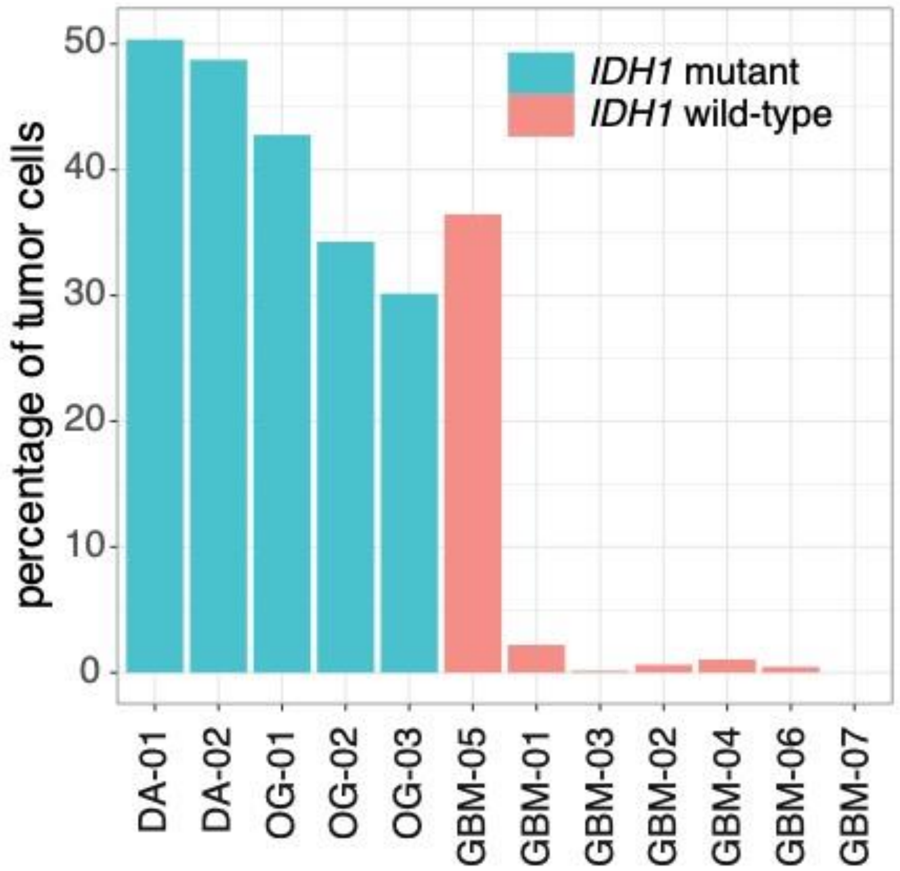

**Extended Data Figure S24. GABA-OPC Tumor Cell Percentages.** GABA-OPC tumor cell percentage barplot across all samples.

**Extended Data Figure S25**

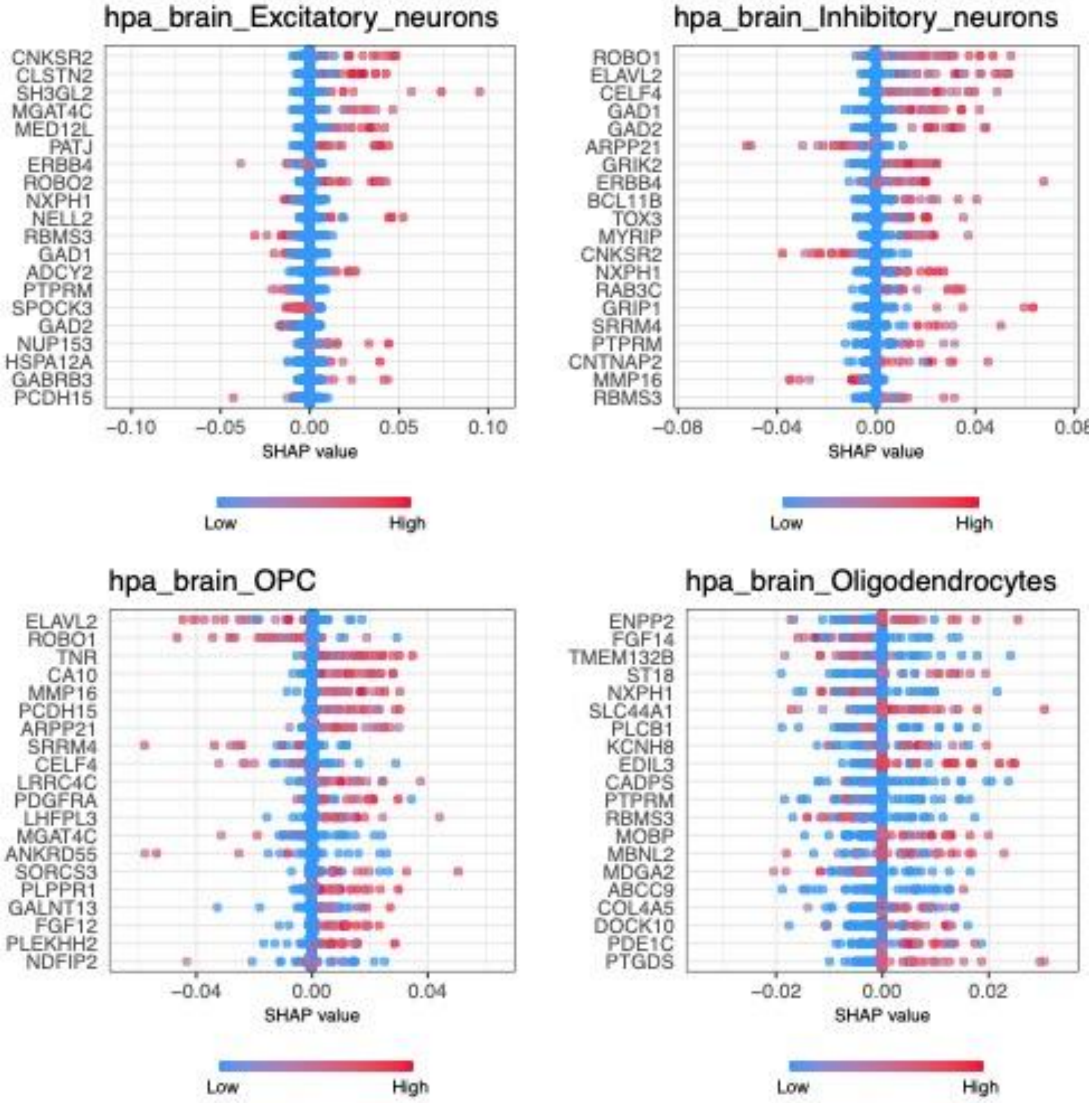

**Extended Data Figure S25. SHAP Analysis for Cell Types in the Human Protein Atlas (HPA) Brain**. The figure displays the results of SHapley Additive exPlanations (SHAP) analysis, showing the top impactful genes from each cell type in our training dataset, Human Protein Atlas (HPA)-Brain[4]. The SHAP impact score plot provides insight into the contribution scores of individual genes in our predictive model, specifically in our glioma single-cell data.

**Extended Data Figure S26.**

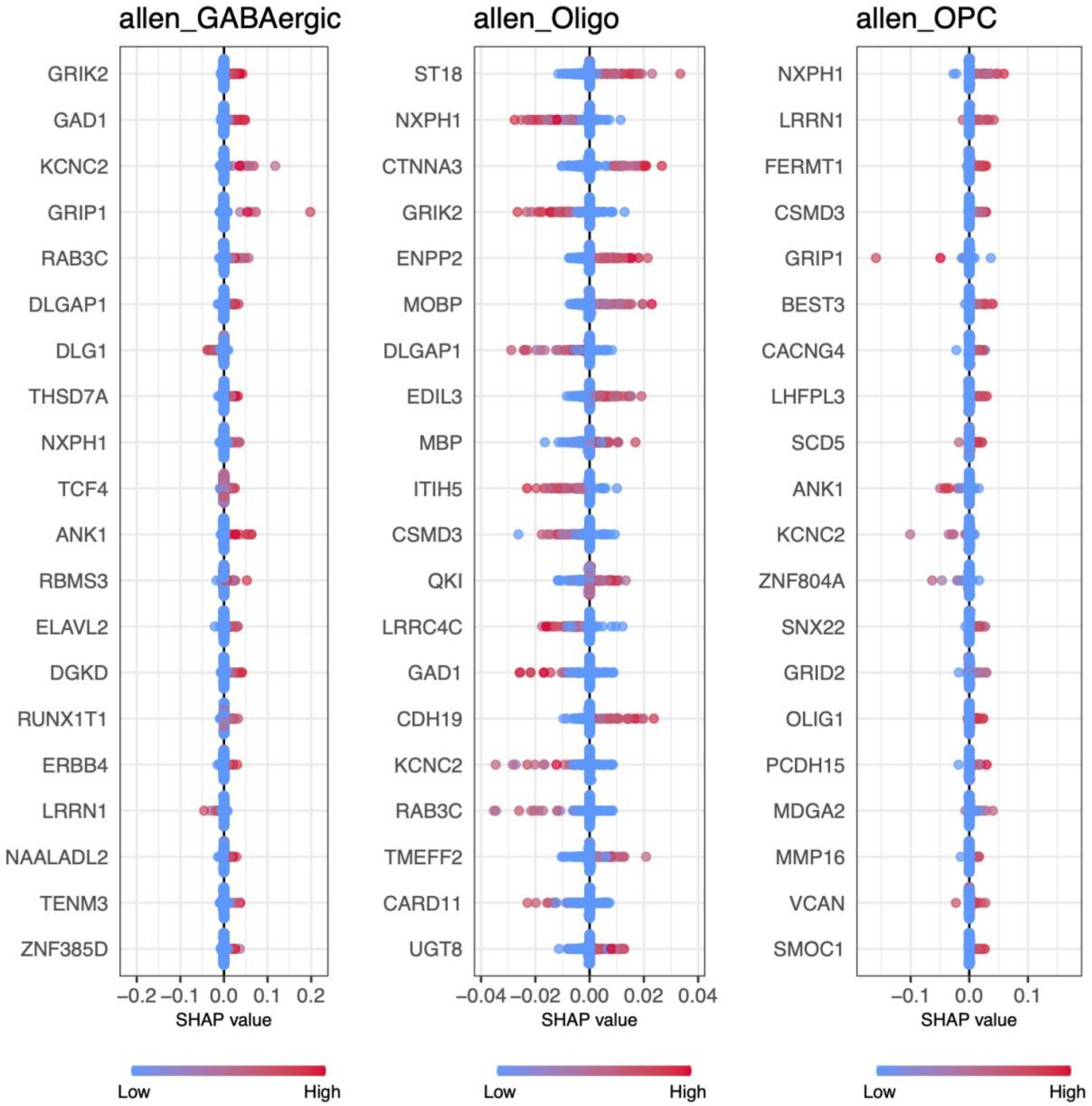

**Extended Data Figure S26. SHAP Analysis for Neuron Cell Types in the Allen Brain Atlas.** The figure displays the results of SHAP analysis, showing the top impactful genes from each cell type in our training dataset, Allen Brain Atlas[5] neuron. The SHAP impact score plot provides insight into the contribution scores of individual genes in our predictive model, specifically in our glioma single-cell data.

**Extended Data Figure S27**

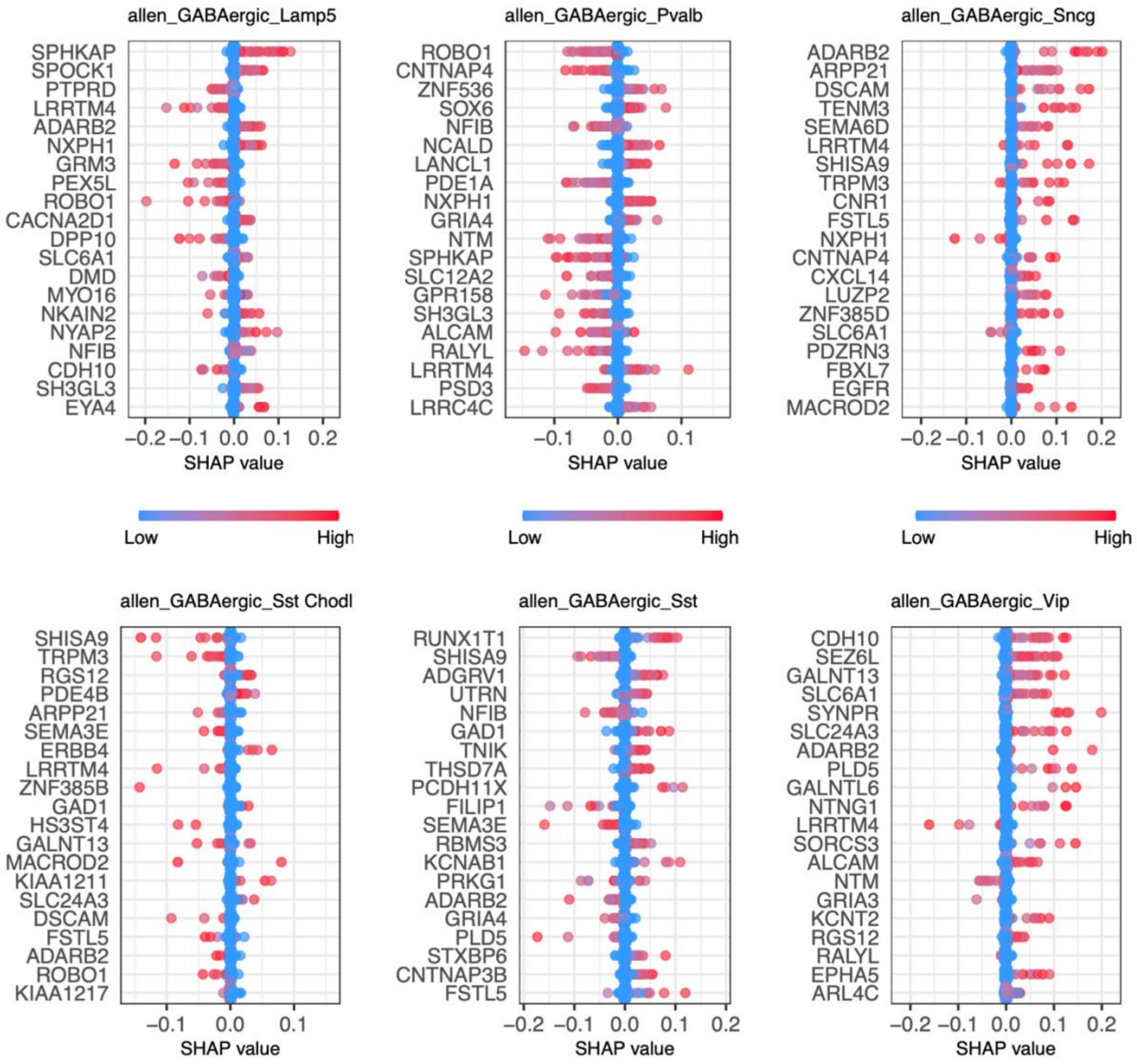

**Extended Data Figure S27. SHAP Analysis for Cell Types in the Allen Brain Atlas.** The figure displays the results of SHAP analysis, showing the top impactful genes from each cell type in our training dataset, Allen Brain Atlas[5]. The SHAP impact score plot provides insight into the contribution scores of individual genes in our predictive model, specifically in our glioma single-cell data.

**Extended Data Figure S28.**

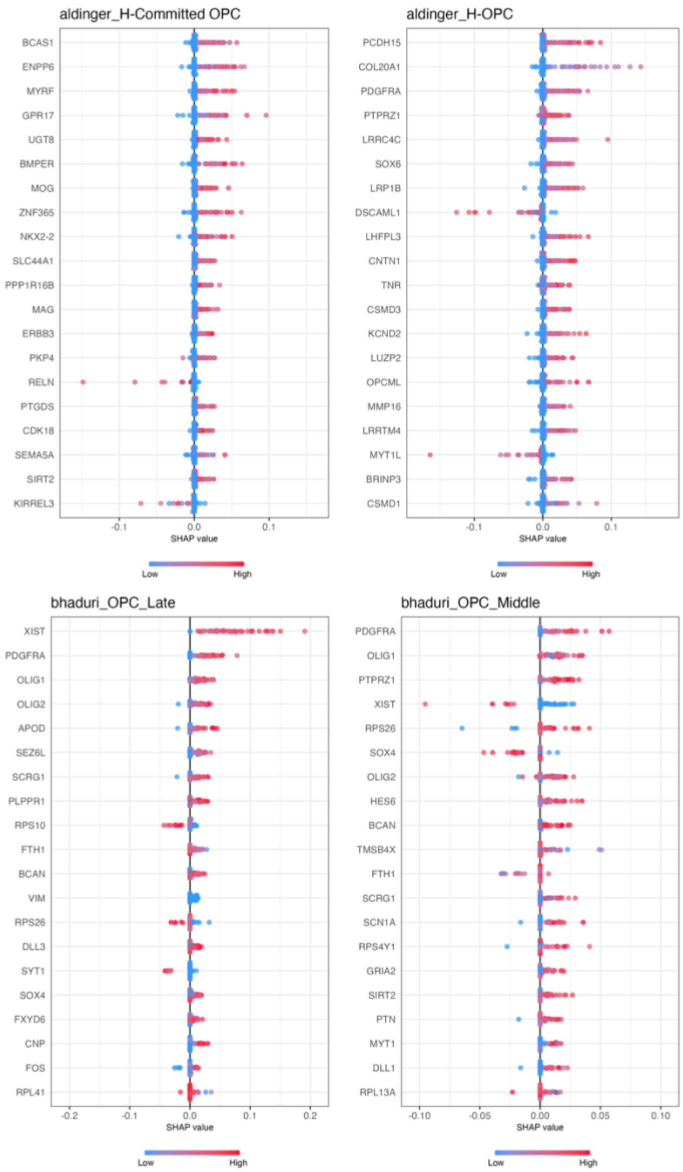

**Extended Data Figure S28. SHAP Analysis for Aldinger et al.[2] and Bhaduri et al[3] Cell Types in the Allen Brain Atlas.** The figure displays the results of SHAP analysis, showing the top impactful genes from each cell type in our training dataset, Aldinger et al.[2] and Bhaduri et al[3]. The SHAP impact score plot provides insight into the contribution scores of individual genes in our predictive model, specifically in our glioma single-cell data.

**Extended Data Figure S29**

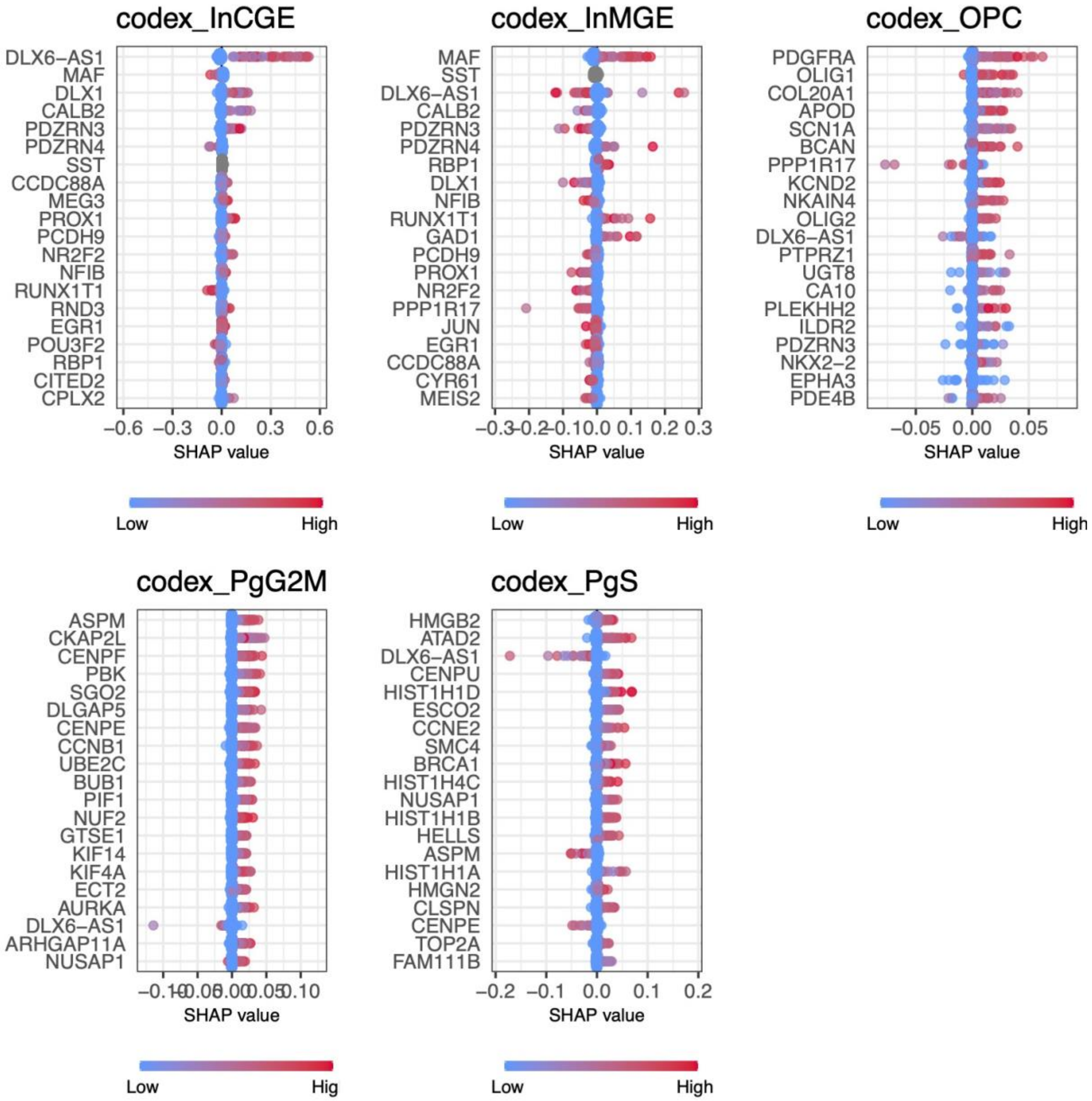

**Extended Data Figure S29. SHAP Analysis for CoDEx Cell Types in the Allen Brain Atlas.** The figure displays the results of SHAP analysis, showing the top impactful genes from each cell type in our training dataset, CoDEx[1]. The SHAP impact score plot provides insight into the contribution scores of individual genes in our predictive model, specifically in our glioma single-cell data.

**Extended Data Figure S30**

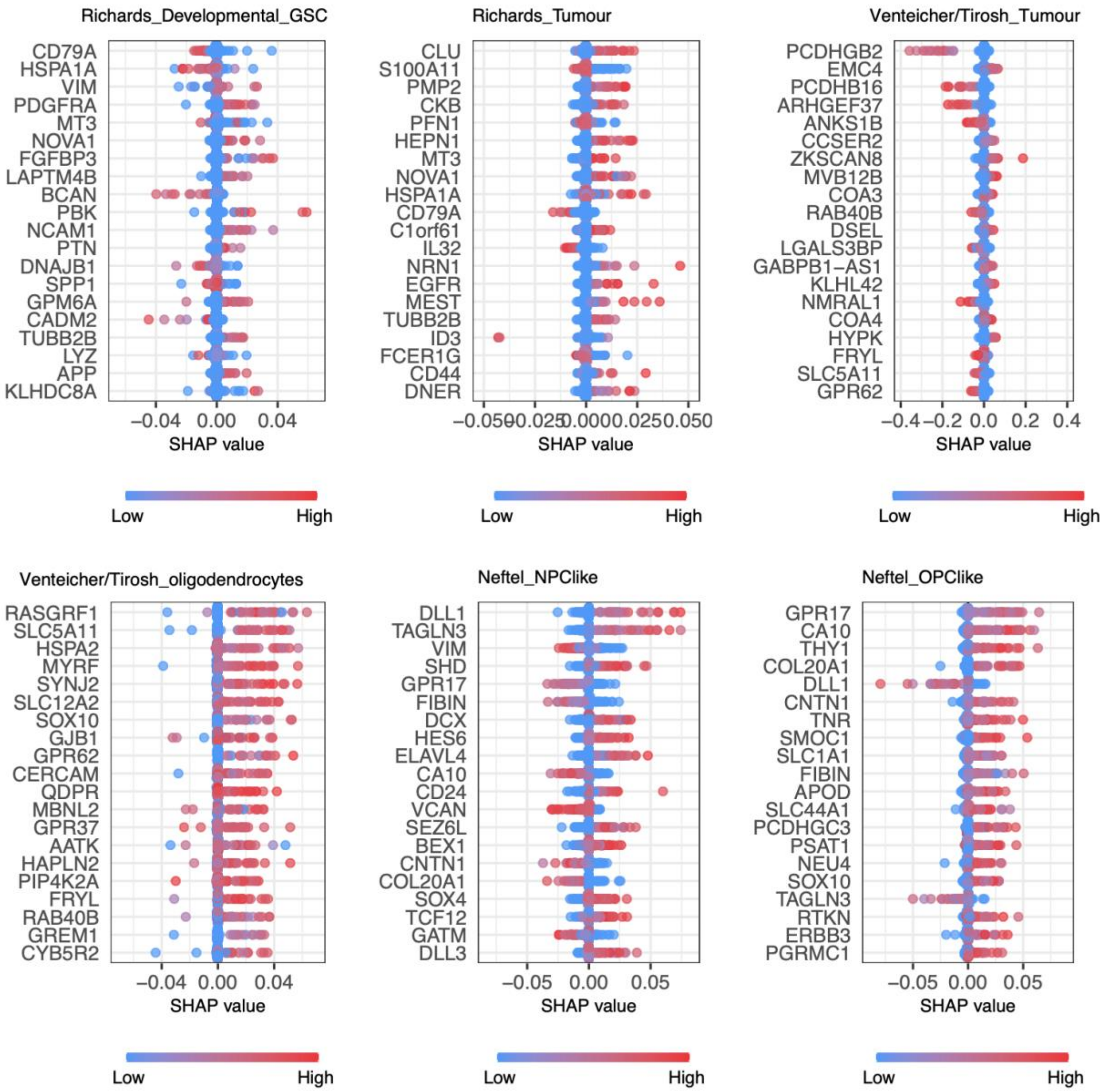

**Extended Data Figure S30. SHAP Analysis Neftel et al., Richards et al, Venteicher et al. and Tirosh et al Cell Types in the Allen Brain Atlas** The figure displays the results of SHAP analysis, showing the top impactful genes from each cell type in our training dataset, Neftel et al., Richards et al, Venteicher et al. and Tirosh et al datasets [6–9]. The SHAP impact score plot provides insight into the contribution scores of individual genes in our predictive model, specifically in our glioma single-cell data.

#### Extended Data Figure S31

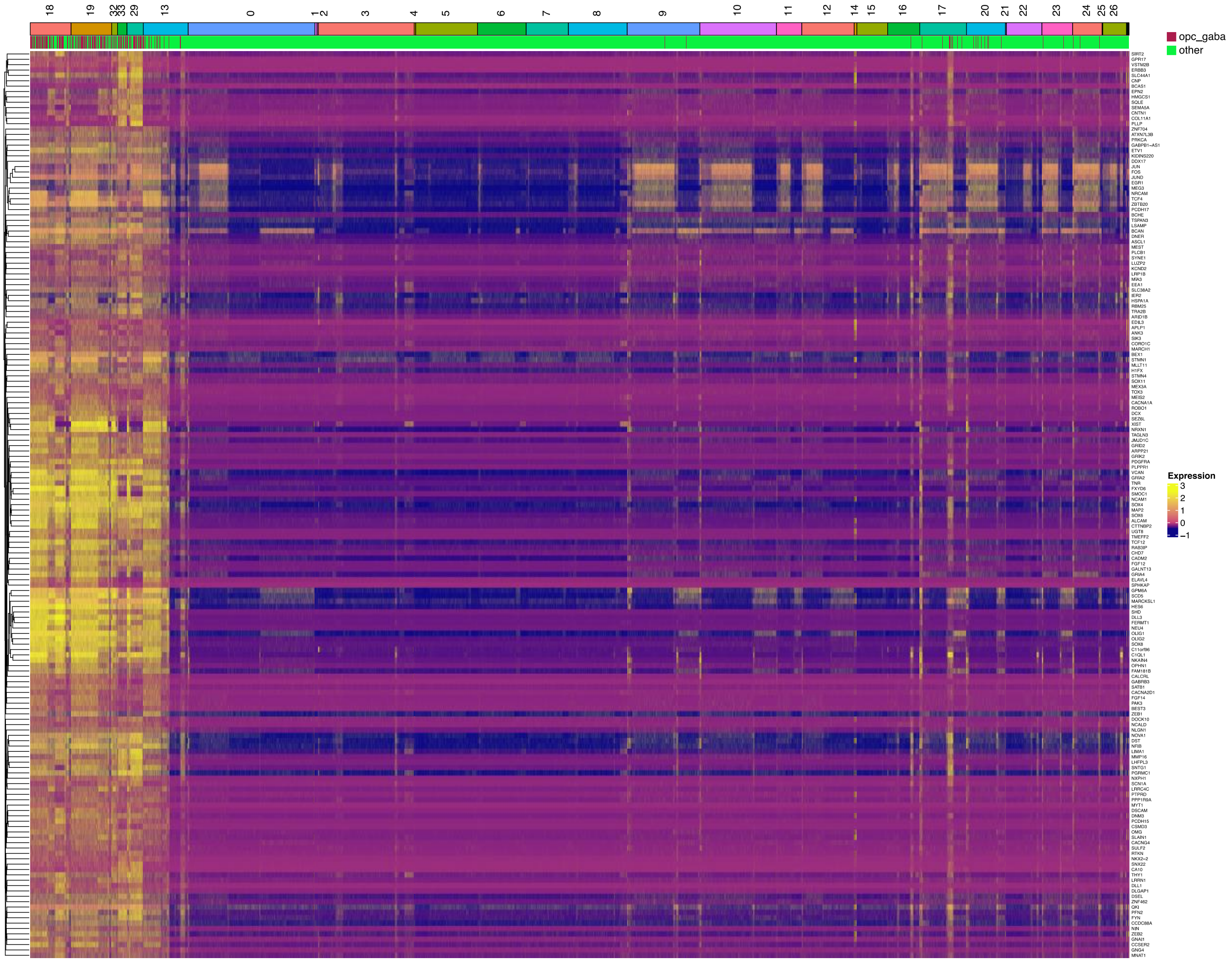

**Extended Data Figure S31. Heatmap of GABA-OPC Signature Genes in Tumor Cells.** The heatmap illustrates the expression of GABA-OPC signature genes in a glioma tumor cells, derived through the analysis of overlapping differentially expressed genes (DEGs) and SHAP (Shapley Additive exPlanations) analysis. GABA-OPC tumor cells and SCRAM UMAP clusters are shown on top of the heatmap.

#### Extended Data Figure S32

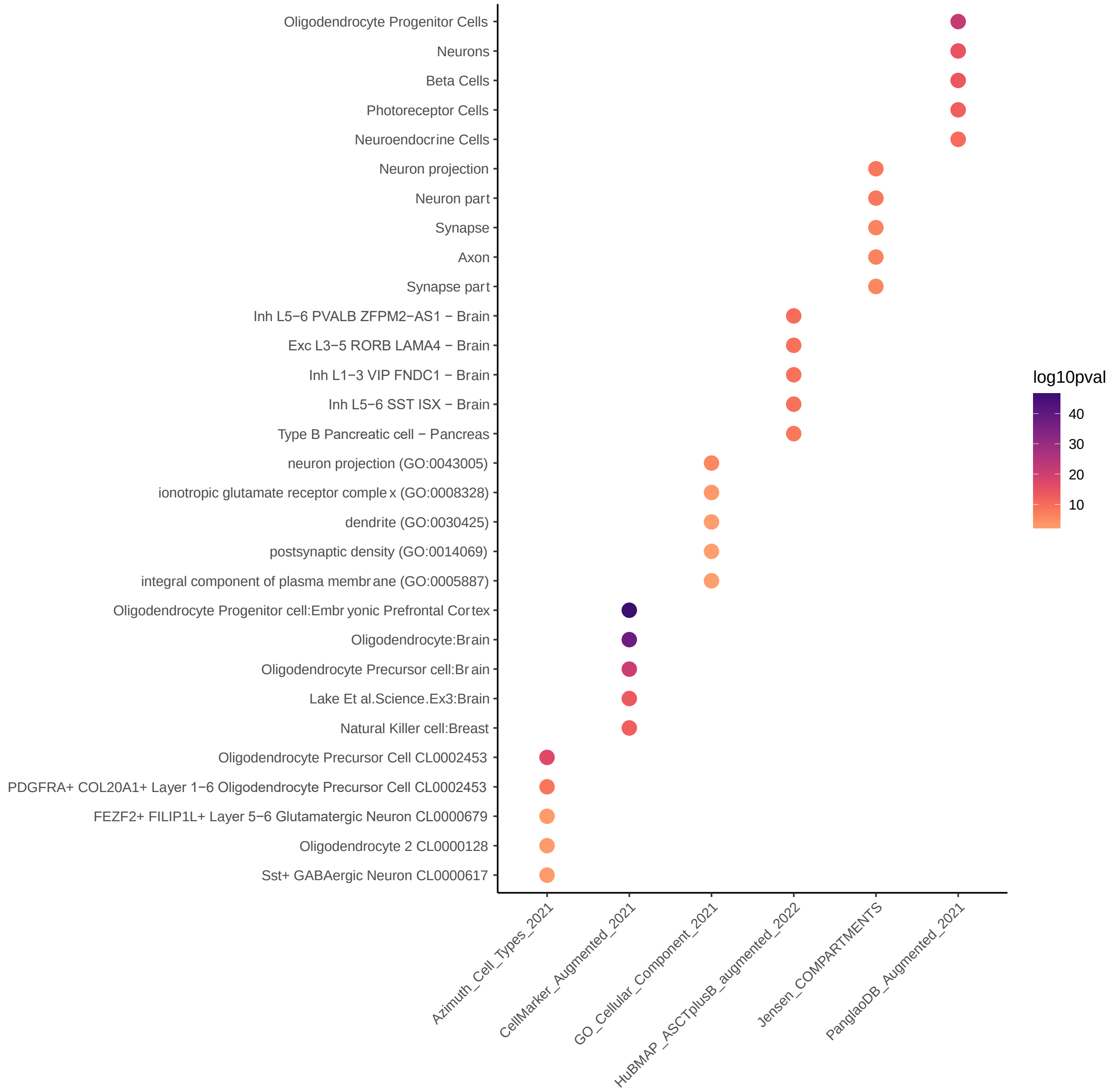

**Extended Data Figure S32. Enrichment Analysis of GABA-OPC Signature Genes.** Enrichment analysis of GABA-OPC signature genes derived from the overlapping analysis of DEGs and SHAP analysis.

#### Extended Data Figure S33

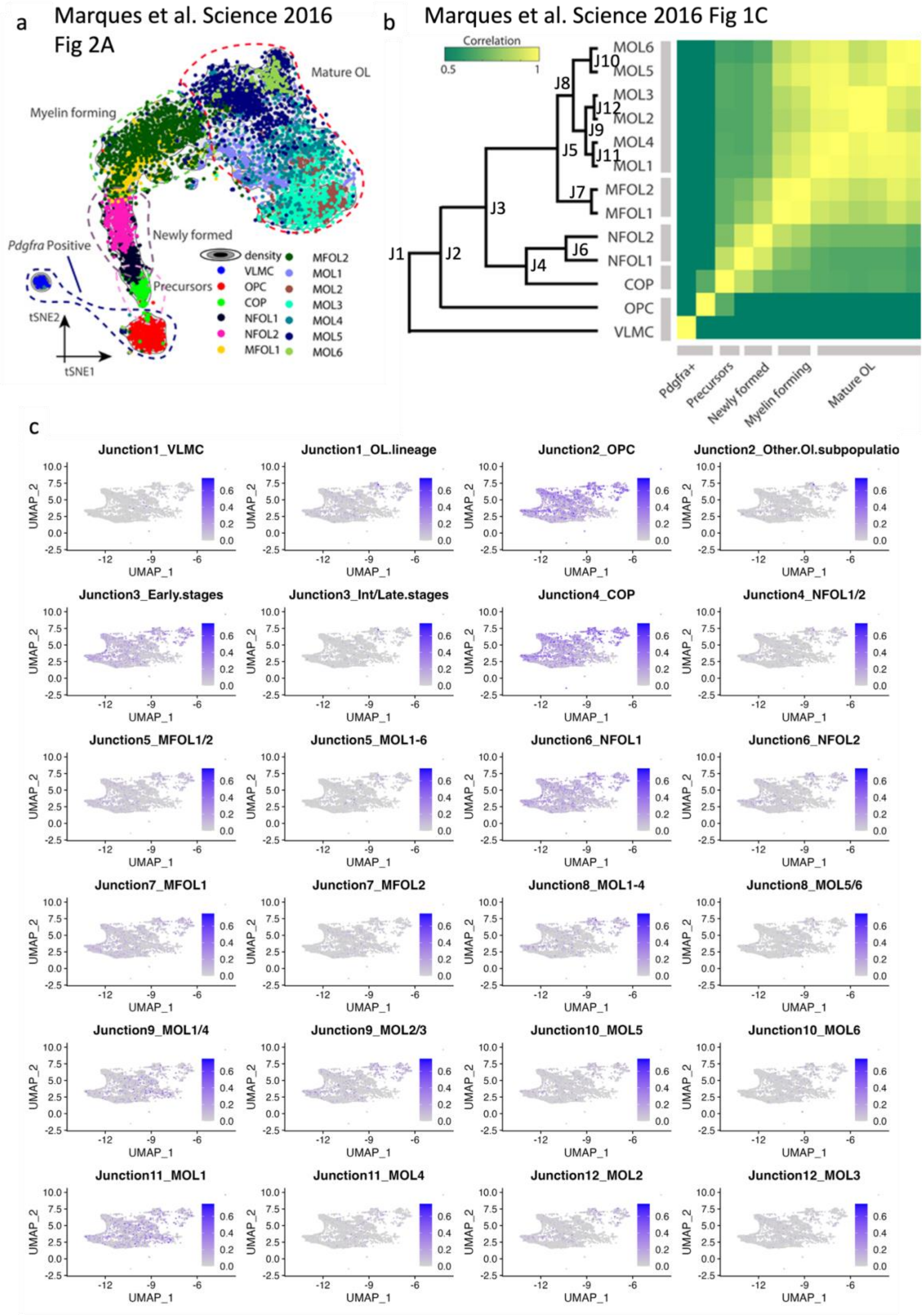

**Extended Data Figure S33. Characterizing Oligodendrocyte Lineage Classes of GABA-OPC Gene Set (**a) t-SNE projection showing the trajectory from OPCs to mature oligodendrocytes in Marques et al.[12] paper. (b) Hierarchical clustering of expression correlation matrix of oligodendrocyte lineage classes from Marques et al. Science 2016 paper. (c) Feature Plot for the oligodendrocyte lineage classes gene set scoring in our GABA-OPC tumor cells.

**Extended Data Figure S34**

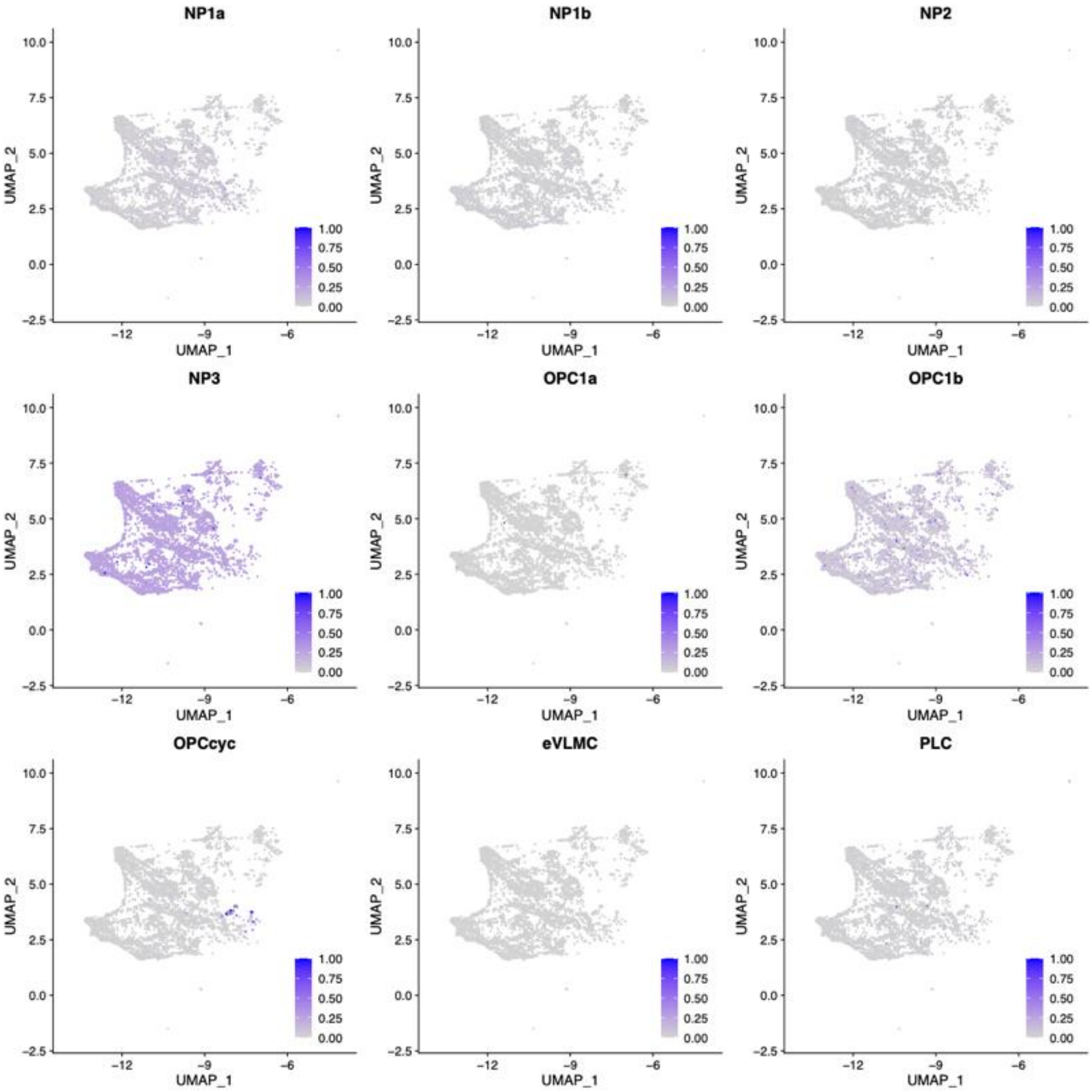

**Extended Data Figure S34 Characterizing Oligodendrocyte Lineage Classes of GABA-OPC Gene Set.** Feature Plot for the Marques et al.[13] classes gene set scoring in our GABA-OPC tumor cells.

### Extended Data Figure S35

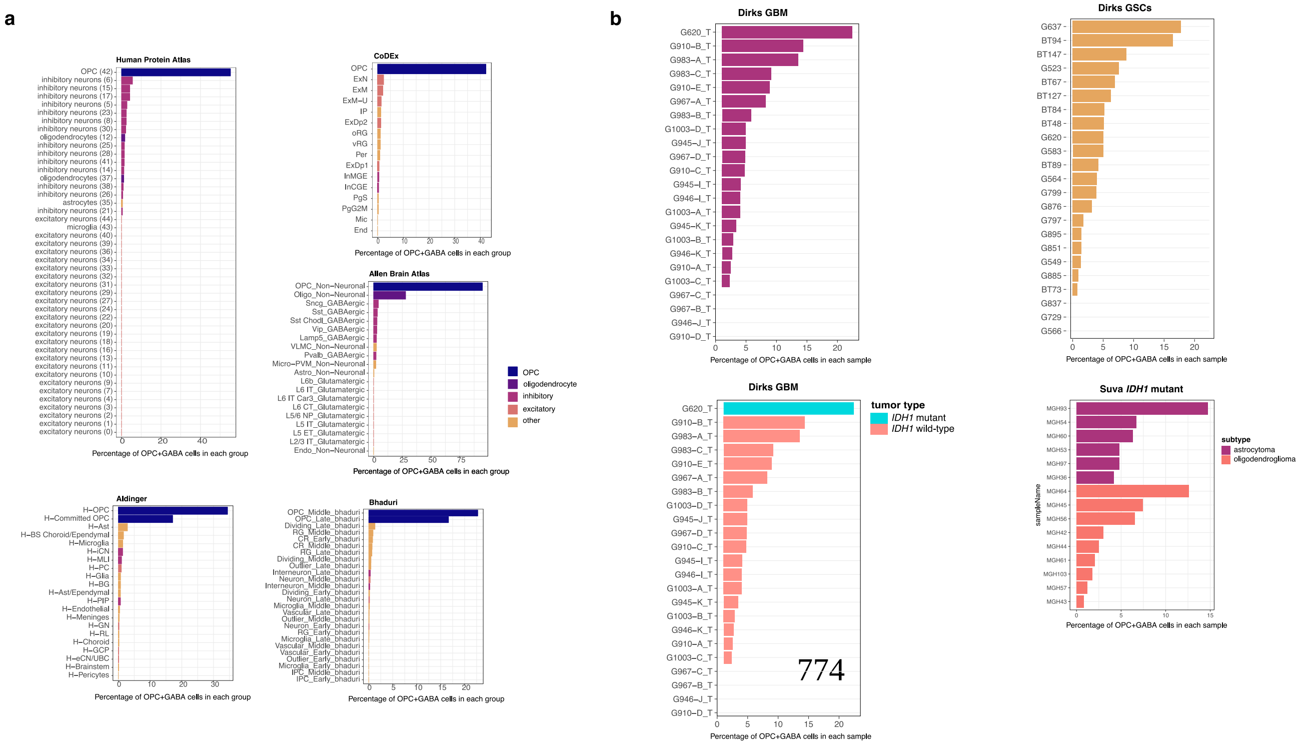

**Extended Data Figure S35. Quantifying the Percentage of GABA-OPC Cells in Various Cell Types and Tumor Cells.** (a) Percentage of GABA-OPC cells among different cell types (b) Percentage of GABA-OPC cells among tumor cells.

**Extended Data Figure S36**

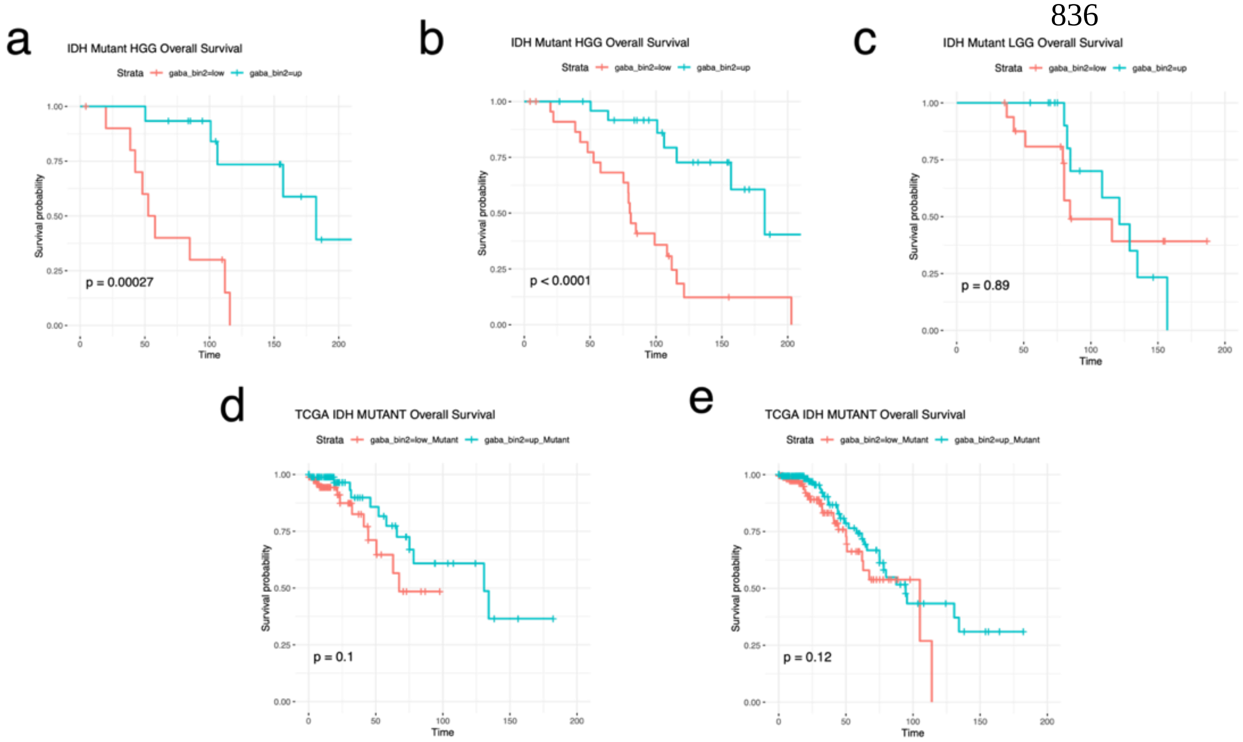

**Extended Data Figure S36. Survival analyses.** (a) Survival plot for in-house IDH1 mutant HGG samples split into low and high GABA-OPC groups based on median scoring. (b) Survival plot for in-house IDH1 mutant HGG samples split into low and high GABA-OPC groups based on upper and lower quartile scoring. Samples in the middle are removed. (c) Survival plot for in-house IDH1 mutant LGG samples split into low and high GABA-OPC groups based on median scoring. (d) Survival plot for TCGA IDH1 mutant samples split into low and high GABA-OPC groups based on upper and lower quartile scoring. Samples in the middle are removed. (e) Survival plot for TCGA IDH1 mutant samples split into low and high GABA-OPC groups based on median scoring.

#### Extended Data Figure S37

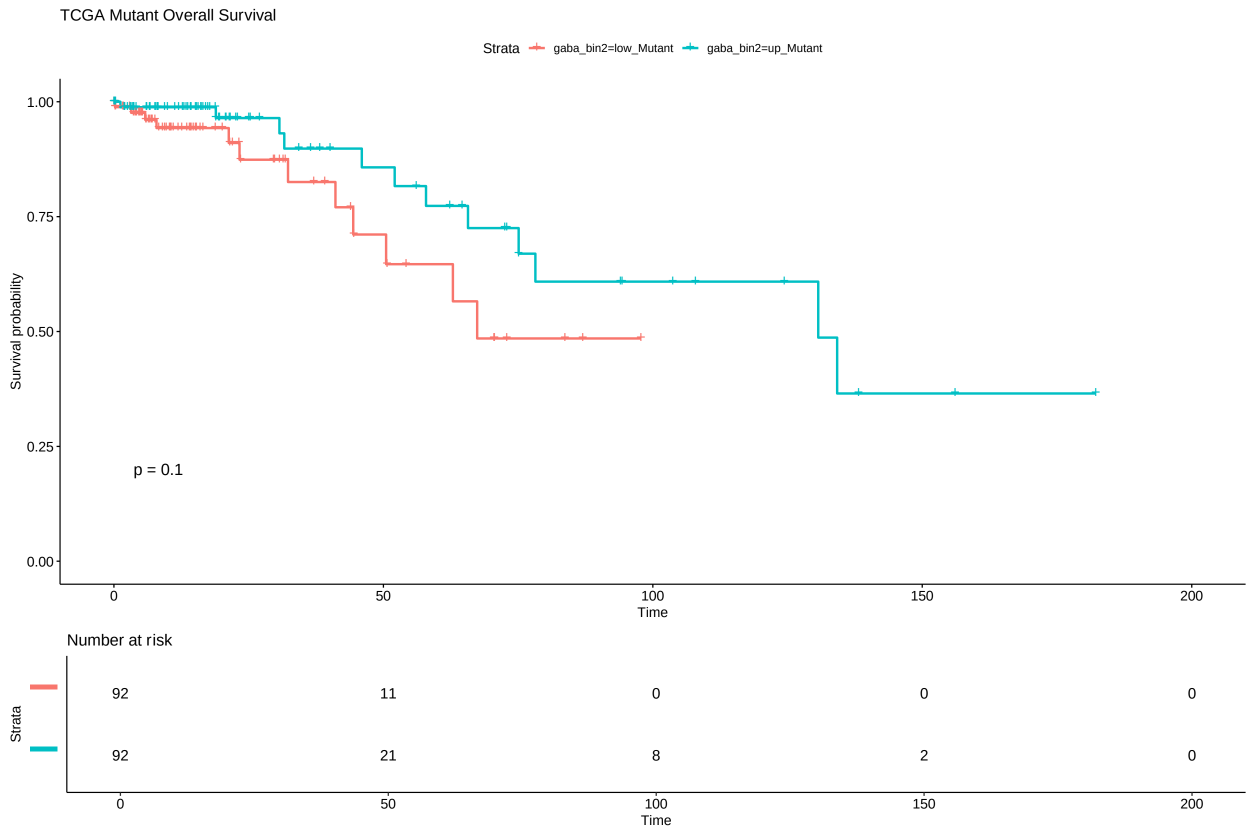

**Extended Data Figure S37. Survival analyses in IDH$^{mut}$ samples.** Survival analysis comparing IDH Mutant TCGA cohort samples with high GABA-OPC scores versus low GABA-OPC scores.

**Extended Data Figure S38**

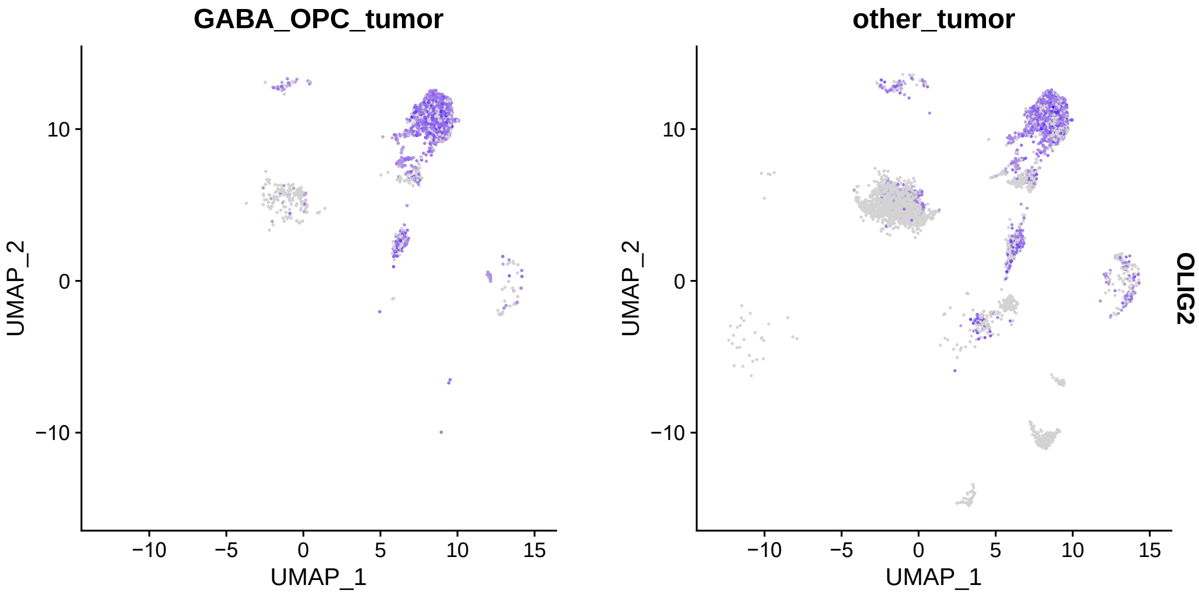

**Extended Data Figure S38. OLIG2 Expression in IDH-Mutant GABA-OPC Tumor Cells Compared to Other IDH-Mutant Tumor Cells.** The feature plot illustrates OLIG2 expression in IDH mutant tumor cells splitted by GABA-OPC tumor and other tumor cells.

#### Extended Data Figure S39

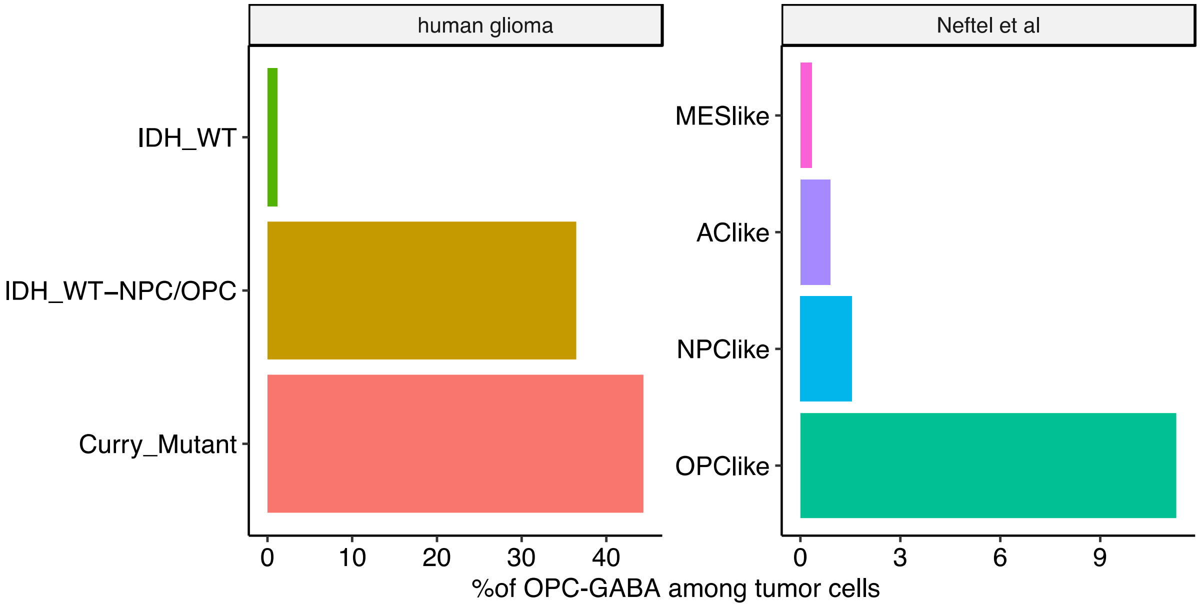

**Extended Data Figure S39. Percentage of OPC-GABA Cells Among Tumor Cells.** Percentage of OPC-GABA cells among tumor cells in in-house glioma and Neftel et al.'s IDH WT glioma datasets.

**Extended Data Figure S40**

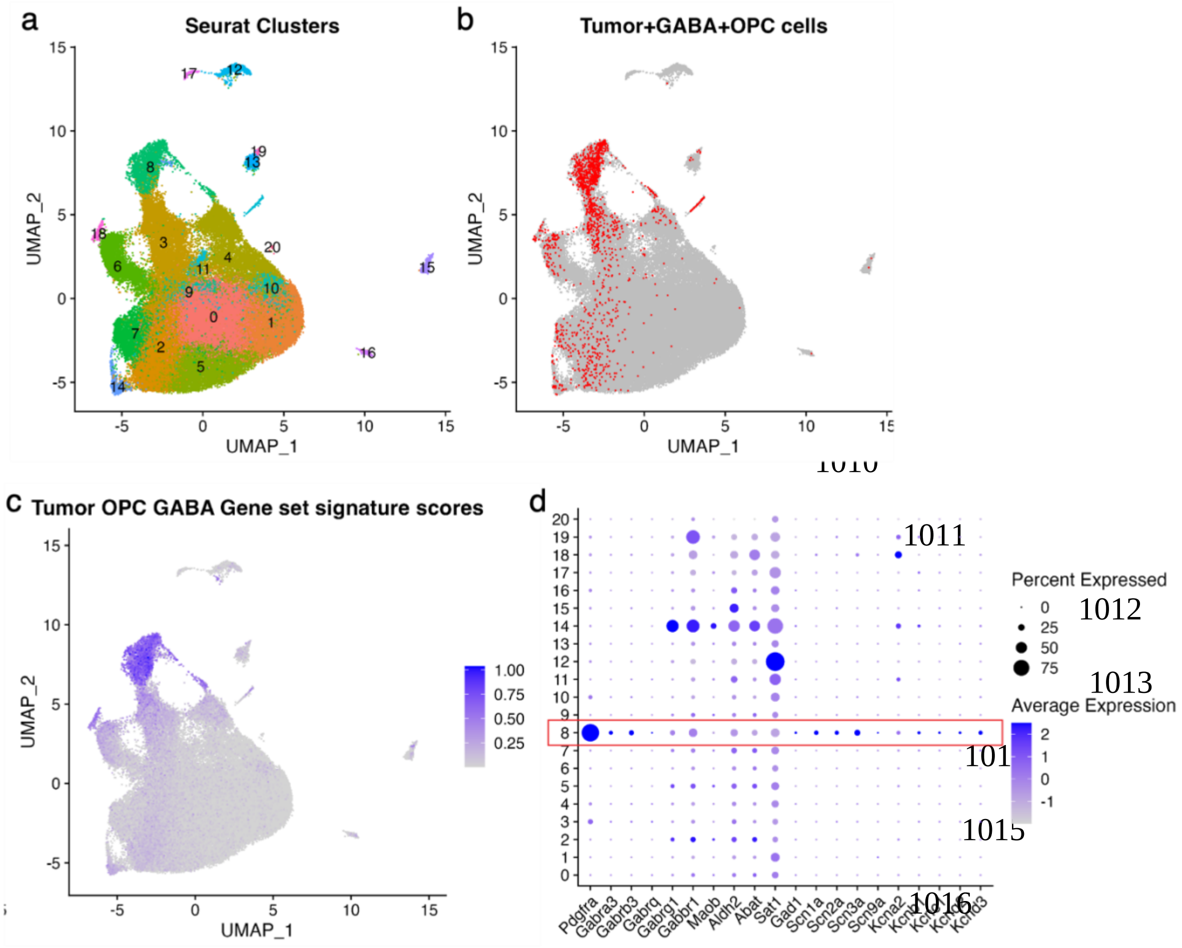

**Extended Data Figure S40. GABA-OPC tumor cells are detected in our scRNA-seq data of the IUE mouse model of glioma.** (a) Seurat clusters for scRNA-seq cells from n=3 IUE tumor mice. (b) Feature plot shows GABA-OPC tumor cells identified. (c) Feature plot showing cells with high tumor GABA-OPC geneset scores. (d) Dot plot showing that Seurat Cluster 8 contains GABA-OPC tumor cells that possess the GABA metabolism and voltage-gated ion channel genes found in human GABA-OPC tumor and non-tumor cells.

**Extended Data Figure S41**

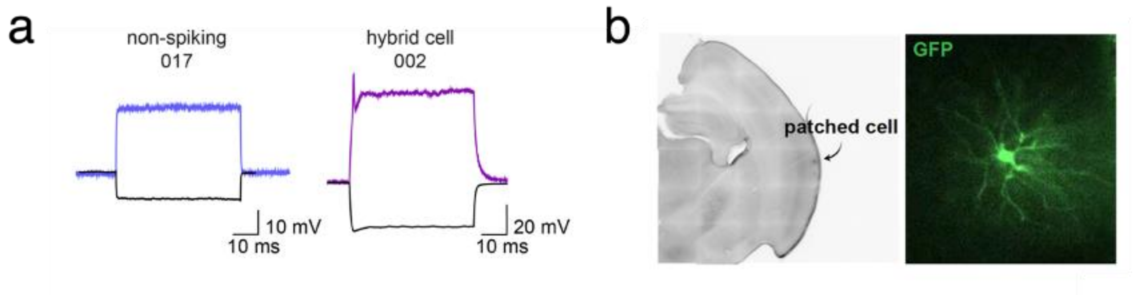

**Extended Data Figure S41. Hybrid Cells are detected in our IUE mouse model of glioma.** (a) Whole cell electrophysiology recordings from patch clamp experiments in our IUE model of *de novo* GFP-labeled glioma tumor cells confirm non-spiking (glia) and spiking (hybrid cell; HC) profiles. Spiking HCs in mouse glioma show analogous properties to those found in human glioma and are capable of firing single, short action potentials. (b) Immunohistochemistry of a mouse glioma cortical slice used for patch clamp experiments showing GFP reactivity and a corresponding immunofluorescence image showing GFP-labeling of the patched tumor cell.

#### Extended Data Figure S42

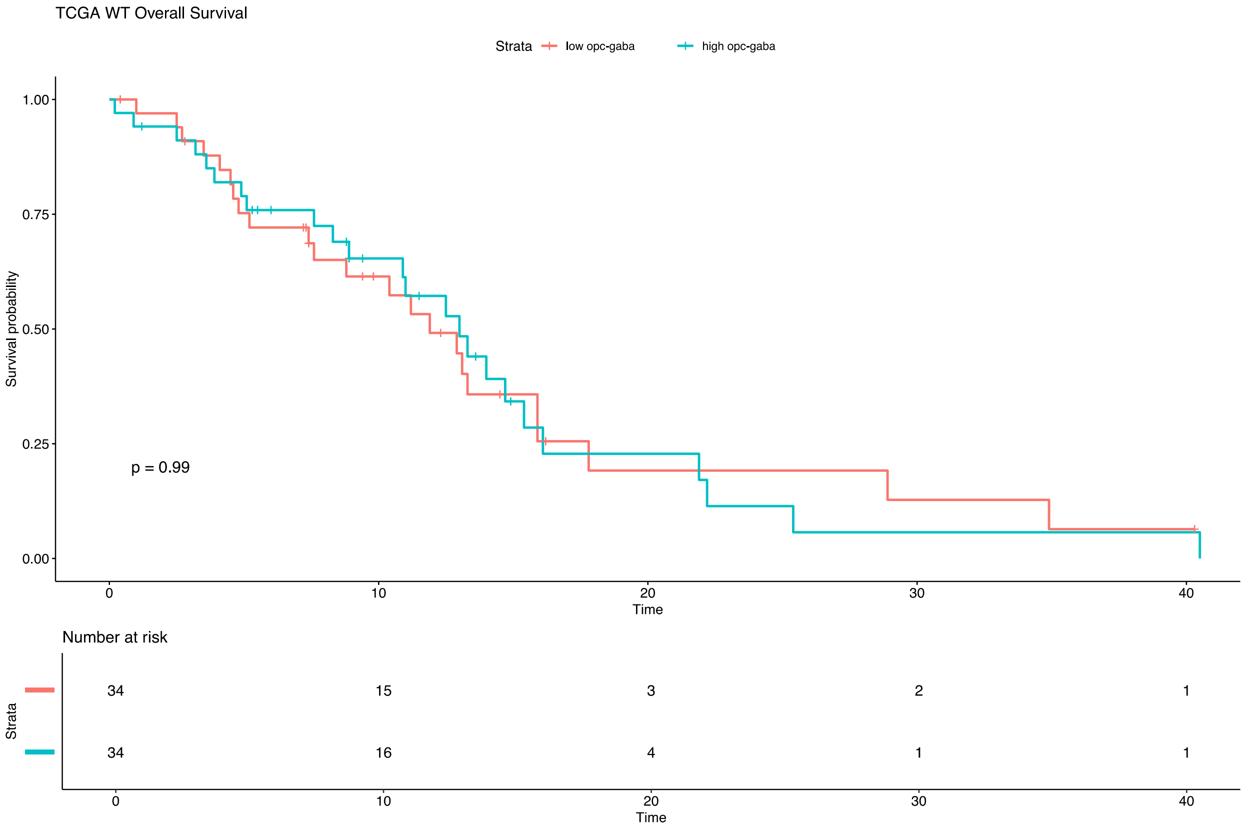

**Extended Data Figure S42. Survival Analysis in IDH WT TCGA Samples.** Survival analysis comparing IDH WT TCGA cohort samples with high GABA-OPC scores versus low GABA-OPC scores based on upper and lower quartile scoring.

**Extended Data Figure S43**

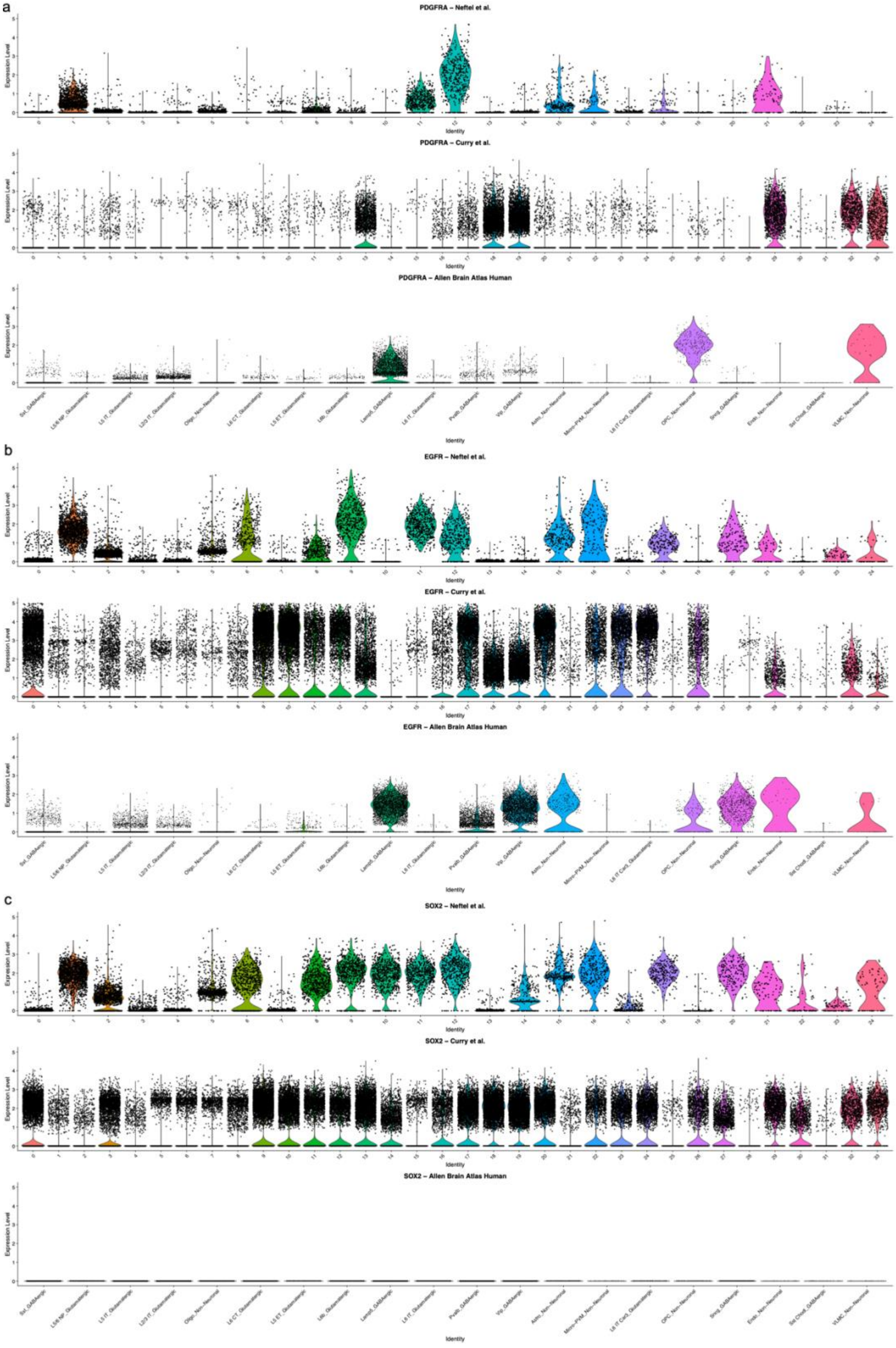

**Extended Data Figure S43 Tumor Marker Expression in Various Tumor and Non-Tumor Single-Cell RNA-Sequencing Datasets.** Tumour marker expression models used to generate expression thresholds are shown for (a) *PDGFRA* ($PDGFRA^{hi} \geq 3.63$) (b) *EGFR* ($EGFR^{hi} \geq 3.77$), and (c) *SOX2* ($SOX2^{hi} \geq 2.18$).

**Table S1**. Characteristics of whole-cell patch clamp recorded patient samples included in this study.

| Patient ID | Tumor | Pathology | Grade | Recurrence | Age | HC | GL | IN | PC | Experiments |
|---|---|---|---|---|---|---|---|---|---|---|
| PS-A1 | IDH mutant | astrocytoma | III | primary | 56 | 23 | 6 | 4 | 9 | Patch-seq |
| PS-A2 | IDH mutant | astrocytoma | II | primary | 26 | 3 | 2 | 1 | 5 | electrophysiology |
| PS-A3 | IDH mutant | astrocytoma | IV | primary | 37 | 0 | 0 | 5 | 6 | Patch-seq |
| PS-A4 | IDH mutant | astrocytoma | III | primary | 78 | 0 | 9 | 0 | 0 | Patch-seq |
| PS-G1 | IDH wildtype | astrocytoma | IV | primary | 70 | 0 | 3 | 0 | 0 | electrophysiology |
| PS-G2 | IDH wildtype | GBM | IV | primary | 44 | 0 | 2 | 3 | 12 | Patch-seq |
| PS-N1 | non-tumor | hypercellular glial mass <1% KI67 positivity and <5% BRAFV600E | NA | primary | 27 | 3 | 2 | 2 | 3 | electrophysiology |
| PS-O1 | IDH mutant | oligodendroglioma | II | primary | 59 | 8 | 1 | 4 | 5 | electrophysiology |
| PS-O2 | IDH mutant | oligodendroglioma | II | recurrent | 57 | 6 | 2 | 5 | 14 | Patch-seq |

Abbreviations: Patch-seq, whole-cell patch clamp recordings, followed by singe-cell RNA-sequencing; WHO, World Health Organization.

**Table S2.** Summary of Patch-seq cells used for this study. For *IDH1R132H* status, ✓ denotes heterozygous *IDH1R132H* mutation and ✓✓ denotes homozygous *IDH1R132H* mutation.

| Cell ID | Cell Type | Morphology | Sequencing | IDH1R132H | CNV | Patient ID | Subtype | Recurrence | Pathology |
|---|---|---|---|---|---|---|---|---|---|
| R1 | HC | ✓ | ✓ | | | PS-A1 | IDH mutant | primary | astrocytoma |
| R2 | GL | | ✓ | | | PS-A1 | IDH mutant | primary | astrocytoma |
| R3 | GL | | ✓ | | | PS-A1 | IDH mutant | primary | astrocytoma |
| R4 | PC | ✓ | ✓ | | | PS-A1 | IDH mutant | primary | astrocytoma |
| R5 | PC | | ✓ | | | PS-A1 | IDH mutant | primary | astrocytoma |
| R6 | HC | | ✓ | | | PS-A1 | IDH mutant | primary | astrocytoma |
| R7 | HC | | ✓ | | | PS-A1 | IDH mutant | primary | astrocytoma |
| R8 | HC | ✓ | ✓ | | | PS-A1 | IDH mutant | primary | astrocytoma |
| R9 | IN | | ✓ | | | PS-A1 | IDH mutant | primary | astrocytoma |
| R10 | HC | | ✓ | | | PS-A1 | IDH mutant | primary | astrocytoma |
| R11 | IN | | ✓ | | | PS-A1 | IDH mutant | primary | astrocytoma |
| R12 | IN | | ✓ | | | PS-A1 | IDH mutant | primary | astrocytoma |
| R13 | HC | | ✓ | ✓ | | PS-A1 | IDH mutant | primary | astrocytoma |
| R14 | HC | | ✓ | ✓ | | PS-A1 | IDH mutant | primary | astrocytoma |
| R15 | HC | | ✓ | ✓ | | PS-A1 | IDH mutant | primary | astrocytoma |
| R16 | IN | ✓ | ✓ | | | PS-A1 | IDH mutant | primary | astrocytoma |
| R17 | PC | | ✓ | | | PS-A1 | IDH mutant | primary | astrocytoma |
| R18 | HC | | ✓ | | | PS-A1 | IDH mutant | primary | astrocytoma |
| R19 | PC | | ✓ | | | PS-A1 | IDH mutant | primary | astrocytoma |
| R20 | PC | | ✓ | | | PS-A1 | IDH mutant | primary | astrocytoma |
| R21 | HC | | ✓ | | | PS-A1 | IDH mutant | primary | astrocytoma |
| R22 | HC | ✓ | ✓ | ✓ | | PS-A1 | IDH mutant | primary | astrocytoma |
| R23 | HC | ✓ | ✓ | ✓ | | PS-A1 | IDH mutant | primary | astrocytoma |
| R24 | HC | ▪ | ✓ | | | PS-A1 | IDH mutant | primary | astrocytoma |
| R25 | HC | ▪ | ✓ | | | PS-A1 | IDH mutant | primary | astrocytoma |
| R26 | PC | | ✓ | | | PS-A1 | IDH mutant | primary | astrocytoma |
| R27 | PC | | ✓ | | | PS-A1 | IDH mutant | primary | astrocytoma |
| R28 | PC | | ✓ | | | PS-A1 | IDH mutant | primary | astrocytoma |
| R29 | PC | | ✓ | | | PS-A1 | IDH mutant | primary | astrocytoma |
| R30 | GL | | ✓ | | | PS-A1 | IDH mutant | primary | astrocytoma |
| R31 | HC | | ✓ | no | | PS-A1 | IDH mutant | primary | astrocytoma |
| R32 | GL | ✓ | ▪ | | | PS-A2 | IDH mutant | primary | astrocytoma |

| | | | | | | | | | |
|---|---|---|---|---|---|---|---|---|---|
| R33 | HC | ✓ | | | | PS-A2 | IDH mutant | primary | astrocytoma |
| R34 | HC | | | | | PS-A2 | IDH mutant | primary | astrocytoma |
| R35 | PC | ✓ | | | | PS-A2 | IDH mutant | primary | astrocytoma |
| R36 | IN | ✓ | | | | PS-A2 | IDH mutant | primary | astrocytoma |
| R37 | PC | ✓ | | | | PS-A2 | IDH mutant | primary | astrocytoma |
| R38 | PC | ✓ | | | | PS-A2 | IDH mutant | primary | astrocytoma |
| R39 | PC | ✓ | | | | PS-A2 | IDH mutant | primary | astrocytoma |
| R40 | HC | ✓ | | | | PS-A2 | IDH mutant | primary | astrocytoma |
| R41 | PC | ✓ | | | | PS-A2 | IDH mutant | primary | astrocytoma |
| R42 | GL | ✓ | | | | PS-A2 | IDH mutant | primary | astrocytoma |
| R43 | HC | | | | | PS-A1 | IDH mutant | primary | astrocytoma |
| R44 | GL | ✓ | | | | PS-A1 | IDH mutant | primary | astrocytoma |
| R45 | HC | ✓ | | | | PS-A1 | IDH mutant | primary | astrocytoma |
| R46 | HC | | | | | PS-A1 | IDH mutant | primary | astrocytoma |
| R47 | HC | | | | | PS-A1 | IDH mutant | primary | astrocytoma |
| R48 | HC | ✓ | | | | PS-A1 | IDH mutant | primary | astrocytoma |
| R49 | GL | | | | | PS-A1 | IDH mutant | primary | astrocytoma |
| R50 | HC | | | | | PS-A1 | IDH mutant | primary | astrocytoma |
| R51 | HC | | | | | PS-A1 | IDH mutant | primary | astrocytoma |
| R52 | GL | | | | | PS-A1 | IDH mutant | primary | astrocytoma |
| R53 | HC | ✓ | | | | PS-A1 | IDH mutant | primary | astrocytoma |
| R54 | GL | | | | | PS-G1 | IDH wild-type | primary | astrocytoma |
| R55 | GL | | | | | PS-G1 | IDH wild-type | primary | astrocytoma |
| R56 | GL | | | | | PS-G1 | IDH wild-type | primary | astrocytoma |
| R57 | GL | | ✓ | | | PS-N1 | non-tumor | primary | hypercellular glial tissue |
| R58 | HC | | ✓ | | | PS-N1 | non-tumor | primary | hypercellular glial tissue |
| R59 | GL | | ✓ | | | PS-N1 | non-tumor | primary | hypercellular glial tissue |
| R60 | HC | | ✓ | | | PS-N1 | non-tumor | primary | hypercellular glial tissue |
| R61 | IN | | ✓ | | | PS-N1 | non-tumor | primary | hypercellular glial tissue |
| R62 | PC | | ✓ | | | PS-N1 | non-tumor | primary | hypercellular glial tissue |
| R63 | PC | | ✓ | | | PS-N1 | non-tumor | primary | hypercellular glial tissue |
| R64 | PC | | ✓ | | | PS-N1 | non-tumor | primary | hypercellular glial tissue |
| R65 | HC | | ✓ | | | PS-N1 | non-tumor | primary | hypercellular glial tissue |
| R66 | IN | | ✓ | | | PS-N1 | non-tumor | primary | hypercellular glial tissue |
| R67 | HC | | ✓ | | | PS-O1 | IDH mutant | primary | oligodendroglioma |
| R68 | GL | | ✓ | | | PS-O1 | IDH mutant | primary | oligodendroglioma |
| R69 | PC | | ✓ | | | PS-O1 | IDH mutant | primary | oligodendroglioma |
| R70 | PC | | ✓ | | | PS-O1 | IDH mutant | primary | oligodendroglioma |

| | | | | | | | | | |
|---|---|---|---|---|---|---|---|---|---|
| R71 | IN | | ✓ | | | PS-O1 | IDH mutant | primary | oligodendroglioma |
| R72 | IN | | ✓ | | | PS-O1 | IDH mutant | primary | oligodendroglioma |
| R73 | PC | | ✓ | | | PS-O1 | IDH mutant | primary | oligodendroglioma |
| R74 | PC | | ✓ | | | PS-O1 | IDH mutant | primary | oligodendroglioma |
| R75 | PC | | ▪ | | | PS-O1 | IDH mutant | primary | oligodendroglioma |
| R76 | IN | | ✓ | | | PS-O1 | IDH mutant | primary | oligodendroglioma |
| R77 | HC | | ✓ | | | PS-O1 | IDH mutant | primary | oligodendroglioma |
| R78 | HC | | ✓ | | | PS-O1 | IDH mutant | primary | oligodendroglioma |
| R79 | HC | | ✓ | | | PS-O1 | IDH mutant | primary | oligodendroglioma |
| R80 | HC | | ✓ | | | PS-O1 | IDH mutant | primary | oligodendroglioma |
| R81 | IN | | ✓ | | | PS-O1 | IDH mutant | primary | oligodendroglioma |
| R82 | HC | | ✓ | | | PS-O1 | IDH mutant | primary | oligodendroglioma |
| R83 | HC | | ✓ | | | PS-O1 | IDH mutant | primary | oligodendroglioma |
| R84 | HC | | ✓ | | | PS-O1 | IDH mutant | primary | oligodendroglioma |
| R85 | PC | | ✓ | | 19q deletion | PS-A3 | IDH mutant | primary | astrocytoma |
| R86 | PC | | ✓ | | | PS-A3 | IDH mutant | primary | astrocytoma |
| R87 | PC | | ✓ | | | PS-A3 | IDH mutant | primary | astrocytoma |
| R88 | IN | | ✓ | | | PS-A3 | IDH mutant | primary | astrocytoma |
| R89 | IN | | ✓ | | | PS-A3 | IDH mutant | primary | astrocytoma |
| R90 | IN | | ✓ | | | PS-A3 | IDH mutant | primary | astrocytoma |
| R91 | IN | | ✓ | | | PS-A3 | IDH mutant | primary | astrocytoma |
| R92 | IN | ✓ | ✓ | | | PS-A3 | IDH mutant | primary | astrocytoma |
| R93 | PC | | ✓ | | | PS-A3 | IDH mutant | primary | astrocytoma |
| R94 | PC | ✓ | ✓ | | | PS-A3 | IDH mutant | primary | astrocytoma |
| R95 | PC | ✓ | ✓ | | | PS-A3 | IDH mutant | primary | astrocytoma |
| R96 | IN | ✓ | ✓ | | | PS-G2 | IDH wild-type | primary | GBM |
| R97 | PC | ✓ | ✓ | | | PS-G2 | IDH wild-type | primary | GBM |
| R98 | IN | | ✓ | | | PS-G2 | IDH wild-type | primary | GBM |
| R99 | PC | | ✓ | | | PS-G2 | IDH wild-type | primary | GBM |
| R100 | GL | | ✓ | | 7p amplification | PS-G2 | IDH wild-type | primary | GBM |
| R101 | PC | | ✓ | | | PS-G2 | IDH wild-type | primary | GBM |
| R102 | PC | | ✓ | | | PS-G2 | IDH wild-type | primary | GBM |
| R103 | PC | | ✓ | | | PS-G2 | IDH wild-type | primary | GBM |
| R104 | PC | ✓ | ✓ | | | PS-G2 | IDH wild-type | primary | GBM |
| R105 | PC | ✓ | ✓ | | | PS-G2 | IDH wild-type | primary | GBM |
| R106 | PC | | ✓ | | | PS-G2 | IDH wild-type | primary | GBM |
| R107 | GL | | ✓ | | 7p amplification | PS-G2 | IDH wild-type | primary | GBM |

| | | | | | | | | | |
|---|---|---|---|---|---|---|---|---|---|
| R108 | IN | ✓ | ✓ | | | PS-G2 | IDH wild-type | primary | GBM |
| R109 | PC | | ✓ | | | PS-G2 | IDH wild-type | primary | GBM |
| R110 | PC | | ✓ | | | PS-G2 | IDH wild-type | primary | GBM |
| R111 | PC | | ✓ | | | PS-G2 | IDH wild-type | primary | GBM |
| R112 | PC | | ✓ | | | PS-G2 | IDH wild-type | primary | GBM |
| R113 | PC | | ✓ | | | PS-O2 | IDH mutant | recurrent | oligodendroglioma |
| R114 | PC | | ✓ | | 1p deletion | PS-O2 | IDH mutant | recurrent | oligodendroglioma |
| R115 | PC | | ✓ | | | PS-O2 | IDH mutant | recurrent | oligodendroglioma |
| R116 | PC | | ✓ | ✓ ✓ | | PS-O2 | IDH mutant | recurrent | oligodendroglioma |
| R117 | HC | | ✓ | ✓ | 1p deletion | PS-O2 | IDH mutant | recurrent | oligodendroglioma |
| R118 | PC | | ✓ | ✓ ✓ | | PS-O2 | IDH mutant | recurrent | oligodendroglioma |
| R119 | HC | | ✓ | | | PS-O2 | IDH mutant | recurrent | oligodendroglioma |
| R120 | PC | | ✓ | | 1p deletion | PS-O2 | IDH mutant | recurrent | oligodendroglioma |
| R121 | IN | | ✓ | | | PS-O2 | IDH mutant | recurrent | oligodendroglioma |
| R122 | HC | | ✓ | no | | PS-O2 | IDH mutant | recurrent | oligodendroglioma |
| R123 | PC | | ✓ | | | PS-O2 | IDH mutant | recurrent | oligodendroglioma |
| R124 | GL | | ✓ | | | PS-O2 | IDH mutant | recurrent | oligodendroglioma |
| R125 | GL | | ✓ | | | PS-O2 | IDH mutant | recurrent | oligodendroglioma |
| R126 | PC | | ✓ | | | PS-O2 | IDH mutant | recurrent | oligodendroglioma |
| R127 | PC | | ✓ | | | PS-O2 | IDH mutant | recurrent | oligodendroglioma |
| R128 | HC | | ✓ | | | PS-O2 | IDH mutant | recurrent | oligodendroglioma |
| R129 | PC | | ✓ | | | PS-O2 | IDH mutant | recurrent | oligodendroglioma |
| R130 | PC | | ✓ | X | 1p deletion | PS-O2 | IDH mutant | recurrent | oligodendroglioma |
| R131 | IN | | ✓ | | | PS-O2 | IDH mutant | recurrent | oligodendroglioma |
| R132 | PC | | ✓ | | | PS-O2 | IDH mutant | recurrent | oligodendroglioma |
| R133 | IN | | ✓ | | | PS-O2 | IDH mutant | recurrent | oligodendroglioma |
| R134 | IN | | ✓ | | | PS-O2 | IDH mutant | recurrent | oligodendroglioma |
| R135 | PC | | ✓ | X | | PS-O2 | IDH mutant | recurrent | oligodendroglioma |
| R136 | HC | | ✓ | | 1p deletion | PS-O2 | IDH mutant | recurrent | oligodendroglioma |
| R137 | IN | | ✓ | | | PS-O2 | IDH mutant | recurrent | oligodendroglioma |
| R138 | PC | | ✓ | ✓ ✓ | | PS-O2 | IDH mutant | recurrent | oligodendroglioma |
| R139 | HC | | ✓ | | | PS-O2 | IDH mutant | recurrent | oligodendroglioma |
| R140 | GL | | ✓ | | | PS-A4 | IDH mutant | primary | astrocytoma |
| R141 | GL | | ✓ | | | PS-A4 | IDH mutant | primary | astrocytoma |
| R142 | GL | | ✓ | ✓ | | PS-A4 | IDH mutant | primary | astrocytoma |
| R143 | GL | | ✓ | | | PS-A4 | IDH mutant | primary | astrocytoma |
| R144 | GL | | ✓ | | | PS-A4 | IDH mutant | primary | astrocytoma |
| R145 | GL | | ✓ | | 7p amplification | PS-A4 | IDH mutant | primary | astrocytoma |
| R146 | GL | | ✓ | | | PS-A4 | IDH mutant | primary | astrocytoma |

| | | | | | | | | | |
|---|---|---|---|---|---|---|---|---|---|
| R147 | GL | | ✓ | | 7p amplification | PS-A4 | IDH mutant | primary | astrocytoma |
| R148 | GL | | ✓ | | | PS-A4 | IDH mutant | primary | astrocytoma |

**Table S3.** Summary of Patch-seq cells mapping quality.

| Cell ID | nCount_RNA | nFeature_RNA | Electrophysiology | IDH1R132H |
|---|---|---|---|---|
| R31 | 1311512 | 3040 | HC | no |
| R122 | 11024537 | 5286 | HC | no |
| R13 | 992716 | 6585 | HC | yes |
| R14 | 1108817 | 3104 | HC | yes |
| R15 | 1146113 | 4400 | HC | yes |
| R22 | 918071 | 4378 | HC | yes |
| R23 | 33798 | 2000 | HC | yes |
| R117 | 12884695 | 6037 | HC | yes |
| R10 | 1145337 | 3043 | HC | NA |
| R18 | 932416 | 2790 | HC | NA |
| R01 | 717406 | 3737 | HC | NA |
| R21 | 634160 | 2835 | HC | NA |
| R24 | 875008 | 2944 | HC | NA |
| R25 | 1130843 | 3222 | HC | NA |
| R06 | 887796 | 3116 | HC | NA |
| R07 | 950548 | 3519 | HC | NA |
| R08 | 937441 | 2622 | HC | NA |
| R119 | 8671686 | 3133 | HC | NA |
| R128 | 12358045 | 3834 | HC | NA |
| R136 | 8685235 | 4067 | HC | NA |
| R139 | 15476517 | 4444 | HC | NA |
| R144 | 8818984 | 4118 | GL | no |
| R142 | 9792491 | 3162 | GL | yes-Homozygous |
| R02 | 917267 | 3052 | GL | NA |
| R30 | 924450 | 3296 | GL | NA |
| R03 | 342341 | 3270 | GL | NA |
| R100 | 3575432 | 3486 | GL | NA |
| R107 | 1607476 | 3242 | GL | NA |
| R124 | 18733378 | 4324 | GL | NA |
| R125 | 15329110 | 4666 | GL | NA |
| R140 | 7110350 | 3413 | GL | NA |
| R141 | 11504613 | 3077 | GL | NA |
| R143 | 14007976 | 3275 | GL | NA |
| R145 | 12047789 | 3573 | GL | NA |
| R146 | 12231412 | 3136 | GL | NA |
| R147 | 13331102 | 3779 | GL | NA |

| | | | | |
|---|---|---|---|---|
| R148 | 10746842 | 2917 | GL | NA |
| R11 | 554372 | 5032 | IN | NA |
| R12 | 654303 | 5701 | IN | NA |
| R16 | 1071436 | 5441 | IN | NA |
| R09 | 886334 | 4934 | IN | NA |
| R108 | 2362845 | 4005 | IN | NA |
| R88 | 2744506 | 8052 | IN | NA |
| R89 | 3458751 | 8103 | IN | NA |
| R90 | 2450864 | 3729 | IN | NA |
| R91 | 2616041 | 6523 | IN | NA |
| R92 | 3309424 | 7339 | IN | NA |
| R96 | 403410 | 9388 | IN | NA |
| R98 | 5379305 | 6691 | IN | NA |
| R121 | 6852804 | 5619 | IN | NA |
| R131 | 13536663 | 4808 | IN | NA |
| R133 | 4933751 | 4623 | IN | NA |
| R134 | 5362094 | 3557 | IN | NA |
| R137 | 6881135 | 5607 | IN | NA |
| R130 | 12610728 | 4340 | PC | no |
| R118 | 16384828 | 7119 | PC | yes |
| R116 | 8100008 | 4035 | PC | yes-Homozygous |
| R138 | 12494633 | 3650 | PC | yes-Homozygous |
| R17 | 339606 | 5859 | PC | NA |
| R19 | 744841 | 5594 | PC | NA |
| R20 | 567694 | 5393 | PC | NA |
| R26 | 937697 | 3852 | PC | NA |
| R27 | 32579 | 2754 | PC | NA |
| R28 | 257812 | 2485 | PC | NA |
| R29 | 426433 | 2470 | PC | NA |
| R04 | 472149 | 6012 | PC | NA |
| R05 | 390775 | 4430 | PC | NA |
| R101 | 1541534 | 3166 | PC | NA |
| R102 | 2521389 | 5884 | PC | NA |
| R103 | 2137060 | 3713 | PC | NA |
| R104 | 2196949 | 5583 | PC | NA |
| R105 | 3643594 | 6351 | PC | NA |
| R106 | 2629007 | 5563 | PC | NA |
| R109 | 2097984 | 3993 | PC | NA |
| R110 | 2157596 | 5741 | PC | NA |
| R111 | 2183293 | 4198 | PC | NA |
| R112 | 3144677 | 6023 | PC | NA |

| | | | | |
|---|---|---|---|---|
| R85 | 1465652 | 5568 | PC | NA |
| R86 | 1764337 | 5108 | PC | NA |
| R87 | 2687307 | 7262 | PC | NA |
| R93 | 1659230 | 5096 | PC | NA |
| R94 | 2246992 | 3067 | PC | NA |
| R95 | 1676040 | 5138 | PC | NA |
| R97 | 2883603 | 6315 | PC | NA |
| R99 | 2852483 | 4396 | PC | NA |
| R113 | 7575821 | 5010 | PC | NA |
| R114 | 5098854 | 4181 | PC | NA |
| R115 | 10483311 | 4283 | PC | NA |
| R120 | 12304741 | 4472 | PC | NA |
| R123 | 9321767 | 4583 | PC | NA |
| R126 | 12063244 | 3681 | PC | NA |
| R127 | 8479414 | 3733 | PC | NA |
| R129 | 10011213 | 5148 | PC | NA |
| R132 | 6916015 | 4381 | PC | NA |
| R135 | 8804751 | 3964 | PC | NA |

**Table S4.** Summary of training datasets

| Dataset | num of cells | Single Cell Technology | Type |
|---|---|---|---|
| Bhaduri et al. | 404455 | 10X Genomics | Developing Human Brain |
| CODEX | 33739 | Drop-seq | Developing Human Brain |
| Aldinger et al. | 69174 | SPLiT-seq | Developing Human Cerebellar Brain |
| Glioma Immune Atlas | 21303 | 10X Genomics | Glioma Immune Atlas |
| Venteicher/Tirosh et al. | 9537 | SMART-Seq2 | IDH Mutant Oligodendroglioma and Astrocytoma Glioma |
| Richards et al | 87541 | 10X Genomics | IDH WT and GSC Glioma |
| Neftel et al. | 3429 | SMART-Seq2 | IDH WT Glioma |
| Allen Brain Atlas | 76533 | 10X Genomics | Normal Brain Atlas |
| HPA Brain Atlas | 76533 | 10X Genomics | Normal Brain Atlas |
| Tissue Immune Atlas | 329762 | 10X Genomics | Normal Immune Atlas |

**Table S5.** Cell Types in training datasets

| Cell Type | Cell Type Simplified | Cell Class | Dataset |
|---|---|---|---|
| H-Ast_aldinger | astrocyte | neural | Aldinger et al. |
| H-Ast/Ependymal_aldinger | astrocyte | neural | Aldinger et al. |
| H-BG_aldinger | other | other | Aldinger et al. |
| H-Brainstem_aldinger | other | other | Aldinger et al. |
| H-BS Choroid/Ependymal_aldinger | other | other | Aldinger et al. |
| H-Choroid_aldinger | other | other | Aldinger et al. |
| H-Committed OPC_aldinger | oligodendrocyte precursor | neural | Aldinger et al. |
| H-eCN/UBC_aldinger | other | other | Aldinger et al. |
| H-Endothelial_aldinger | endothelial | vascular | Aldinger et al. |
| H-GCP_aldinger | other | other | Aldinger et al. |
| H-Glia_aldinger | other | other | Aldinger et al. |
| H-GN_aldinger | other | other | Aldinger et al. |
| H-iCN_aldinger | other | other | Aldinger et al. |
| H-Meninges_aldinger | other | other | Aldinger et al. |
| H-Microglia_aldinger | immune | immune | Aldinger et al. |
| H-MLI_aldinger | other | other | Aldinger et al. |
| H-OPC_aldinger | oligodendrocyte precursor | neural | Aldinger et al. |
| H-PC_aldinger | other | other | Aldinger et al. |
| H-Pericytes_aldinger | other | other | Aldinger et al. |
| H-PIP_aldinger | other | other | Aldinger et al. |
| H-RL_aldinger | other | other | Aldinger et al. |
| Astro | astrocyte | neural | Allen Brain Atlas |
| GABAergic | GABAergic | neural | Allen Brain Atlas |
| Glutamatergic | Glutamatergic | neural | Allen Brain Atlas |
| Micro-PVM | immune | immune | Allen Brain Atlas |
| Oligo | oligodendrocytes | neural | Allen Brain Atlas |
| OPC | oligodendrocyte precursor | neural | Allen Brain Atlas |
| Vascular | vascular | vascular | Allen Brain Atlas |
| GABAergic_Lamp5_allen | GABAergic | neural | Allen Brain Atlas |
| GABAergic_Pvalb_allen | GABAergic | neural | Allen Brain Atlas |
| GABAergic_Sncg_allen | GABAergic | neural | Allen Brain Atlas |
| GABAergic_Sst Chodl_allen | GABAergic | neural | Allen Brain Atlas |
| GABAergic_Sst_allen | GABAergic | neural | Allen Brain Atlas |
| GABAergic_Vip_allen | GABAergic | neural | Allen Brain Atlas |

| | | | |
|---|---|---|---|
| Glutamatergic_L2/3 IT_allen | Glutamatergic | neural | Allen Brain Atlas |
| Glutamatergic_L5 ET_allen | Glutamatergic | neural | Allen Brain Atlas |
| Glutamatergic_L5 IT_allen | Glutamatergic | neural | Allen Brain Atlas |
| Glutamatergic_L5/6 NP_allen | Glutamatergic | neural | Allen Brain Atlas |
| Glutamatergic_L6 CT_allen | Glutamatergic | neural | Allen Brain Atlas |
| Glutamatergic_L6 IT Car3_allen | Glutamatergic | neural | Allen Brain Atlas |
| Glutamatergic_L6 IT_allen | Glutamatergic | neural | Allen Brain Atlas |
| Glutamatergic_L6b_allen | Glutamatergic | neural | Allen Brain Atlas |
| CR_Middle_bhaduri | other | other | Bhaduri et al. |
| Dividing_Early_bhaduri | cycling progenitor cell | tumor | Bhaduri et al. |
| Dividing_Late_bhaduri | cycling progenitor cell | tumor | Bhaduri et al. |
| Dividing_Middle_bhaduri | cycling progenitor cell | tumor | Bhaduri et al. |
| Interneuron_Late_bhaduri | Interneuron | neural | Bhaduri et al. |
| Interneuron_Middle_bhaduri | Interneuron | neural | Bhaduri et al. |
| IPC_Early_bhaduri | other | other | Bhaduri et al. |
| IPC_Middle_bhaduri | other | other | Bhaduri et al. |
| Microglia_Late_bhaduri | immune | immune | Bhaduri et al. |
| Microglia_Middle_bhaduri | immune | immune | Bhaduri et al. |
| Neuron_Early_bhaduri | neuron | neural | Bhaduri et al. |
| Neuron_Late_bhaduri | neuron | neural | Bhaduri et al. |
| Neuron_Middle_bhaduri | neuron | neural | Bhaduri et al. |
| OPC_Late_bhaduri | oligodendrocyte precursor | neural | Bhaduri et al. |
| OPC_Middle_bhaduri | oligodendrocyte precursor | neural | Bhaduri et al. |
| Outlier_Early_bhaduri | other | other | Bhaduri et al. |
| Outlier_Late_bhaduri | other | other | Bhaduri et al. |
| Outlier_Middle_bhaduri | other | other | Bhaduri et al. |
| RG_Early_bhaduri | radial glia | neural | Bhaduri et al. |
| RG_Late_bhaduri | radial glia | neural | Bhaduri et al. |
| RG_Middle_bhaduri | radial glia | neural | Bhaduri et al. |
| Vascular_Early_bhaduri | vascular | vascular | Bhaduri et al. |
| Vascular_Late_bhaduri | vascular | vascular | Bhaduri et al. |
| Vascular_Middle_bhaduri | vascular | vascular | Bhaduri et al. |
| End_codex | other | other | CoDEx |
| ExDp1_codex | other | other | CoDEx |
| ExDp2_codex | other | other | CoDEx |
| ExM_codex | other | other | CoDEx |
| ExM-U_codex | other | other | CoDEx |

| | | | |
|---|---|---|---|
| ExN_codex | other | other | CoDEx |
| InCGE_codex | other | other | CoDEx |
| InMGE_codex | other | other | CoDEx |
| IP_codex | other | other | CoDEx |
| Mic_codex | other | other | CoDEx |
| OPC_codex | oligodendrocyte precursor | neural | CoDEx |
| oRG_codex | radial glia | neural | CoDEx |
| Per_codex | other | other | CoDEx |
| PgG2M_codex | cycling progenitor cell | tumor | CoDEx |
| PgS_codex | cycling progenitor cell | tumor | CoDEx |
| vRG_codex | radial glia | neural | CoDEx |
| B cells_primary_gbm | immune | immune | Glioma Immune Atlas and Richards et al. |
| DC_primary_gbm | immune | immune | Glioma Immune Atlas and Richards et al. |
| Developmental_GSC | Developmental_GSC | Developmental_GSC | Glioma Immune Atlas and Richards et al. |
| Immune | immune | immune | Glioma Immune Atlas and Richards et al. |
| Injury_GSC | Injury_GSC | Injury_GSC | Glioma Immune Atlas and Richards et al. |
| Monocytes_primary_gbm | immune | immune | Glioma Immune Atlas and Richards et al. |
| NK cells_primary_gbm | immune | immune | Glioma Immune Atlas and Richards et al. |
| NormalBrain | oligodendrocytes | neural | Glioma Immune Atlas and Richards et al. |
| prol. TAM_primary_gbm | immune | immune | Glioma Immune Atlas and Richards et al. |
| T cells_primary_gbm | immune | immune | Glioma Immune Atlas and Richards et al. |

| | | | |
|---|---|---|---|
| TAM 1_primary_gbm | immune | immune | Glioma Immune Atlas and Richards et al. |
| TAM 2_primary_gbm | immune | immune | Glioma Immune Atlas and Richards et al. |
| Tumour | tumor | tumor | Glioma Immune Atlas and Richards et al. |
| Astrocytes_hpa_brain | astrocyte | neural | HPA Brain Atlas |
| Excitatory_neurons_hpa_brain | excitatory | neural | HPA Brain Atlas |
| Inhibitory_neurons_hpa_brain | Inhibitory | neural | HPA Brain Atlas |
| Microglial_cells_hpa_brain | immune | immune | HPA Brain Atlas |
| Oligodendrocyte_precursor_cells_hpa_brain | oligodendrocyte precursor | neural | HPA Brain Atlas |
| Oligodendrocytes_hpa_brain | oligodendrocytes | neural | HPA Brain Atlas |
| immune_suva | immune | immune | Venteicher et al., Tirosh et al. |
| malignant_suva | tumor | tumor | Venteicher et al., Tirosh et al. |
| oligodendrocytes_suva | oligodendrocytes | neural | Venteicher et al., Tirosh et al. |
| AClike_suva | AClike_suva | AClike_suva | Neftel et al. |
| MESlike_suva | MESlike_suva | MESlike_suva | Neftel et al. |
| NPClike_suva | NPClike_suva | NPClike_suva | Neftel et al. |
| OPClike_suva | OPClike_suva | OPClike_suva | Neftel et al. |
| ABCs_tissueImmune | immune | immune | Tissue Immune Atlas |
| Alveolar macrophages_tissueImmune | immune | immune | Tissue Immune Atlas |
| Classical monocytes_tissueImmune | immune | immune | Tissue Immune Atlas |
| Cycling T&NK_tissueImmune | cycling progenitor cell | tumor | Tissue Immune Atlas |
| Cycling_tissueImmune | cycling progenitor cell | tumor | Tissue Immune Atlas |
| DC1_tissueImmune | immune | immune | Tissue Immune Atlas |
| DC2_tissueImmune | immune | immune | Tissue Immune Atlas |
| Erythroid_tissueImmune | immune | immune | Tissue Immune Atlas |
| Erythrophagocytic macrophages_tissueImmune | immune | immune | Tissue Immune Atlas |
| GC_B (I)_tissueImmune | immune | immune | Tissue Immune Atlas |

| | | | |
|---|---|---|---|
| ILC3_tissueImmune | immune | immune | Tissue Immune Atlas |
| Intermediate macrophages_tissueImmune | immune | immune | Tissue Immune Atlas |
| Intestinal macrophages_tissueImmune | immune | immune | Tissue Immune Atlas |
| MAIT_tissueImmune | immune | immune | Tissue Immune Atlas |
| Mast cells_tissueImmune | immune | immune | Tissue Immune Atlas |
| Megakaryocytes_tissueImmune | immune | immune | Tissue Immune Atlas |
| Memory B cells_tissueImmune | immune | immune | Tissue Immune Atlas |
| migDC_tissueImmune | immune | immune | Tissue Immune Atlas |
| MNP/B doublets_tissueImmune | immune | immune | Tissue Immune Atlas |
| MNP/T doublets_tissueImmune | immune | immune | Tissue Immune Atlas |
| Naive B cells_tissueImmune | immune | immune | Tissue Immune Atlas |
| NK_CD16+_tissueImmune | immune | immune | Tissue Immune Atlas |
| NK_CD56bright_CD16-_tissueImmune | immune | immune | Tissue Immune Atlas |
| Nonclassical monocytes_tissueImmune | immune | immune | Tissue Immune Atlas |
| pDC_tissueImmune | immune | immune | Tissue Immune Atlas |
| Plasma cells_tissueImmune | immune | immune | Tissue Immune Atlas |
| Plasmablasts_tissueImmune | immune | immune | Tissue Immune Atlas |
| Pre-B_tissueImmune | immune | immune | Tissue Immune Atlas |
| Progenitor_tissueImmune | immune | immune | Tissue Immune Atlas |
| T_CD4/CD8_tissueImmune | immune | immune | Tissue Immune Atlas |
| T/B doublets_tissueImmune | immune | immune | Tissue Immune Atlas |
| Teffector/EM_CD4_tissueImmune | immune | immune | Tissue Immune Atlas |
| Tem/emra_CD8_tissueImmune | immune | immune | Tissue Immune Atlas |

| | | | |
|---|---|---|---|
| Tfh_tissueImmune | immune | immune | Tissue Immune Atlas |
| Tgd_CRTAM+_tissueImmune | immune | immune | Tissue Immune Atlas |
| Tnaive/CM_CD4_activated_tissueImmune | immune | immune | Tissue Immune Atlas |
| Tnaive/CM_CD4_tissueImmune | immune | immune | Tissue Immune Atlas |
| Tnaive/CM_CD8_tissueImmune | immune | immune | Tissue Immune Atlas |
| Tregs_tissueImmune | immune | immune | Tissue Immune Atlas |
| Trm_gut_CD8_tissueImmune | immune | immune | Tissue Immune Atlas |
| Trm_Tgd_tissueImmune | immune | immune | Tissue Immune Atlas |
| Trm_Th1/Th17_tissueImmune | immune | immune | Tissue Immune Atlas |
| Trm/em_CD8_tissueImmune | immune | immune | Tissue Immune Atlas |

**Table S6**. Number of tumor cells detected by SCRAM in developing and healthy brain datasets

| | #tumor cells | #total cells |
|---|---|---|
| ABA | 559 | 76533 |
| HPA-Brain | 1333 | 76533 |
| Bhaduri et al | 4237 | 404218 |
| Aldinger et al | 586 | 69174 |
| CoDEx | 559 | 75974 |

**Table S7**. Characteristics of scRNA-seq patient samples included in this study

| **Patient Identity** | **Tumor Type** | ***IDH1* Status** | **1p19 Codeletion** | **WHO Grade** | **Samples** | **Experiments** |
|---|---|---|---|---|---|---|
| DA-01 | diffuse astrocytoma | mutant | no | IV | core<br>leading edge | scRNA-seq |
| GBM-01 | GBM | wild-type | no | IV | core<br>leading edge | scRNA-seq |
| GBM-02 | GBM | wild-type | no | IV | core<br>leading edge | scRNA-seq |
| OG-01 | oligodendroglioma | mutant | yes | II | core | scRNA-seq |
| GBM-03 | GBM | wild-type | no | IV | core<br>leading edge | scRNA-seq |
| GBM-04 | GBM | wild-type | no | IV | core | scRNA-seq |
| DA-02 | diffuse astrocytoma | mutant | no | IV | core<br>leading edge | scRNA-seq |
| OG-02 | oligodendroglioma | mutant | yes | II | core | scRNA-seq |
| OG-03 | oligodendroglioma | mutant | yes | III | core | scRNA-seq |
| GBM-05 | GBM | wild-type | no | IV | leading edge | scRNA-seq |
| GBM-06 | GBM | wild-type | no | IV | core | scRNA-seq |
| GBM-07 | GBM | wild-type | no | IV | core | scRNA-seq |

Abbreviations: GBM, glioblastoma;; scRNA-seq, single-cell RNA-sequencing; WHO, World Health Organization.

**Table S8.** GABAergic OPC DEGs.

| log2FC | pct.1 | pct.2 | p_val_adj | gene | SHAP_features |
|---|---|---|---|---|---|
| 2.69 | 0.66 | 0.057 | 0 | DLL3 | NPClike_suva,OPC_Late_bhaduri, |
| 2.27 | 0.828 | 0.158 | 0 | HES6 | NPClike_suva,OPC_Middle_bhaduri, |
| 2.11 | 0.853 | 0.191 | 0 | VCAN | OPClike_suva,OPC_codex,OPC_Middle_bhaduri, |
| 2.11 | 0.845 | 0.187 | 0 | SOX4 | NPClike_suva,OPC_Late_bhaduri, |
| 2.05 | 0.844 | 0.211 | 0 | OLIG1 | Oligodendrocyte_precursor_cells_hpa_brain,OPC_codex,OPC_Middle_bhaduri,OPC_Late_bhaduri,allenOPC, |
| 2.05 | 0.767 | 0.088 | 0 | C1QL1 | OPC_Middle_bhaduri, |
| 1.97 | 0.746 | 0.119 | 0 | FXYD6 | InCGE_codex,OPC_Late_bhaduri, |
| 1.92 | 0.799 | 0.134 | 0 | MAP2 | NPClike_suva, |
| 1.88 | 0.681 | 0.072 | 0 | SOX8 | aldinger_neuralNetwork_H-OPC_aldinger, |
| 1.87 | 0.737 | 0.113 | 0 | GRIA2 | OPC_codex,OPC_Middle_bhaduri, |
| 1.86 | 0.627 | 0.051 | 0 | FERMT1 | allenOPC, |
| 1.86 | 0.737 | 0.091 | 0 | OLIG2 | OPC_codex,OPC_Middle_bhaduri,OPC_Late_bhaduri, |
| 1.77 | 0.706 | 0.083 | 0 | SOX6 | InMGE_codex,OPC_Middle_bhaduri,allen_GABAergic_Sst Chodl_allen,allen_GABAergic_Pvalb_allen,aldinger_neuralNetwork_H-OPC_aldinger, |
| 1.75 | 0.895 | 0.317 | 0 | MARCKSL1 | allenGABAergic, |
| 1.72 | 0.649 | 0.08 | 0 | TNR | OPClike_suva,Oligodendrocyte_precursor_cells_hpa_brain,allenOPC,aldinger_neuralNetwork_H-OPC_aldinger, |
| 1.72 | 0.765 | 0.117 | 0 | NCAM1 | dirks_primary_gbm_Developmental_GSC, |
| 1.71 | 0.825 | 0.193 | 0 | SCD5 | dirks_primary_gbm_Tumour,allenOPC, |
| 1.68 | 0.588 | 0.054 | 0 | PDGFRA | Oligodendrocyte_precursor_cells_hpa_brain,dirks_primary_gbm_Developmental_GSC,OPC_codex,OPC_Middle_bhaduri,OPC_Late_bhaduri,aldinger_neuralNetwork_H-OPC_aldinger, |
| 1.65 | 0.487 | 0.073 | 0 | XIST | OPC_Late_bhaduri, |
| 1.64 | 0.518 | 0.043 | 0 | SMOC1 | OPClike_suva,suva_idh_a_o_malignant,OPC_Late_bhaduri,allenOPC,aldinger_neuralNetwork_H-Committed OPC_aldinger, |
| 1.61 | 0.647 | 0.066 | 0 | CTTNBP2 | allen_GABAergic_Lamp5_allen, |
| 1.60 | 0.735 | 0.139 | 0 | CADM2 | dirks_primary_gbm_Tumour,aldinger_neuralNetwork_H-OPC_aldinger, |
| 1.59 | 0.591 | 0.046 | 0 | NEU4 | OPClike_suva, |
| 1.56 | 0.719 | 0.152 | 0 | TCF12 | NPClike_suva,OPC_Late_bhaduri, |
| 1.55 | 0.685 | 0.159 | 0 | NRXN1 | allen_GABAergic_Sncg_allen, |
| 1.55 | 0.62 | 0.072 | 0 | NKAIN4 | OPClike_suva,OPC_codex,OPC_Late_bhaduri, |
| 1.53 | 0.641 | 0.071 | 0 | ALCAM | OPClike_suva,allen_GABAergic_Vip_allen,allen_GABAergic_Lamp5_allen, |
| 1.51 | 0.669 | 0.102 | 0 | C11orf96 | InMGE_codex, |
| 1.46 | 0.532 | 0.038 | 0 | SHD | NPClike_suva,OPC_codex, |
| 1.44 | 0.533 | 0.047 | 0 | FGF12 | Oligodendrocyte_precursor_cells_hpa_brain, |
| 1.41 | 0.784 | 0.19 | 0 | NFIB | Excitatory_neurons_hpa_brain,InCGE_codex,allen_GABAergic_Vip_allen,allen_GABAergic_Sst Chodl_allen, |
| 1.38 | 0.612 | 0.095 | 0 | RAB3IP | allen_GABAergic_Pvalb_allen, |
| 1.36 | 0.664 | 0.133 | 0 | GRIA4 | allen_GABAergic_Vip_allen,allen_GABAergic_Pvalb_allen, |

| | | | | | |
|---|---|---|---|---|---|
| 1.34 | 0.797 | 0.213 | 0 | NOVA1 | dirks_primary_gbm_Tumour,OPC_codex,OPC_Middle_bhaduri, |
| 1.33 | 0.539 | 0.066 | 0 | SNTG1 | Oligodendrocyte_precursor_cells_hpa_brain,aldinger_neuralNetwork_H-OPC_aldinger, |
| 1.31 | 0.509 | 0.173 | 0 | HSPA1A | Oligodendrocytes_hpa_brain,dirks_primary_gbm_Tumour, |
| 1.30 | 0.753 | 0.199 | 0 | ETV1 | OPC_codex,OPC_Late_bhaduri, |
| 1.29 | 0.689 | 0.145 | 0 | LIMA1 | OPC_Late_bhaduri, |
| 1.28 | 0.472 | 0.034 | 0 | PLPPR1 | Oligodendrocyte_precursor_cells_hpa_brain,OPC_Late_bhaduri, |
| 1.24 | 0.931 | 0.41 | 0 | ZBTB20 | allen_GABAergic_Vip_allen, |
| 1.23 | 0.738 | 0.175 | 0 | PGRMC1 | OPClike_suva, |
| 1.20 | 0.63 | 0.118 | 0 | JMJD1C | OPC_codex, |
| 1.18 | 0.499 | 0.054 | 0 | ARPP21 | Oligodendrocyte_precursor_cells_hpa_brain,allen_GABAergic_Sncg_allen,allenOPC, |
| 1.18 | 0.55 | 0.105 | 0 | SLC44A1 | OPClike_suva,aldinger_neuralNetwork_H-Committed OPC_aldinger, |
| 1.17 | 0.375 | 0.039 | 0 | SEZ6L | NPClike_suva,Oligodendrocyte_precursor_cells_hpa_brain,OPC_Late_bhaduri,allen_GABAergic_Vip_allen, |
| 1.15 | 0.942 | 0.45 | 0 | BCAN | OPC_codex,OPC_Middle_bhaduri,OPC_Late_bhaduri, |
| 1.14 | 0.482 | 0.055 | 0 | GALNT13 | allen_GABAergic_Vip_allen, |
| 1.12 | 0.518 | 0.062 | 0 | CHD7 | NPClike_suva,OPC_codex, |
| 1.12 | 0.479 | 0.055 | 0 | OPHN1 | allenOPC, |
| 1.11 | 0.634 | 0.137 | 0 | FAM181B | OPC_Middle_bhaduri, |
| 1.10 | 0.457 | 0.042 | 0 | GRID2 | allenOPC, |
| 1.09 | 0.453 | 0.037 | 0 | LHFPL3 | OPClike_suva,Oligodendrocyte_precursor_cells_hpa_brain,allenOPC,aldinger_neuralNetwork_H-OPC_aldinger, |
| 1.09 | 0.871 | 0.33 | 0 | TCF4 | allenGABAergic, |
| 1.06 | 0.43 | 0.055 | 0 | MMP16 | Oligodendrocyte_precursor_cells_hpa_brain,allen_GABAergic_Sncg_allen,allenOPC,aldinger_neuralNetwork_H-OPC_aldinger, |
| 1.04 | 0.623 | 0.125 | 0 | DNER | allen_GABAergic_Pvalb_allen, |
| 1.04 | 0.407 | 0.03 | 0 | UGT8 | OPC_codex,aldinger_neuralNetwork_H-Committed OPC_aldinger, |
| 1.03 | 0.695 | 0.176 | 0 | LSAMP | OPC_Middle_bhaduri,allenGABAergic, |
| 1.02 | 0.404 | 0.031 | 0 | GRIK2 | Inhibitory_neurons_hpa_brain,allen_GABAergic_Sncg_allen,allenOPC,allenGABAergic, |
| 1.02 | 0.615 | 0.133 | 0 | FYN | OPC_Late_bhaduri,allenOligo, |
| 1.01 | 0.576 | 0.254 | 0 | EGR1 | InCGE_codex, |
| 1.01 | 0.36 | 0.022 | 0 | PCDH15 | Oligodendrocyte_precursor_cells_hpa_brain,allen_GABAergic_Pvalb_allen, allen_GABAergic_Lamp5_allen, allenOPC,aldinger_neuralNetwork_H-OPC_aldinger, |
| 0.99 | 0.392 | 0.028 | 0 | TMEFF2 | allenOligo, |
| 0.98 | 0.521 | 0.093 | 0 | CCSER2 | suva_idh_a_o_malignant, |
| 0.98 | 0.425 | 0.041 | 0 | DNM3 | aldinger_neuralNetwork_H-Committed OPC_aldinger, |
| 0.98 | 0.75 | 0.274 | 0 | NRCAM | allenOPC, |
| 0.97 | 0.91 | 0.447 | 0 | JUND | OPC_Middle_bhaduri, |
| 0.96 | 0.394 | 0.039 | 0 | DCX | NPClike_suva, |
| 0.96 | 0.891 | 0.419 | 0 | QKI | OPC_Middle_bhaduri, |
| 0.95 | 0.569 | 0.127 | 0 | GABPB1-AS1 | suva_idh_a_o_malignant, |

| | | | | | |
|---|---|---|---|---|---|
| 0.95 | 0.424 | 0.08 | 0 | SIRT2 | dirks_primary_gbm_Developmental_GSC,OPC_codex,OPC_Middle_bhaduri ,aldinger_neuralNetwork_H-Committed OPC_aldinger, |
| 0.94 | 0.584 | 0.136 | 0 | DSEL | suva_idh_a_o_malignant, |
| 0.94 | 0.352 | 0.021 | 0 | NXPH1 | Oligodendrocyte_precursor_cells_hpa_brain,Inhibitory_neurons_hpa_brain, InMGE_codex,allen_GABAergic_Sst Chodl_allen,allen_GABAergic_Pvalb_allen,allenOPC,allenGABAergic, |
| 0.93 | 0.377 | 0.031 | 0 | TAGLN3 | NPClike_suva, |
| 0.92 | 0.382 | 0.047 | 0 | ROBO1 | Inhibitory_neurons_hpa_brain,allen_GABAergic_Vip_allen,allenGABAergic, |
| 0.91 | 0.361 | 0.044 | 0 | LRRC4C | Oligodendrocyte_precursor_cells_hpa_brain,allen_GABAergic_Pvalb_allen, aldinger_neuralNetwork_H-OPC_aldinger, |
| 0.88 | 0.525 | 0.108 | 0 | ASCL1 | NPClike_suva, |
| 0.88 | 0.481 | 0.075 | 0 | MLLT11 | NPClike_suva, |
| 0.87 | 0.468 | 0.083 | 0 | ZNF462 | aldinger_neuralNetwork_H-OPC_aldinger, |
| 0.87 | 0.817 | 0.328 | 0 | STMN1 | NPClike_suva,PgS_codex, |
| 0.87 | 0.381 | 0.052 | 0 | STMN4 | dirks_primary_gbm_Tumour, |
| 0.87 | 0.374 | 0.049 | 0 | SLAIN1 | OPC_Late_bhaduri, |
| 0.86 | 0.463 | 0.068 | 0 | THY1 | OPClike_suva, |
| 0.85 | 0.778 | 0.455 | 0 | JUN | dirks_primary_gbm_Tumour,InCGE_codex, |
| 0.85 | 0.378 | 0.042 | 0 | OMG | OPClike_suva, |
| 0.84 | 0.62 | 0.169 | 0 | H1FX | PgS_codex, |
| 0.83 | 0.324 | 0.027 | 0 | CALCRL | allen_GABAergic_Pvalb_allen, |
| 0.83 | 0.774 | 0.272 | 0 | BEX1 | NPClike_suva, |
| 0.81 | 0.323 | 0.031 | 0 | DLGAP1 | allenGABAergic, |
| 0.81 | 0.359 | 0.048 | 0 | LRRN1 | OPC_codex,allenOPC, |
| 0.79 | 0.325 | 0.026 | 0 | DSCAM | allen_GABAergic_Sncg_allen, |
| 0.78 | 0.282 | 0.022 | 0 | ELAVL4 | NPClike_suva, |
| 0.78 | 0.462 | 0.086 | 0 | SLC38A 2 | PgS_codex, |
| 0.77 | 0.398 | 0.066 | 0 | SOX11 | InMGE_codex, |
| 0.77 | 0.543 | 0.13 | 0 | KIDINS2 20 | InCGE_codex, |
| 0.77 | 0.418 | 0.072 | 0 | NLGN1 | aldinger_neuralNetwork_H-OPC_aldinger, |
| 0.76 | 0.257 | 0.025 | 0 | SCN1A | Oligodendrocyte_precursor_cells_hpa_brain,OPC_codex,OPC_Middle_bhad uri, |
| 0.76 | 0.278 | 0.021 | 0 | CSMD3 | allenOPC,aldinger_neuralNetwork_H-OPC_aldinger, |
| 0.74 | 0.339 | 0.042 | 0 | SATB1 | OPC_Late_bhaduri, |
| 0.73 | 0.361 | 0.068 | 0 | LUZP2 | allen_GABAergic_Sncg_allen,allen_GABAergic_Lamp5_allen,aldinger_neur alNetwork_H-OPC_aldinger, |
| 0.72 | 0.626 | 0.175 | 0 | RBM25 | InCGE_codex, |
| 0.72 | 0.317 | 0.03 | 0 | GNAI1 | aldinger_neuralNetwork_H-Committed OPC_aldinger, |
| 0.71 | 0.341 | 0.042 | 0 | NIN | NPClike_suva,Oligodendrocytes_hpa_brain,PgS_codex, |
| 0.71 | 0.31 | 0.031 | 0 | FGF14 | Oligodendrocyte_precursor_cells_hpa_brain, |
| 0.70 | 0.304 | 0.033 | 0 | CACNG4 | allenOPC, |
| 0.69 | 0.638 | 0.213 | 0 | ZEB1 | OPC_Late_bhaduri, |
| 0.68 | 0.746 | 0.264 | 0 | DDX17 | allenOligo, |

| | | | | | |
|---|---|---|---|---|---|
| 0.67 | 0.205 | 0.013 | 0 | GPR17 | OPClike_suva,aldinger_neuralNetwork_H-Committed OPC_aldinger, |
| 0.66 | 0.59 | 0.18 | 0 | PCDH17 | allen_GABAergic_Pvalb_allen,allen_GABAergic_Lamp5_allen, |
| 0.66 | 0.24 | 0.016 | 0 | DLL1 | NPClike_suva,OPC_Middle_bhaduri, |
| 0.65 | 0.279 | 0.029 | 0 | NCALD | allen_GABAergic_Pvalb_allen,aldinger_neuralNetwork_H-OPC_aldinger, |
| 0.65 | 0.434 | 0.094 | 0 | BCHE | OPC_Middle_bhaduri, |
| 0.64 | 0.227 | 0.014 | 0 | SPHKAP | allen_GABAergic_Lamp5_allen, |
| 0.64 | 0.266 | 0.042 | 0 | CNTN1 | OPClike_suva,aldinger_neuralNetwork_H-OPC_aldinger, |
| 0.63 | 0.27 | 0.031 | 0 | KCND2 | OPC_codex,aldinger_neuralNetwork_H-OPC_aldinger, |
| 0.63 | 0.461 | 0.144 | 0 | EPN2 | Oligodendrocyte_precursor_cells_hpa_brain, |
| 0.63 | 0.248 | 0.034 | 0 | PTPRD | allen_GABAergic_Sncg_allen, |
| 0.62 | 0.228 | 0.016 | 0 | MYT1 | OPClike_suva,OPC_Middle_bhaduri, |
| 0.62 | 0.333 | 0.069 | 0 | SYNE1 | allenGABAergic, |
| 0.62 | 0.35 | 0.083 | 0 | CNP | OPC_Late_bhaduri, |
| 0.62 | 0.269 | 0.032 | 0 | PAK3 | Inhibitory_neurons_hpa_brain, |
| 0.61 | 0.254 | 0.029 | 0 | TOX3 | Inhibitory_neurons_hpa_brain,InCGE_codex, |
| 0.61 | 0.185 | 0.032 | 0 | PLLP | OPClike_suva,OPC_Late_bhaduri, |
| 0.60 | 0.301 | 0.049 | 0 | LRP1B | aldinger_neuralNetwork_H-OPC_aldinger, |
| 0.60 | 0.264 | 0.026 | 0 | MEX3A | NPClike_suva, |
| 0.60 | 0.42 | 0.095 | 0 | PRKCA | InCGE_codex, |
| 0.60 | 0.253 | 0.036 | 0 | SULF2 | aldinger_neuralNetwork_H-OPC_aldinger, |
| 0.59 | 0.297 | 0.034 | 0 | GNG4 | Inhibitory_neurons_hpa_brain, |
| 0.58 | 0.71 | 0.258 | 0 | TSPAN3 | InMGE_codex, |
| 0.58 | 0.253 | 0.031 | 0 | MEIS2 | Excitatory_neurons_hpa_brain,InCGE_codex, |
| 0.58 | 0.209 | 0.015 | 0 | CA10 | OPClike_suva,Oligodendrocyte_precursor_cells_hpa_brain,OPC_codex,allen_GABAergic_Pvalb_allen,allenOPC, |
| 0.58 | 0.249 | 0.061 | 0 | SEMA5A | allen_GABAergic_Sncg_allen,aldinger_neuralNetwork_H-Committed OPC_aldinger, |
| 0.57 | 0.227 | 0.027 | 0 | CACNA2D1 | allen_GABAergic_Lamp5_allen, |
| 0.57 | 0.303 | 0.037 | 0 | DOCK10 | allen_GABAergic_Sst Chodl_allen, |
| 0.57 | 0.185 | 0.015 | 0 | ERBB3 | OPClike_suva,aldinger_neuralNetwork_H-Committed OPC_aldinger, |
| 0.57 | 0.324 | 0.06 | 0 | ZNF704 | allen_GABAergic_Lamp5_allen, |
| 0.56 | 0.44 | 0.112 | 0 | EEA1 | InCGE_codex, |
| 0.56 | 0.293 | 0.049 | 0 | PLCB1 | allenOligo, |
| 0.56 | 0.21 | 0.016 | 0 | SNX22 | OPC_Middle_bhaduri,allenOPC, |
| 0.56 | 0.264 | 0.03 | 0 | RTKN | OPClike_suva, |
| 0.56 | 0.294 | 0.04 | 0 | MARCH1 | allenOPC, |
| 0.55 | 0.17 | 0.02 | 0 | EDIL3 | Oligodendrocytes_hpa_brain,allen_GABAergic_Lamp5_allen,allenOligo, |
| 0.55 | 0.212 | 0.02 | 0 | GABRB3 | Excitatory_neurons_hpa_brain, |
| 0.55 | 0.344 | 0.07 | 0 | MEST | dirks_primary_gbm_Tumour, |
| 0.54 | 0.239 | 0.03 | 0 | BEST3 | allenOPC, |

| | | | | | |
|---|---|---|---|---|---|
| 0.54 | 0.402 | 0.086 | 0 | MNAT1 | Excitatory_neurons_hpa_brain, |
| 0.53 | 0.26 | 0.04 | 0 | APLP1 | suva_idh_a_o_oligodendrocytes_suva, |
| 0.52 | 0.206 | 0.013 | 0 | VSTM2B | allen_GABAergic_Lamp5_allen, |
| 0.52 | 0.202 | 0.022 | 0 | COL11A1 | Oligodendrocyte_precursor_cells_hpa_brain, |
| 0.51 | 0.217 | 0.018 | 0 | NKX2-2 | aldinger_neuralNetwork_H-Committed OPC_aldinger, |
| 0.51 | 0.293 | 0.049 | 0 | SIK3 | Oligodendrocytes_hpa_brain, |
| 0.50 | 0.135 | 0.031 | 0 | BCAS1 | aldinger_neuralNetwork_H-Committed OPC_aldinger, |
| 0.50 | 0.369 | 0.086 | 0 | MIA3 | allenOligo, |
| 0.50 | 0.251 | 0.04 | 0 | PEG10 | PgS_codex, |
| 0.48 | 0.189 | 0.018 | 0 | TMEM132B | allen_GABAergic_Pvalb_allen, |
| 0.48 | 0.357 | 0.108 | 0 | AMER2 | suva_idh_a_o_malignant, |
| 0.47 | 0.355 | 0.094 | 0 | SRGAP3 | allen_GABAergic_Vip_allen, |
| 0.47 | 0.235 | 0.038 | 0 | THSD7A | allen_GABAergic_Sst_allen,allenGABAergic, |
| 0.47 | 0.193 | 0.018 | 0 | ATP8A1 | aldinger_neuralNetwork_H-Committed OPC_aldinger, |
| 0.46 | 0.209 | 0.026 | 0 | UNC80 | OPClike_suva,NPClike_suva, |
| 0.46 | 0.328 | 0.076 | 0 | PHIP | InMGE_codex, |
| 0.46 | 0.309 | 0.066 | 0 | APBB2 | InMGE_codex, |
| 0.46 | 0.161 | 0.036 | 0 | SFRP1 | PgS_codex, |
| 0.45 | 0.265 | 0.059 | 0 | SLC6A1 | allen_GABAergic_Vip_allen,allen_GABAergic_Pvalb_allen, |
| 0.45 | 0.167 | 0.017 | 0 | MIAT | NPClike_suva, |
| 0.45 | 0.165 | 0.009 | 0 | ZNF804A | allenOligo,allenGABAergic, |
| 0.44 | 0.334 | 0.082 | 0 | VPS13C | InMGE_codex, |
| 0.44 | 0.164 | 0.009 | 0 | COL20A1 | OPClike_suva,OPC_codex,aldinger_neuralNetwork_H-OPC_aldinger, |
| 0.43 | 0.262 | 0.047 | 0 | CBX6 | allen_GABAergic_Pvalb_allen, |
| 0.43 | 0.205 | 0.034 | 0 | ADGRB3 | Excitatory_neurons_hpa_brain, |
| 0.43 | 0.437 | 0.127 | 0 | RNF13 | allen_GABAergic_Pvalb_allen, |
| 0.43 | 0.156 | 0.013 | 0 | RANBP17 | allen_GABAergic_Sst Chodl_allen,allenGABAergic, |
| 0.43 | 0.499 | 0.15 | 0 | SNRNP70 | NPClike_suva, |
| 0.41 | 0.224 | 0.039 | 0 | KLF12 | allenGABAergic, |
| 0.41 | 0.197 | 0.025 | 0 | TSHZ1 | allen_GABAergic_Pvalb_allen, |
| 0.41 | 0.184 | 0.032 | 0 | SLITRK2 | OPC_Late_bhaduri, |
| 0.40 | 0.689 | 0.28 | 0 | EIF4G2 | dirks_primary_gbm_Tumour, |
| 0.40 | 0.633 | 0.233 | 0 | NDUFC2 | dirks_primary_gbm_Tumour,dirks_primary_gbm_Developmental_GSC, |
| 0.40 | 0.821 | 0.404 | 0 | SET | dirks_primary_gbm_Tumour, |
| 0.40 | 0.517 | 0.162 | 0 | MORF4L2 | PgS_codex, |
| 0.39 | 0.129 | 0.01 | 0 | SOX10 | suva_idh_a_o_oligodendrocytes_suva, |
| 0.39 | 0.189 | 0.02 | 0 | UST | allenOPC, |
| 0.39 | 0.227 | 0.043 | 0 | CLASP1 | InCGE_codex, |

| | | | | | |
|---|---|---|---|---|---|
| 0.38 | 0.166 | 0.018 | 0 | ATRNL1 | Oligodendrocyte_precursor_cells_hpa_brain, |
| 0.38 | 0.768 | 0.375 | 0 | SCRG1 | OPC_Middle_bhaduri,OPC_Late_bhaduri, |
| 0.38 | 0.16 | 0.011 | 0 | MDGA2 | allenOPC,aldinger_neuralNetwork_H-Committed OPC_aldinger, |
| 0.37 | 0.202 | 0.03 | 0 | SLC1A1 | OPClike_suva, |
| 0.37 | 0.41 | 0.123 | 0 | PUM1 | NPClike_suva, |
| 0.37 | 0.174 | 0.02 | 0 | DGKB | allenGABAergic, |
| 0.36 | 0.132 | 0.008 | 0 | ELAVL2 | Inhibitory_neurons_hpa_brain,allenGABAergic, |
| 0.36 | 0.266 | 0.063 | 0 | KLHL42 | suva_idh_a_o_malignant, |
| 0.36 | 0.13 | 0.013 | 0 | CSMD1 | aldinger_neuralNetwork_H-OPC_aldinger, |
| 0.35 | 0.176 | 0.025 | 0 | GSE1 | allen_GABAergic_Vip_allen, |
| 0.34 | 0.441 | 0.144 | 0 | MT-ATP8 | dirks_primary_gbm_Developmental_GSC, |
| 0.34 | 0.167 | 0.023 | 0 | ZNF276 | OPC_codex, |
| 0.34 | 0.115 | 0.011 | 0 | EYA1 | allenOPC, |
| 0.34 | 0.246 | 0.06 | 0 | NRXN2 | InMGE_codex, |
| 0.34 | 0.449 | 0.16 | 0 | PDE4B | allen_GABAergic_Vip_allen,allen_GABAergic_Sst_allen,allen_GABAergic_Sst Chodl_allen, |
| 0.33 | 0.48 | 0.156 | 0 | SYNCRIP | InCGE_codex, |
| 0.33 | 0.108 | 0.009 | 0 | NPPA | NPClike_suva, |
| 0.33 | 0.146 | 0.022 | 0 | ERBB4 | Inhibitory_neurons_hpa_brain,allen_GABAergic_Sst Chodl_allen,allen_GABAergic_Lamp5_allen, |
| 0.33 | 0.214 | 0.066 | 0 | RASD1 | dirks_primary_gbm_Tumour, |
| 0.33 | 0.277 | 0.07 | 0 | RPAP2 | OPC_codex, |
| 0.33 | 0.207 | 0.043 | 0 | PTBP2 | allenOligo, |
| 0.32 | 0.23 | 0.058 | 0 | PCDH7 | aldinger_neuralNetwork_H-Committed OPC_aldinger, |
| 0.32 | 0.139 | 0.015 | 0 | GAD1 | Inhibitory_neurons_hpa_brain,allen_GABAergic_Sst Chodl_allen,allenGABAergic, |
| 0.32 | 0.256 | 0.084 | 0 | PPP1R15A | InCGE_codex, |
| 0.32 | 0.131 | 0.01 | 0 | AFAP1L2 | OPClike_suva, |
| 0.31 | 0.14 | 0.018 | 0 | PTPRK | allen_GABAergic_Pvalb_allen,aldinger_neuralNetwork_H-OPC_aldinger, |
| 0.31 | 0.119 | 0.013 | 0 | LRRTM4 | allen_GABAergic_Sncg_allen,allen_GABAergic_Pvalb_allen,allenOPC,aldinger_neuralNetwork_H-OPC_aldinger, |
| 0.31 | 0.172 | 0.039 | 0 | SLC35F1 | aldinger_neuralNetwork_H-OPC_aldinger, |
| 0.31 | 0.179 | 0.035 | 0 | PCDHGC3 | OPClike_suva, |
| 0.30 | 0.178 | 0.032 | 0 | HOOK2 | allenOligo, |
| 0.30 | 0.151 | 0.021 | 0 | KIAA1549 | Excitatory_neurons_hpa_brain, |
| 0.29 | 0.705 | 0.306 | 0 | MAP1B | allen_GABAergic_Vip_allen, |
| 0.29 | 0.133 | 0.016 | 0 | CBLB | OPC_Middle_bhaduri, |
| 0.29 | 0.181 | 0.041 | 0 | LANCL1 | allen_GABAergic_Pvalb_allen, |
| 0.29 | 0.114 | 0.01 | 0 | RAB3C | Inhibitory_neurons_hpa_brain,allenGABAergic, |
| 0.29 | 0.239 | 0.08 | 0 | SCD | suva_idh_a_o_oligodendrocytes_suva, |
| 0.29 | 0.185 | 0.035 | 0 | ACACA | allen_GABAergic_Sst Chodl_allen, |

| | | | | | |
|---|---|---|---|---|---|
| 0.29 | 0.114 | 0.024 | 0 | HAPLN1 | allen_GABAergic_Sncg_allen, |
| 0.28 | 0.281 | 0.083 | 0 | PSAT1 | OPClike_suva, |
| 0.28 | 0.241 | 0.074 | 0 | PROX1 | InCGE_codex,allen_GABAergic_Lamp5_allen, |
| 0.28 | 0.123 | 0.016 | 0 | RUNX1T1 | Inhibitory_neurons_hpa_brain,InMGE_codex,allen_GABAergic_Sst_allen,allenOPC,allenGABAergic, |
| 0.28 | 0.345 | 0.105 | 0 | MRPL32 | Excitatory_neurons_hpa_brain, |
| 0.28 | 0.17 | 0.032 | 0 | GRIK3 | allen_GABAergic_Sst Chodl_allen, |
| 0.28 | 0.109 | 0.016 | 0 | PLEKHH2 | Oligodendrocyte_precursor_cells_hpa_brain,OPC_codex,aldinger_neuralNetwork_H-OPC_aldinger, |
| 0.27 | 0.241 | 0.07 | 0 | FRYL | suva_idh_a_o_oligodendrocytes_suva, |
| 0.27 | 0.124 | 0.027 | 0 | SHC3 | OPClike_suva, |
| 0.27 | 0.138 | 0.025 | 0 | FGFBP3 | dirks_primary_gbm_Developmental_GSC, |
| 0.26 | 0.147 | 0.026 | 0 | LRRC49 | Excitatory_neurons_hpa_brain, |
| 0.26 | 0.219 | 0.057 | 0 | PCF11 | InCGE_codex, |
| 0.26 | 0.15 | 0.041 | 0 | CDH10 | Oligodendrocyte_precursor_cells_hpa_brain,allen_GABAergic_Vip_allen,aldinger_neuralNetwork_H-OPC_aldinger, |
| 0.26 | 0.149 | 0.034 | 0 | BRINP3 | aldinger_neuralNetwork_H-OPC_aldinger, |
| 0.26 | 0.11 | 0.018 | 0 | CELF4 | Inhibitory_neurons_hpa_brain,allen_GABAergic_Sncg_allen, |

#### References

1. Polioudakis, D. *et al.* A single-cell transcriptomic atlas of human neocortical development during mid-gestation. *Neuron* **103**, 785-801.e8 (2019).
2. Aldinger, K. A. *et al.* Spatial and cell type transcriptional landscape of human cerebellar development. *Nat. Neurosci.* **24**, 1163–1175 (2021).
3. Bhaduri, A. *et al.* An atlas of cortical arealization identifies dynamic molecular signatures. *Nature* **598**, 200–204 (2021).
4. Sjöstedt, E. *et al.* An atlas of the protein-coding genes in the human, pig, and mouse brain. *Science* **367**, (2020).
5. Bakken, T. E. *et al.* Comparative cellular analysis of motor cortex in human, marmoset and mouse. *Nature* **598**, 111–119 (2021).
6. Venteicher, A. S. *et al.* Decoupling genetics, lineages, and microenvironment in IDH-mutant gliomas by single-cell RNA-seq. *Science* **355**, (2017).
7. Tirosh, I. *et al.* Single-cell RNA-seq supports a developmental hierarchy in human oligodendroglioma. *Nature* **539**, 309–313 (2016).
8. Neftel, C. *et al.* An integrative model of cellular states, plasticity, and genetics for glioblastoma. *Cell* **178**, 835-849.e21 (2019).
9. Richards, L. M. *et al.* Gradient of Developmental and Injury Response transcriptional states defines functional vulnerabilities underpinning glioblastoma heterogeneity. *Nat Cancer* **2**, 157–173 (2021).
10. Pombo Antunes, A. R. *et al.* Single-cell profiling of myeloid cells in glioblastoma across species and disease stage reveals macrophage competition and specialization. *Nat. Neurosci.* **24**, 595–610 (2021).

11. Domínguez Conde, C. *et al.* Cross-tissue immune cell analysis reveals tissue-specific features in humans. *Science* **376**, eabl5197 (2022).

12. Marques, S. *et al.* Oligodendrocyte heterogeneity in the mouse juvenile and adult central nervous system. *Science* **352**, 1326–1329 (2016).

13. Marques, S. *et al.* Transcriptional convergence of oligodendrocyte lineage progenitors during development. *Dev. Cell* **46**, 504-517.e7 (2018).
